# Can growth in captivity alter the calcaneal microanatomy of a wild ungulate?

**DOI:** 10.1101/2022.08.22.504790

**Authors:** Romain Cottereau, Katia Ortiz, Yann Locatelli, Alexandra Houssaye, Thomas Cucchi

**Affiliations:** CNRS, UMR 7179 Mécanismes Adaptatifs et Evolution, Muséum d’Histoire Naturelle de Paris, France; Réserve Zoologique de la Haute-Touche, Muséum National d’Histoire Naturelle, Obterre, France; Institut de Systématique, Evolution, Biodiversité, UMR 7205, Muséum National d’Histoire Naturelle CNRS UPMC EPHE, UA, Paris, France; hysiologie de la Reproduction et des Comportements, UMR 7247, INRAE CNRS Université de Tours IFCE, Nouzilly, France; UMR 7209 Archéozoologie, Archéobotanique : Sociétés, Pratiques et Environnements, Muséum d’Histoire Naturelle de Paris, France

**Author notes:** Co last authors (Alexandra Houssaye, Thomas Cucchi). **Cite as**, Cottereau, R., Ortiz, K., Locatelli, Y., Houssaye, A., and Cucchi, T. (2022) Can growth in captivity alter the calcaneal microanatomy of a wild ungulate? BioRxiv, 504790, ver. 5 peer-reviewed and recommended by Peer Community in Archaeology. https://doi.orR/10.1101/2022.08.22.504790.

**Keywords:** calcaneus, bone structure, captivity, domestication, *Sus scrofa*, functional morphology

## Abstract

Reduced mobility associated with captivity induces changes in biomechanical stress on the skeleton of domesticated animals. Due to bone plasticity, bone’s morphology and internal structure can respond to these new biomechanical stresses over individuals’ lifetime. In a context where documenting early process of animal domestication is challenging, this study will test the hypothesis that change in mobility patterns during a wild ungulate’s life will alter the internal structure of its limb bones and provide a proof of concept for the application of this knowledge in Zooarchaeology. Using the calcaneus as a phenotypic marker through qualitative and quantitative 3D microanatomical analyses, we relied on a comparative study across wild boars (*Sus scrofa*) populations from controlled experimental conditions with different mobility patterns (natural habitat, large pen, and stall) and archaeological specimens collected from middle and late Mesolithic as surrogate for the norm of reaction in European wild boar phenotype before the spread of agriculture and domestic pigs. Results provide evidence for compressive and tensile forces as the main elements affecting the variation in the cortical thickness along the calcaneus. Furthermore, changes in the internal structure of the calcaneus between mobility patterns are observed but their intensity is not directly associated with the degree of mobility restriction and only weakly impacted by the size or weight of the individuals. Despite having greater bone volume, the calcaneus of the Mesolithic wild boars displays a very similar microanatomy compared to the present-day hunted or captive wild boars. These results suggest that calcaneal microanatomy is more affected by population differences than by locomotor variation. For all these reasons, this preliminary study doesn’t support the use of microanatomy of the calcaneus as an indicator of change in locomotor behaviour induced by captivity in the archaeological record.

## Introduction

The bones that make up the vertebrate skeleton are plastic organs whose morphology, both external and internal, adapt in response to physical stresses (Roux, 1881; Wolff, 1986; Hall, 1983, 2005; Ruff et al., 2006; Du et al., 2020), beyond the predominant influence of heredity (Hall, 1989; Cubo et al., 2005). Muscle mobilization applies stresses that influence bone growth, development, and remodeling (Marcus, 2002). Numerous studies on human skeletons have shown that intensive physical activity has characteristic consequences on skeletal microanatomy (Zanker & Swaine, 2000; Modlesky et al., 2008; Maïmoun & Sultan 2011; Maïmoun et al., 2013). Conversely, the bone resorption observed in astronauts (Lang et al., 2004) and bedridden individuals (Krølner & Toft., 1983) illustrates bone accommodation to the absence of gravity (Carmeliet & Bouillon, 2001) and inactivity. Therefore, differences in mobility between individuals, which engenders differences in biomechanical stresses, affect the bones’ structure.

Captive animals may grow in limited areas, which does not involve the same range of movement and forces applied to their bones compared to free-ranging individuals. Analysing the biomechanical bone plasticity is a major asset for paleoanthropologists and archaeozoologists trying to decipher individual-scale lifestyle changes from bones anatomy in order to document changes in activity patterns of past humans and other animals (Trinkaus et al., 1994, Agarwal, 2016). In paleoanthropology, this biomechanical component has been used to better understand the evolution of bipedalism in hominids (Ruff, 2018) and to observe the morphological consequences of the transition from a nomadic hunter-gatherer to a farmer-herder lifestyle (Pinhasi & Stock, 2011). In archaeozoology, changes between a domestic and a wild lifestyle related to locomotion and gait changes, have been estimated as a prevalent factor over load-carrying on structural changes of domestic donkeys limb bones (Shackelford et al., 2013). Despite this, archaeozoologists have always considered that the first observable morphological changes occurred only late in the domestication process and could not perceive its initial stages since they would not involve genetic isolation or selective reproduction (Vigne et al., 2011; Colledge et al., 2016). To develop new phenotypic proxy of the first steps of animal domestication, an experimental project (DOMEXP) has been performed to test whether growth in captivity of a wild ungulate would leave a morpho-functional imprint distinguishable from its natural habitat norm of reaction. Hence, early process of cultural control (Hecker 1982) could be traced from bones anatomy long before the so-called “domestication syndrome”. To test this hypothesis, the DOMEXP project used a genetically homogeneous wild boar (*Sus scrofa*) population to control the genetic and environmental factors influencing skeletal variation. From this populations, 6 months old piglets were captured after weaning to be reared until the age of two years under two mobility reduction regimes. Geometric morphometrics’ analyses have provided proof of concept that this mobility reduction in a wild ungulate population could leave morphometric prints on skulls (Neaux et al., 2021) and calcaneus (Harbers et al., 2020a). Microanatomical investigations have also revealed changes in the 3D topography of the cortical thickness in the humeral shaft (Harbers et al., 2020b). However, the impact of mobility reduction on the micraoanatomical features of the skeleton remained to be explored.

For this reason, this study investigates the microanatomical variation of the calcaneus from the experimental specimens of the DOMEXP project. Tarsal bones like the calcaneum are particularly considered as more informative than others in investigating the mechanical forces that apply to individuals during locomotion (Lovejoy et al., 1999; Carter & Beaupré, 2001; Pearson & Lieberman, 2004). Proximally articulated to the fibula and distally to the cuboid bone, the calcaneus is intensely stressed during locomotion in terrestrial tetrapods. It acts as a lever arm for the ankle extensor muscles, and is subject to high tensile, flexural, and compressive forces (Fig. 1; Hussain, 1975; Carrano, 1997; Bleefeld & Bock, 2002; Bassarova et al., 2009; Barone, 2017). The calcaneus is therefore a particularly interesting object of study to investigate differences in mobility between individuals (Harbers et al., 2020a). Moreover, the high compactness of the calcaneus makes it resistant to taphonomic alterations, which gives it a good general preservation in archaeological contexts (Binford, 1978).

**Fig. 1.**
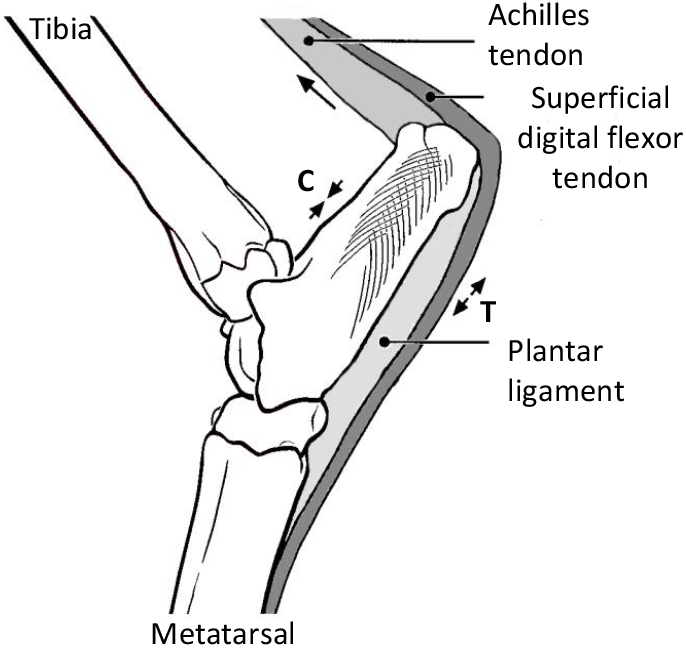
Lateral view of the tarsal region of a skeletally mature artiodactyl, showing the calcaneus with other associated bones, ligaments and tendons. The large arrow in the dorsal direction indicates the orientation of the force exerted by the Achilles tendon during paw extension, loading the dorsal cortex in compression (“C”) and the plantar cortex in tension (“T”). Modified from Su et al. (1999).

This study compares the calcaneus microanatomy variation between adult captive-bred and free-ranging wild boars to assess if captivity can induce microanatomical variation in a wild ungulate. We also include wild boar calcanei from a Mesolithic (ca. 8000 years old) context of hunter-gatherers in northern France to observe how much the variation in wild boar microanatomy has changed since the neolithisation of Europe and whether this proxy can be used to infer change in locomotor behaviour from archaeological calcanei.

## Methods

### Experimental protocol

24 wild boars aged six months were sampled from a control population living in a fenced forest of about 100,000 m^2^ in northern France. These specimens were then divided into two equal samples with the same sex ratio and reared until 24 month in two different contexts of reduced mobility: (1) a stall of 100 m^2^, where males and females were separated (each group had 45 m^2^) and (2) a wooded pen of 3000 m^2^. These two contexts represent respectively 99.9% and 97% of range reduction compared to the control population, which prevents the average daily distances measured in free boar populations (Palencia et al. 2019; Russo et al. 1997). The stall presented no opportunity for foraging. In the wooded enclosure, this opportunity was limited due to space limitations. Dry feed pellets suitable for feeding domestic pigs were provided to both groups.

### Specimens studied

Our dataset includes calcaneus bones from 47 specimens: 26 from the DOMEXP experimental farm - 11 stall specimens (stall), 10 pen specimens (pen), five from the control population (control) - 15 hunted wild boars from two Northern France forests: eight from the Chambord forest and seven from the Compiègne forest (Natural habitat) and six archaeological specimens from the Mesolithic contexts (Meso) of Noyen sur Seine (Mordant et al., 2013, Marinval-Vigne et al., 1989) in Northern France (Table 1).The six archaeological specimens from Mesolithic deposits have been accumulated by hunter-gatherer living in Western Europe before the Neolithic dispersal via the Mediterranean and Danubian pathways, which introduced domestic pigs from the Near East (Larson 2006) that interbred with local populations of wild boars (Frantz et al. 2020). Two specimens are radiocarbon dated from the Middle Mesolithic (between −8000 and −7300 cal. BP) and four specimens to the Final Mesolithic dated between −7000 and −6200 cal. BP (Mordant et al., 2013). All the specimens bar the archaeological samples have associated age and sex informations. Three of the archaeological calcanei belong to adult individuals since their proximal epiphysis is fused which is known to happen around 2 years in wild boars (Bridault et al., 2000).

**Table. 1.**
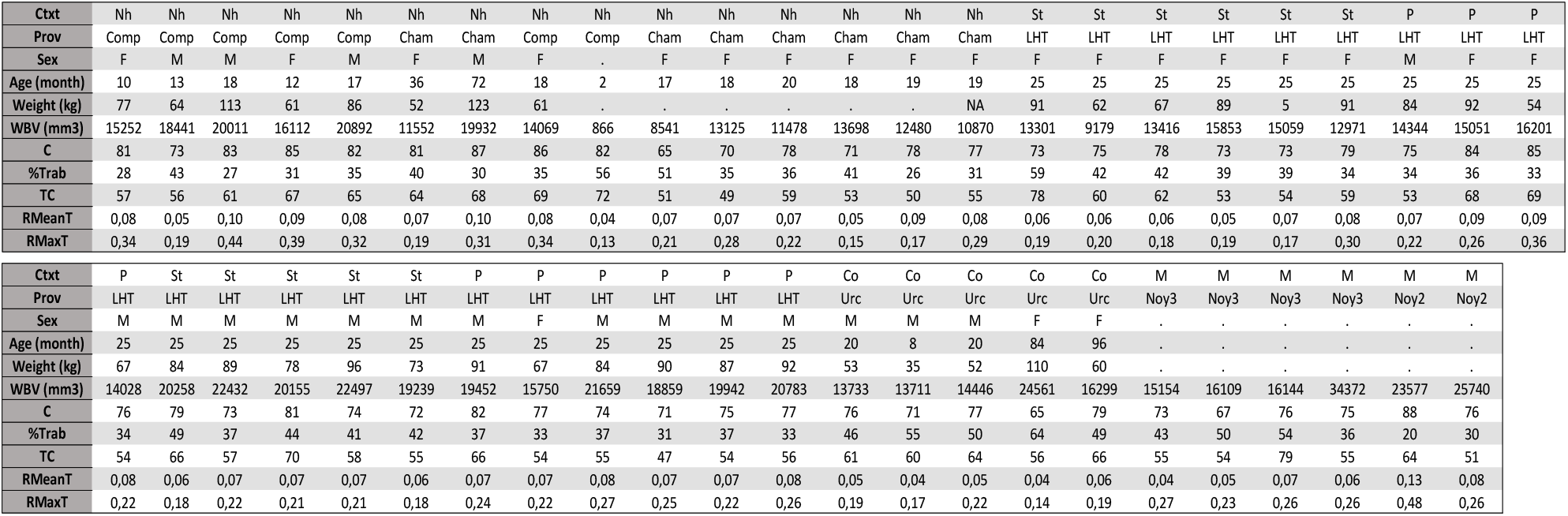
List of material and different parameters used in this study where each column corresponds to a specimen. Ctxt, Context; Prov, Provenance; WBV, total bone volume mm3; C, compactness ratio; %Trab, percentage of trabecular bone volume to cortical bone volume; TC, trabecular compactness; RMeanT, mean relative cortex thickness; RMaxT, maximum relative cortex thickness. Nh, Natural habitat; St, Experimental stall; Co, Experimental control; M, Mesolithic; P, Experimental pen; Comp, Compiègne; Cham, Chambord; Noy2/3, Noyen-sur-seine 2/3; Urc, Urcier.

### Data acquisition

All the calcanei have been scanned with high-resolution microtomography (EasyTom 40-150 scanner, RX Solutions) at the MRI platform, hosted at ISEM, University of Montpellier (UMR 5554); reconstructions were then performed using X-Act (RX Solutions).

### Virtual thin sections

For qualitative comparisons, three virtual sections (Fig. 2, Appendix 1,2 and 3) were made for each calcaneus, following Barone (2017) for terminology and orientations. The bones were oriented as follows: in dorsal view (Fig. 2a), bone’s axis is vertical and the fibular trochlea is oriented toward the observer, its dorsal part aligns with the contour of the bone’s medial border; in medial view (Fig. 2b) the *sustentaculum tali* is directed towards the observer, the observation angle is fixed when the anterior edge of the fibular trochlea is no longer visible upon rotation from the anterior view to the medial view. Sagittal sections (SS) run in dorsal view from the distal end tip and the midpoint of the thickness at the proximal epiphysis base (Fig. 2). The frontal sections (FS) extend from the distal end tip to the midpoint of the proximal epiphysis base (see purple arrows on Fig. 2b). Cross-sections (TS) are taken perpendicular to the FS plane at 1/3 of the total length of the bone (from the proximal epiphysis tip to the distal tip; Fig. 2). These sectional planes were chosen to depict large portions of the bone in order to analyze the microanatomical structure (e.g., trabecular network, cortical thickness) while being easily created with good reproducibility for all specimens. Virtual sections were created using VGSTUDIO MAX, versions 2.2 (Volume Graphics Inc.).

**Fig. 2.**
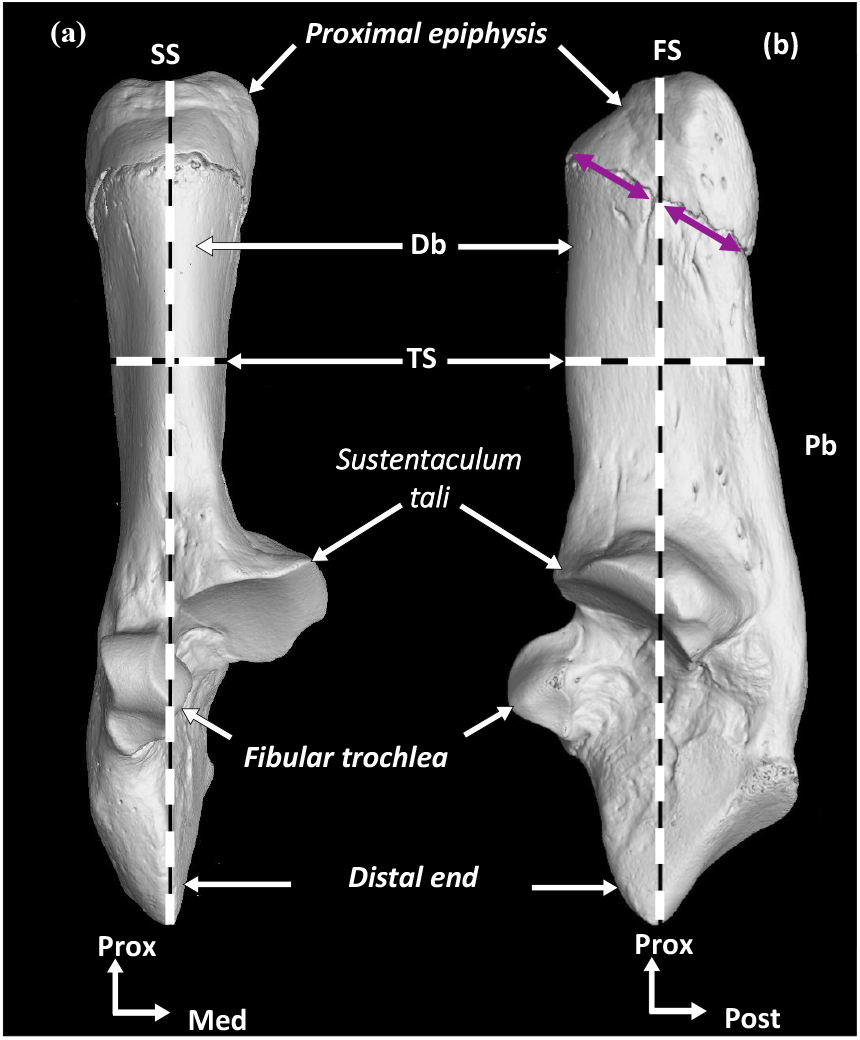
Calcaneus of *Sus scrofa*, specimen 2017-578 illustrating the planes of the virtual sections: SS, Sagital section; FS, Frontal section; TS, Transverse section. Db, Dorsal border; Pb, Plantar border. Purple arrows indicate the midpoint of the proximal epiphysis base.

### Calcaneus 3D mapping

To observe and measure the thickness variations of the compact cortex along the calcaneus, bone tissue was segmented (excluding soft tissue and cavities) using image data reconstructed with Avizo 9.4 (VSG, Burlington, MA, USA). Then, the outer cortex has been isolated from the trabecular bone, limited by the inner surface of the cortex for each bone. This segmentation step was done manually with a combination of Avizo’s “remove islands” (to eliminate isolated volumes that are too small) and “smooth labels” (to connect the slices selection more realistically) functions to optimize the segmentation repeatability and consistency. Then, the distances between the inner and outer surfaces of the cortex were calculated in Avizo 9.4 using the “surface distance” function. Finally, a distance isosurface was obtained with a colour gradient that appears on the external surface of the bone (Fig. 4; Appendix 4). This colour gradient showing the relative variation of cortical thickness within each bone, is specific to each specimen since it varies between the minimum and maximum cortical thicknesses, warmer colours being used for higher thicknesses, and colder colours for lower thicknesses. Therefore, two specimens with similar colorimetry may have different absolute cortical thicknesses.

**Fig. 3.**
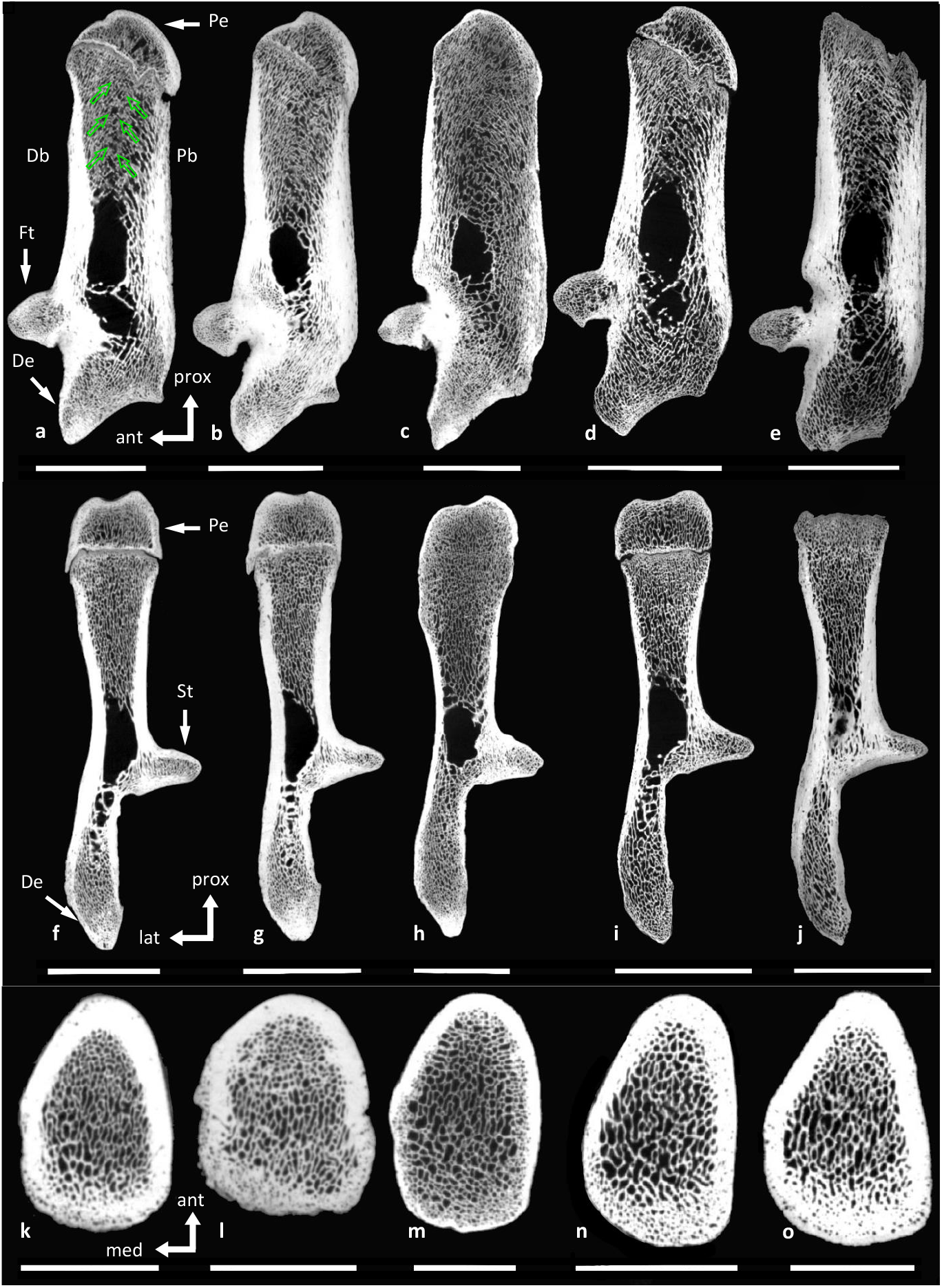
Virtual thin sections of the calcaneus of boars, a,f,k, 2017-555 (Experimental stall); b,g,l 2017-570 (Experimental pen); c,h,m Pradat187 (Experimental control); d,i,n, 2013-1286 (Natural habitat); e,j,o, Calc2139 (Mesolithic). Db, Dorsal border; De, Distal end; Ft, Fibular trochlea; Pb, Plantar border; Pe, Proximal epiphysis; St, *Sustentaculu tali*. Scale bars of sagittal and frontal sections equal 2 cm; Scale bars of transverse sections equal 1 cm.

**Fig. 4.**
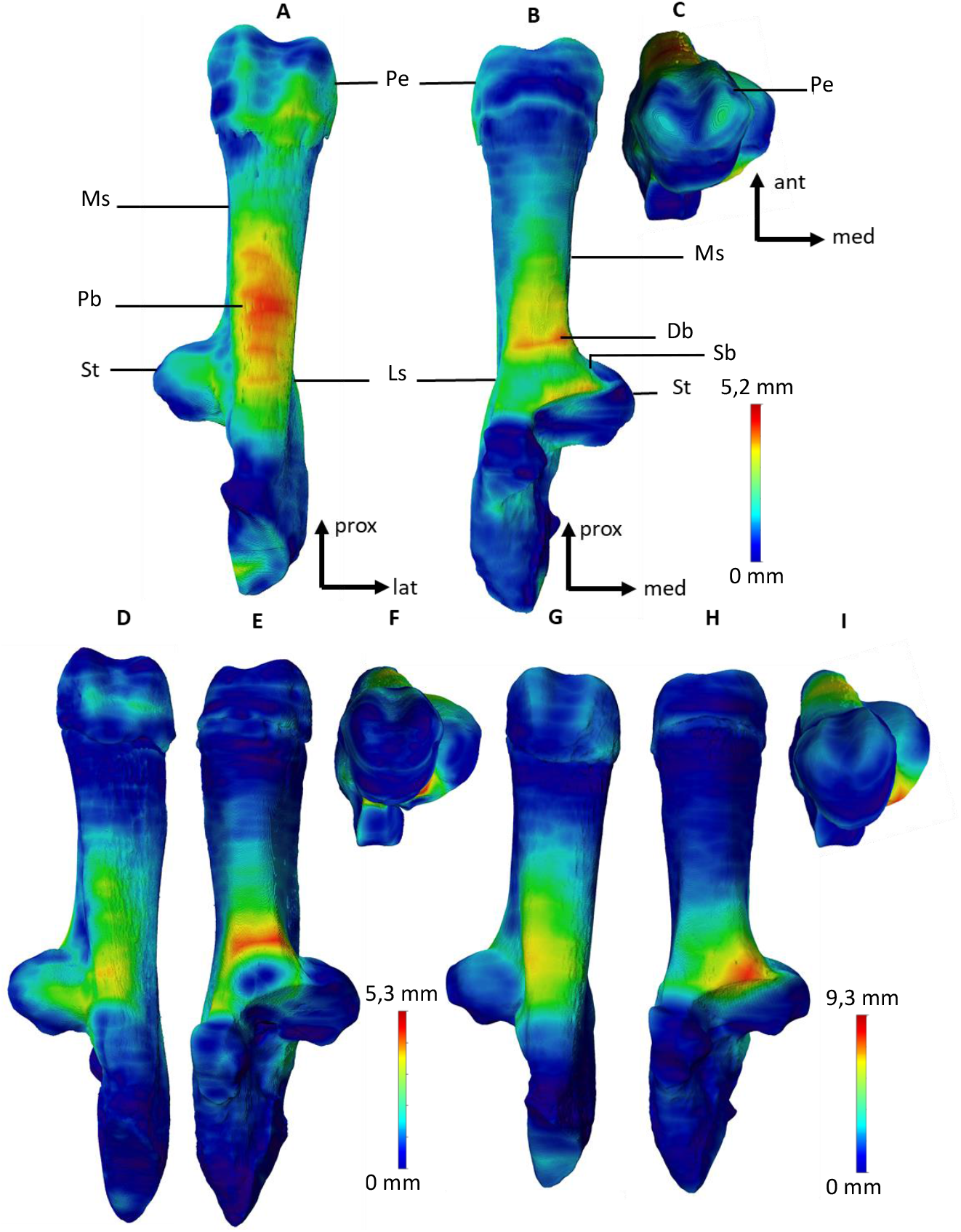
3D mappings of the *Sus scrofa* calcaneus relative cortical thickness. A-C specimen 2017-568; D-F Pradat 185; G-I 2013-1264. Anatomical abbreviations: Db, dorsal border; Ls, lateral side; Ms, medial side; Pb, plantar border; Pe, proximal epiphysis; Sb, Sustentaculum base; St, *Sustentaculum tali*. D, A and G are in plantar view; E, B and H are in dorsal view; F and I and C are in posterior view. Abbreviations for orientations, prox, proximal; lat, lateral; med, medial; post, posterior; ant, anterior.

### Quantitative parameters

Quantitative parameters used to characterize the internal structure of the bones are: (1) cortex/medullary area volumes, (2) overall bone and trabecular tissue compactness, and average and (3) maximum cortical thicknesses following Houssaye et al. (2021). Most are ratios produced using the volume values obtained from the “material statistics” function in Avizo after the segmentation and cortex isolation steps. The parameters used in the statistical analyses in this study are: 1) whole bone volume in mm^3^ (WBV), as an indicator of size; 2) bone compactness (C= bone tissue volume*100/WBV); 3) relative trabecular bone tissue fraction (%Trab = trabecular bone tissue volume*100/bone tissue volume); 4) trabecular compactness (Tc= trabecular bone tissue*100/trabecular volume). From the mean (MeanT) and maximum (MaxT) cortical thicknesses that were obtained directly in Avizo 9.4 using the ‘distance’ function were calculated 5) RMaxT and 6) RMeanT, relative maximum and mean thicknesses, by dividing MaxT and MeanT by a mean radius r, obtained from the whole bone volume and considering that the calcanea are cylinders (as v=πr^2^ h, r=√(v /hπ)).

### Statistical analysis

Statistical tests and graphical representations were performed in R (R Core Team. 2017) using the Rstudio software (see appendix 5 and 6 for the script and the data Table). A linear regression model (“lm” function of the “stats” package) as well as a regression coefficient (“cor” function of the “stats” package) were used to assess the linear relationships between the variables (1) whole bone volume (WBV; considered as an estimate of bone size), (2) weight, and (3) age of each individual with all the other quantitative parameters used (Table 2). To explore the distribution of specimens based on their quantitative microanatomical parameters and the variation patterns, we performed a standardized Principal Component Analyses (PCA; “dudi.pca” function of the ade4 package; David and Jacobs 2014).

**Table. 2.** Values obtained for tests of the effect of total volume, age, and weight on the different parameters and the PCA axes. r: correlation coefficient; p: p-value of the linear regression model; WBV: whole bone volume; RMeanT, mean relative cortical thickness; RMaxT, maximum relative cortical thickness; PC1, PC2: position of individuals on the first two axes of the PCA.

|  | WBV | Age | Weight | C | %Trab | TC | RMeanT | RMaxT | PC1 | PC2 |
| --- | --- | --- | --- | --- | --- | --- | --- | --- | --- | --- |
| WBV<br>n=46 | X | X | X | $r=0,07$<br>$p=0,63$ | $r=-0,13$<br>$p=0,38$ | $r=-0,05$<br>$p=0,75$ | $r=0,13$<br>$p=0,37$ | $r=0,20$<br>$p=0,18$ | **$r=-0,46$**<br>**$P<0,01$** | **$r=0,36$**<br>**$p=0,01$** |
| Age<br>n=40 | $r=0,30$<br>$P=0,06$ | X | X | $r=0,47$<br>$p=0,92$ | **$r=0,32$**<br>**$p=0,04$** | $r=0,17$<br>$p=0,29$ | $r=-0,19$<br>$p=0,23$ | $r=-0,20$<br>$p=0,21$ | $r=0,07$<br>$p=0,65$ | $r=0,25$<br>$p=0,12$ |
| Weight<br>n=34 | **$r=0,56$**<br>**$p<0,01$** | $r=0,27$<br>$p=0,12$ | X | $r=0,08$<br>$p=0,65$ | $r=-0,16$<br>$p=0,35$ | $r=0,02$<br>$p=0,93$ | $r=0,30$<br>$p=0,08$ | $r=0,26$<br>$p=0,14$ | $r=0,34$<br>$p=0,05$ | $r=0,31$<br>$p=0,07$ |

To estimate the role of factors such as Sex (male/female/indeterminate), origin (Compiègne/ Chambord/ Urcier/ La Haute Touche/ Noyen sur Seine) and mobility status (Natural habitat /Experimental control/ Experimental pen/ Experimental stall/Mesolithic wild boars) in the variation of the quantitative microanatomical parameters, we used analyses of variance (ANOVA; function “anova_test” of the “rstatix” package) after checking the conditions of normality (function “shapiro_test” of the “rstatix” package) and homogeneity of variances (function “levene_test” of the “rstatix” package). When overall difference is significant, we computed pairwise comparison tests using Tukey hsd tests (“tukey_hsd” function of the “rstatix” package). When the variables did not meet the conditions of homogeneity of variances and/or normality, we used the kruskal-Wallis test (function “Kruskal_test” of the “rstatix” package), a non-parametric alternative to ANOVA. When this test is significant, a Dunn’s test (“dunn_test (p.adjust.method=“bonferroni”)” function of the “rstatix” package) is used to compare pairwise differences between the groups concerned.

MANOVA (“res.man” function) was also used to test the overall difference in microanatomical variables between mobility contexts.

## Results

### Qualitative descriptions

#### General microanatomical pattern

The general microanatomical pattern of the wild boar calcaneus is close to the structure of long bones of terrestrial mammals, with the cortex forming a tubular diaphysis but with a very short diaphyseal part (sections Fig. 3). The proximal epiphysis is not fused to the rest of the bone for the most part of the specimens, except for the oldest. The cortex surrounds cancellous bone (trabecular bone and intertrabecular spaces), which is essentially quite dense, and a small open medullary cavity, about 1cm (Pradat187; Fig. 3c,h) to 2 cm long (2013-1286; Fig. 3d,i) and 1 cm wide, at the level of the *Sustentaculum tali* (FigXa-j). The thickness of the trabeculae is relatively homogeneous along the calcaneus, except around the medullary cavity, where they are generally thicker (Fig. 3a-j). Also, the trabecular density is heterogeneous, with some specimens having many trabeculae (2017-570; Fig. 3b,g,l) while others have twice as less but thicker trabeculae (2013-1286; Fig. 3 d,I,n). Finally, the bone density in the distal part strongly varies between individuals from compact (2017-570; Fig. 3b,g) to spongious (Calc2139; Fig. 3e,j).

#### Sagittal sections

In all specimens, the dorsal and plantar margins (Db and Pb) show a relatively high cortical thickness, especially at mid-diaphysis. Spongious bone shows anisotropic trabeculae (i.e. with a preferential direction) above the medullary cavity in the bone center (Fig. 3a-e). Anisotropic trabeculae follow the two main directions that are represented with green/outline arrows in the figure 3a and schematized with intersecting lines in figure 1. The cortical thicknesses of the plantar and dorsal margins vary from thick (1 cm in specimen 2017-570; Fig. 3b) to proportionally twice as thin (0.5 cm in Pradat187; Fig. 3c). A variation in the cortical thickness of the plantar border of the proximal epiphysis is also observed, it is very thin (1mm) in some specimens like Pradat 187 (Fig. 3c) while others, such as 2017-555 (Fig. 3a), show a clear thickening of the cortex (2mm) in this area. Similarly, bone is highly compact next to the fibular trochlea for numerous specimens (Calc2139; Fig. 3e) or rather spongious for some individuals (2013-1286; Fig. 3d).

#### Frontal sections

All individuals have a cortex that varies relatively little in thickness (about 2-3mm) on the medial and lateral sides (Fig. 3f-e). In contrast, the cortical thickness in the proximal epiphysis varies from thin (0.5 cm in specimen 2013-1286; Fig. 3i) to twice thicker (0.3 cm in specimen 2017-555; Fig. 3f). Similarly, the compactness and the cortical thickness of the *sustentaculum tali* varies between individuals, some of which show compact bone and thick cortex (Padat187; Fig. 3h) while others show spongious bone and thin cortex (2013-1286; Fig. 3i).

#### Transverse sections Transverse sections

The transverse sections’ shapes are generally oval and elongate (2017-555; Fig. 3k) but several specimens present a rounder section (2017-570; Fig. 3l). The cortical thickness is fairly constant across the sections, but some specimens show a cortical thickening at the plantar border (2013-1286; Fig. 3n).

None of the variation observed between specimens is clearly qualitatively associated to any main parameter of the study, namely the context, provenance, sex, size or weight.

### 3D mapping of the cortical thickness

The 3D mappings of the cortical thickness and its variation across each bone, are quite similar between specimens (Appendix 1). In agreement with the observation of the virtual sections, there is a fairly extensive area with greater cortical thickness at the plantar border of the calcaneus (Pb; Fig. 4A). Although generally less extensive, thickenings of the cortex can also be noted on the dorsal margin (Db) and on the dorsal base of the sustentaculum tali (Sb; Fig. 4B). There is little thickening of the cortex on the proximal end (Pe). Finally, there is no lateral or medial thickening noted.

Twenty-five of the 47 specimens show strong cortical thickening at the plantar margin (Pb; Appendix 1), like in 2017-568 (Fig. 4A); this is less visible in the 22 others, like Pradat 185 and 2013-1264 (Fig. 4D; Fig. 4G). Twenty-three specimens have thick cortical bone at the dorsal margin (Db; Table 1; Appendix 1), like in Pradat 185 and 2017-568 boars (Fig. 4B and Fig. 4E), whereas the others do not show such thickening like the calcaneus of boar 2013-1264 (Fig. 4H). Twenty-height specimens show a relative thickening at the base of the sustentaculum (Sb; Table 1), this widening is particularly noticeable in boar 2013-1264 (Fig. 4H) and absent in others like Pradat 185 (Fig. 4E). A slight recurrent cortical thickening on the distal end is observed in part of the specimens like 2017-568 (Fig. 4a). Twenty-seven specimens have a slightly thicker cortex on the proximal end (Pe) such as 2017-568 (Fig. 4C), 15 show no or very little thickening, such as Pradat185 and 2013-1264, respectively (Fig. 4F and Fig. 4I). Four of the proximal ends of the Mesolithic specimens were not found because they were not fused to the rest of the bone.

### Quantitative analyses

#### Microanatomical covariation with Weight, age size and sex

The Whole bone volume (WBV) is expectedly only correlated to the body weight and the PCA axes (Table. 2). Only %Trab is significantly correlated with specimen age with a slight increase in the proportion of trabecular bone over cortical bone as boars get older (r=0.32).

Variables C, %Trab, RMeanT, RMaxT, weight, and TC did not differ between males and females. Only whole calcaneus volume (WBV) variation differs with sex (Kruskal Wallis: p<0.01) with males larger than females (Fig. 6). Mesolithic specimens (sex unknown) have larger calcanei than present-day males (Dunn’s test: p<0.01).

**Fig. 5.**
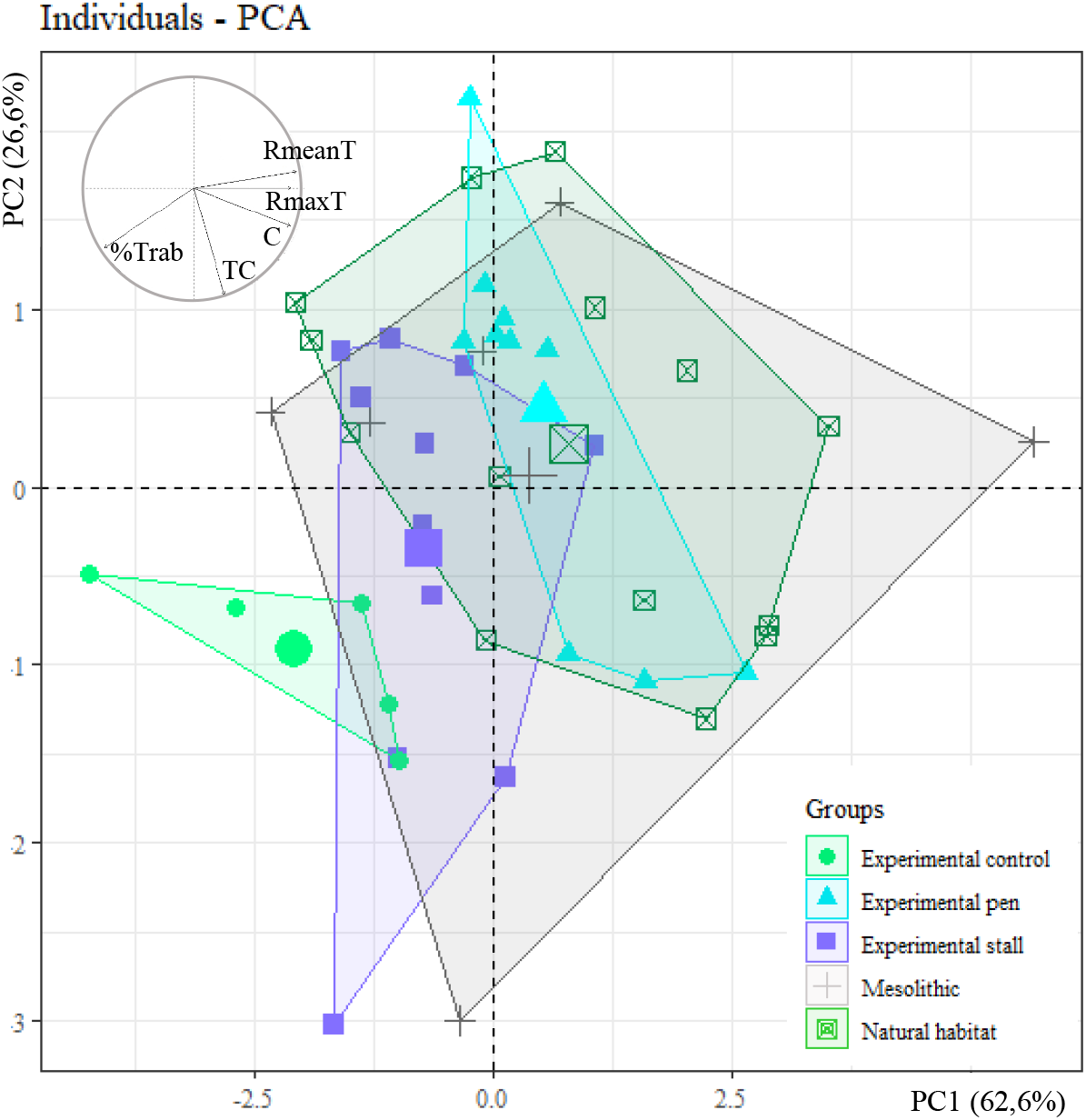
Distribution of the 46 specimens and their associated context groups on the first two axes of the PCA computed on: WBV, total bone volume; RMeanT, relative mean bone thickness; RMaxT, relative maximum bone thickness; C, compactness ratio; TC, trabecular compactness ratio; %Trab, percentage of trabecular bone volume to cortical bone volume. meso, Mesolithic; nat hab, Natural habitat.

**Fig. 6.**
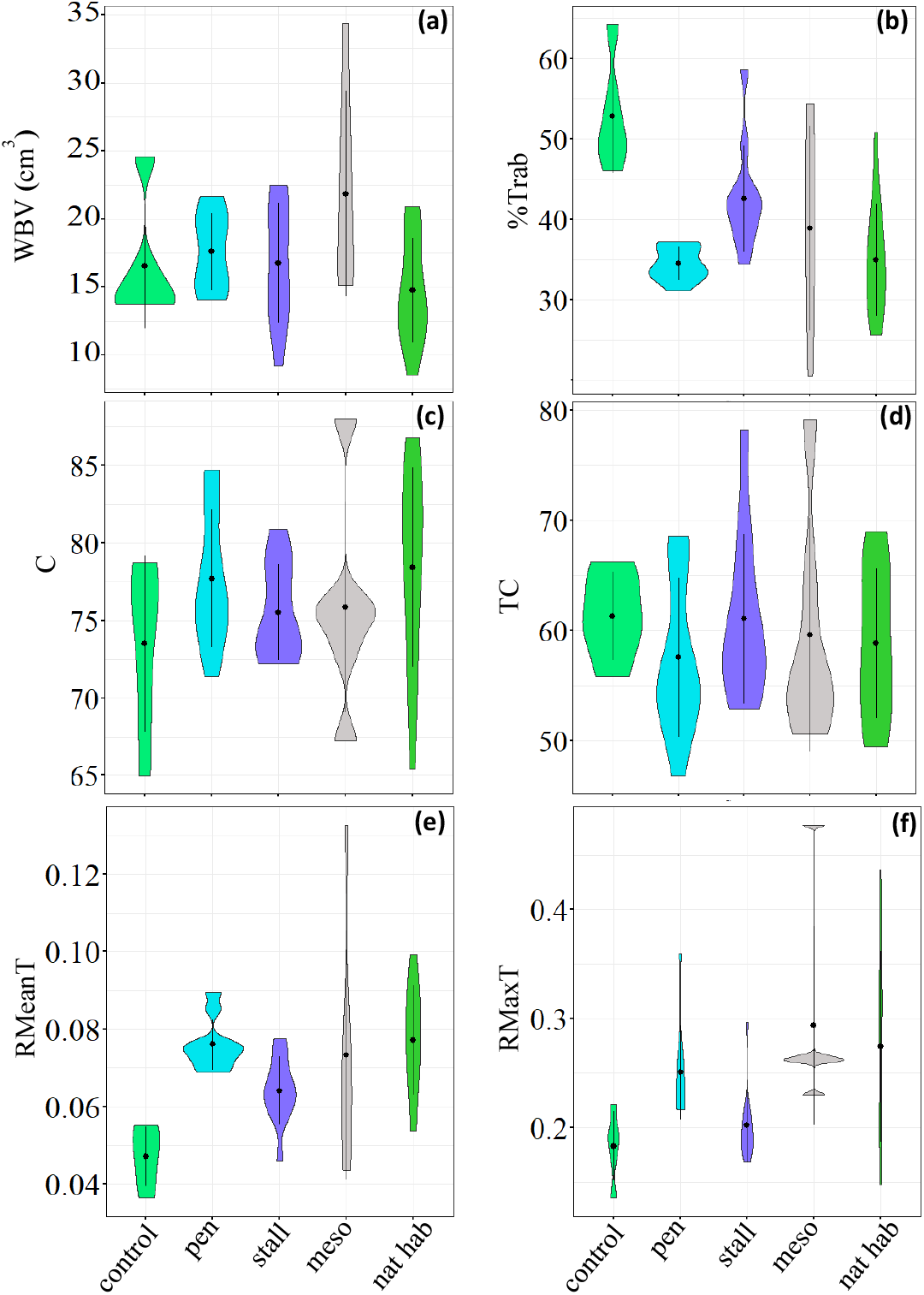
Calcaneus microanatomical variation in *Sus scrofa* from experimental populations and Mesolithic contexts. WBV, whole bone volume; RMeanT, relative mean cortical thickness; RMaxT, relative maximum cortical thickness; C, compactness ratio; TC, trabecular compactness ratio; %Trab, percentage of trabecular bone volume to cortical bone volume. meso, Mesolithic; nat hab, Natural habitat.

#### Patterns of calcanei microanatomical variations and mobility contexts

Specimen 2013-1287, corresponding to the youngest individual (2 months), was removed before performing PCA (n=46; Fig. 5) and other quantitative analyses. Axes 1 and 2 of the PCA explain 75.3% of the total variance. Furthermore, contribution of the variables to the axes (Fig. 5), show that RMeanT and RMaxT covary while WBV and %Trab vary in opposite ways. The first axis is influenced by the cortical thickness (TC), %Trab and C, while only TC greatly structures the second axis. WBV is correlated with the PCA axes whereas age, weight and the microanatomical parameters are not (table 2). The different mobility contexts induce significant microanatomical differences (MANOVA: p-value<0.01). On the PCA (figure 5), experimental penned, stalled, and control wild boars are quite distinct along PC1, whereas the Mesolithic specimens of Noyen-sur-Seine covers almost all variation along PC1. Differences across the locomotor contexts are observed for RMeanT (Kruskal Wallis: p<0.01), RMaxT (Kruskal Wallis: p<0.01) and %Trab (Kruskal Wallis: p<0.01). RMeanT differences are significant between wild boars from the Natural habitat and the Experimental control groups (Dunn test: p<0.01) and between Experimental control and Experimental pen groups (Dunn’s test: p<0.01). RMaxT is significantly different between Experimental control and Mesolithic wild boars (Dunn’s test: p<0.01). %Trab is significant different between Experimental control and natural habitat (Dunn’s test: p<0.01) and between Experimental pen and control (Dunn’s test: p<0.01). Thus, the pen-reared boars have a higher average cortex thickness (RMeanT) than the control group, for which, conversely, it is the percentage of trabecular bone (%Trab) that is higher (Fig.6). The stall group had intermediate RMeanT, RMaxT and %Trab values compared to the pen and control groups. In addition, there is a generally greater standard deviation in the Natural habitat and Mesolithic groups for all parameters, except TC, for which individuals in the stall group are slightly more dispersed than those in the Natural habitat group. On the other hand, no group is distinguished along PC2, the variables C and TC did not vary according to context (ANOVA; p-value C =0.39; p-value TC =0.82; Fig. 6); Finally, the whole volume (WBV) of the calcaneus differed slightly between groups (ANOVA: p=0.04) but only the difference between hunted Mesolithic and modern wild boars in their natural habitat was significant (Tukey hsd test: p=0.02; Fig. 6), with Mesolithic individuals being larger.

## Discussion

### 1. Overall calcaneal microanatomy in wild and captive wild boars

The qualitative description of the 3D maps and virtual thin sections identified a strong cortical thickening on the plantar and dorsal borders (Pb and Db; Fig. 4). These observations are consistent with the main constraints identified in the calcaneus of artiodactyls with significant compression, bending, and tension forces on the plantar and dorsal borders (Lanyon 1973; Fig. 1). Thickenings of the plantar and dorsal margins have also been identified for wild deers (Skedros et al., 2001) and pasture-raised domestic sheeps (Skedros et al., 2007). Moreover, the anisotropric properties of the boar specimens trabeculae to follow an antero-posterior orientations (outline arrows Fig. 3a) are congruent with the distribution of constraints mentioned above, as internal bone structure organized itself to better respond to stress (Wolff, 1986; Ruff et al., 2006; Van der Meulen et al., 2006).

Although the entheses (i.e., areas of ligament and tendon insertion (Djukic et al., 2015)) are regions of stress concentration, these areas show little or no effect on the microanatomy of boar calcaneus. The loads applied to the calcaneus of artiodactyls are primarily shared between the long plantar ligament and the Achilles tendon, which connects the calcaneus to the gastrocnemius and soleus muscles, the latters forming the sural triceps (Lanyon,1973; Woo et al., 1981; Skedros et al., 2001; Skedros et al., 2007; Barone, 2020). However, while the long plantar ligament attaches all along the plantar border, only a part of this edge is thickened in the boar calcaneus. Furthermore, an important thickening of similar proportion is also observed on the dorsal edge, whereas this bone side shows no enthesis. The Achilles tendon insertion at the proximal end of the calcaneus is itself covered by a tendinous structure, the calcaneus cap of the superficial flexor digitorum muscle (*m. flexor superficialis)*, taking an attachment point from the tip of the proximal end of the calcaneus to phalanges (Su et al., 1999; Barone, 2017; Fig. 1). A slight cortical thickening on the proximal epiphysis of our specimens coincides with the insertion of the tendons of the *m. flexor superficialis* muscles of the toes and of the *m. gastrocnemius* muscles (Bénévent & Bressot, 1968; Barone 2017).

Regarding the articular surfaces, the *sustentaculum tali* is a strong protuberance on the medial side forming an articular surface with the talus, the fibular trochlea is an articular surface for the malleolar bone and the end of the distal part articulate with the cuboid bone (Barone, 2017), recurrent slight cortical thickening is also observed in those regions.

Overall, the microanatomy seems to reflect the tension and compression forces with a strong cortical thickening on Pb and Db as well as the anisotropy of the trabeculae. The areas of contact with other bones are also represented with important bone density on the fibular trochlea, the *sustentaculum tali* and the end of the distal part.

### 2. Intra individual variation in calcaneal Microanatomy

Beyond the general pattern, no clear relationship is observed between the variability of the microanatomical parameters and the factors explored in this study (context, sex, weight, size). However, we found important inter-individual differences, notably in the extent and depth of the cortical thickness of the plantar and dorsal edges, and on the proximal epiphysis; the length of the medullary cavity; the number and thickness of the trabeculae; the bone tissue density of the *sustentaculum tali*, at the distal part of the bone and next to the fibular trochlea; and the transverse sectional shape. This suggests that other factors influencing bone development during growth must be explored to further understand inter-individual disparity.

Only a few correlations (positive or negative) of the microanatomical variables with bone size, weight, and age of the individuals (Table 2), are significant. These three parameters, therefore, have a limited impact on the microanatomical organisation. Body weight is only correlated with bone size (WBV). WBV influences the PCA axes (Table 2) while, paradoxically, bone size is not directly correlated with any of the microanatomic variables in isolation, which is congruent with the observations of the sections and 3D maps that have not found any link between specimens’ size and their microanatomy. Thus, it is the covariation between the variables that are themselves weakly correlated with whole volume that makes the relationship between whole volume and all variables significant. Nevertheless, although age is not one of the parameters on which this study focuses, a weak correlation was found between age and the trabecular percentage (%Trab; r=0.32), but trabecular percentage is not significantly correlated with bone size since age and bone size are neither significantly correlated. The increase in the proportion of trabecular bone tissue with age is not related to an increase in trabecular compactness (TC) because this parameter is not significantly correlated with age (p=0.29); the volume of the medullary zone tends to increase since the cortex becomes proportionally thinner. This result is surprising because the opposite phenomenon occurs in the calcaneus of deer (Skedros et al., 2001) and sheep (Skedros et al., 2007), where a thickening of the cortex is observed with size in relation to the medullary zone. Conversely, while it is surprising that age does not correlate with whole bone volume (WBV; Table2), this shows that age and volume do not follow a linear relationship or that intraspecific variability in calcaneus size between specimens exceeds the effect of growth. However, our sample does not adequately test the relationship between the variables and ontogeny because of the large proportion of individuals of the same age (25 months).

### 3. Change in mobility regime and calcaneal Microanatomy

Despite the lack of directly observable influence of the mobility context over the calcaneal microanatomy, we found quantitative effect of mobility differences in the cortical thickness (RmeanT and RmaxT), trabecular percentage (%Trab) and in the overall variation patterns (PCA), indicating that difference in mobility context influences the microanatomical characteristics of the calcaneus, although not in a strongly discriminatory pattern. However, we didn’t find the expected microanatomical proximity of wild boars living in their natural habitat and their dissimilarity from wild boars kept in captivity, as seen in previous studies on the calcaneus 3D externalshape and form (Harbers et al.2020). The variations in relative mean cortical thickness (RMeanT) illustrates the general trend of variations related to the mobility regime (Fig. 6). While we expected to observe similar average thicknesses between animals that had similar mobility conditions (e.g. Natural habitat and Experimental control), we found that control individuals have a much lower cortical thickness than wild individuals from natural habitat. Thus, modern and Mesolithic wild boars hunted in their natural habitat display similar microanatomy with the wild boars which grew in a very small living space (enclosure of 45m^2^ in a hangar of 100m^2^.). The most divergent microanatomy from the wild boar norm of reaction have been observed in the control populations from the wild boar farm of Urcier, which have a much lower cortical thickness than wild boars in their natural habitat. All these results suggest that the microanatomy of the *Sus scrofa* calcaneus does not strongly reflect the mobility regime, contradicting the strong microanatomical signal associated with locomotor restriction evidenced in battery chickens, including osteoporosis related to inactivity (Rath et al., 2000). However, the locomotor restrictions of these reared animals are generally greater than those imposed on the animals of this study.

### 4. Greater microanatomical variability in modern and ancient wild boars hunted in their natural habitat

The six Mesolithic calcanei from the archaeological site of Noyen-sur-Seine show similarity for all parameters and variables with the other groups (Fig. 6). Thus, the Mesolithic individuals share the same micronatomy with their modern relative. We also found greater microanatomical variability in wild boars populations both modern and Mesolithic. External factors were much less controlled than for the DOMEXP groups, thus resulting in more elements to affect bone plasticity. In addition, the animals from Compiègne and Chambord (Natural habitat) and NoyenEns2 and NoyenEns3 (archaeology) are different populations so they have greater genetic variability in comparison to the DOMEXP group. Furthemore, the Mesolithic specimens belong to individuals from multiple generations separated by several centuries. This genetic variability is much a substantial factor that has a greater impact on the observed phenotype than the intrapopulation variability related to motricity in this study. Consequently, when several populations are included in the same group (Natural habitat and Mesolithic), their variability exceeds that observed between the same population placed in different locomotor contexts (stall, pen, control). Thus, the microanatomy of the calcaneus appears to be more affected by population differences than by the locomotor context in which the animals grew.

In addition to this explanation, the wider locomotor regime of wild boars in their natural habitat would foster greater ecophenotypic variation compared to captive specimens with reduced mobility and more stereotypical locomotor behaviour. One the one hand, wild boars in nature have a locomotor repertoire that must respond to several problems that are not encountered in captivity, such as foraging or escape (Spitz & Janeau, 1990, 1995). Their daily travel is generally less than 10km but it could be up to 80 km in one night (Keuling et al., 2009), peak speeds of 40 km/h and high jumps up to 1.5 m have also been observed (Baskin & Danell, 2003). One the other hand, there is no study describing the skeletal repercussion of stereotypic behaviours. However, because these types of behaviours performed to compensate for lack of activity are induced during significant psychological depressions in individuals (Rushen, 1993; Andre, 2007), other experimental approaches involving living specimens are not desirable.

Also, diet plays a role in the development of the skeleton (Randoin & Causeret, 1945). Experimental specimens were fed nutritionally balanced pellets (15% protein) intended for pig breeding to ensure consistent growth and bone formation. In their natural habitat, food availability, seasonal, and geographical variations are major factors influencing food selection by wild boars (Ballari, & Barrios-García, 2014), Thus, the diet factor possibly also contributes to the high variability observed in wild (free-ranging) individuals. Furthemore, it is likely that there are differences in substrates between the forests and areas in which these boars lived; this may have important implications for the autopod (Kappelman, 1988). Indeed, the substrate factor seems potentially important for the stall-reared group where the flat ground, was covered with moss and straw mats, whereas the more irregular natural terrain of wild boars (like control group) implies variable and multidirectional soil reaction forces (Hanot et al., 2017). The group of pens were raised on a flat terrain covered with grass and a few trees.

The overview of the different studies related to the DOMEXP project show that bone plasticity associated with domestication varies between bones. The most surprising result of the present study is that the bone plasticity of the external morphology of wild boar calcaneus’ is more variable than its microanatomy, although the latter is considered to be more plastic than bone external morphology (Dumont et al., 2013; Kivell, 2016; Vlachopoulos et al., 2017). Since the results of the study show that boars living in different contexts can have the same microanatomical pattern, calcaneal microanatomy cannot be used to infer a captive lifestyle. However, a strong diversity is noted in the microanatomy between wild and Mesolithic specimens. A better understanding of the factors that regulate calcaneal variability would possibly allow inferences related to habitat (type of soil, open or closed environment) or diet.

## Supporting information

Appendix 1 frontal sections

Appendix 2 sagittal sections

Appendix 3 transversal sections

Appendix 4 3D mapps of the cortical thickness

R-script

Appendix 6 data table for R.txt

## Acknowledgements

We thank the reviewers Ignacio Aguilar Lazagabaster and Max Price as well as our recommender Nimrod Marom for their helpful and positive comments that have improved this work. Thanks to Florian Bogaert, who helped us to realize the segmentation of a part of specimens. Preprint version 5 of this article has been peer-reviewed and recommended by Peer Community In Archaeology (https://doi.orq/10.24072/pci.archaeo.100023).

## Data, scripts, code, and supplementary information availability

Data, script, and supplementary informations are available online: https://www.biorxiv.org/content/10.1101/2022.08.22.504790v1.supplementary-material?versioned=true

## Conflict of interest disclosure

The authors declare that they comply with the PCI rule of having no financial conflicts of interest in relation to the content of the article. Alexandra Houssaye is recommender of PCI Paleontolgy.

## Funding

This research has been funded by the project Emergence SU-19-3-EMRG-02. This research has also benefited from financial supports of the Muséum national d’Histoire naturelle (Paris) and the CNRS INEE (Institut écologie et environnement).

This research was funded by ANR through the DOMEXP project (ANR\u000213-JSH3-0003-01), the LabEx ANR-10-LABX-0003-BCDiv, in the programme ‘Investissements d’avenir’ ANR-11-IDEX-0004-02, programme Emergence SU-19-3-EMRG-02.

## Notes

### Competing Interest Statement

The authors have declared no competing interest.

### Summary of Updates

add script and table

