## Appendix 2 sagittal sections for "Can growth in captivity alter the calcaneal microanatomy of a wild ungulate?"

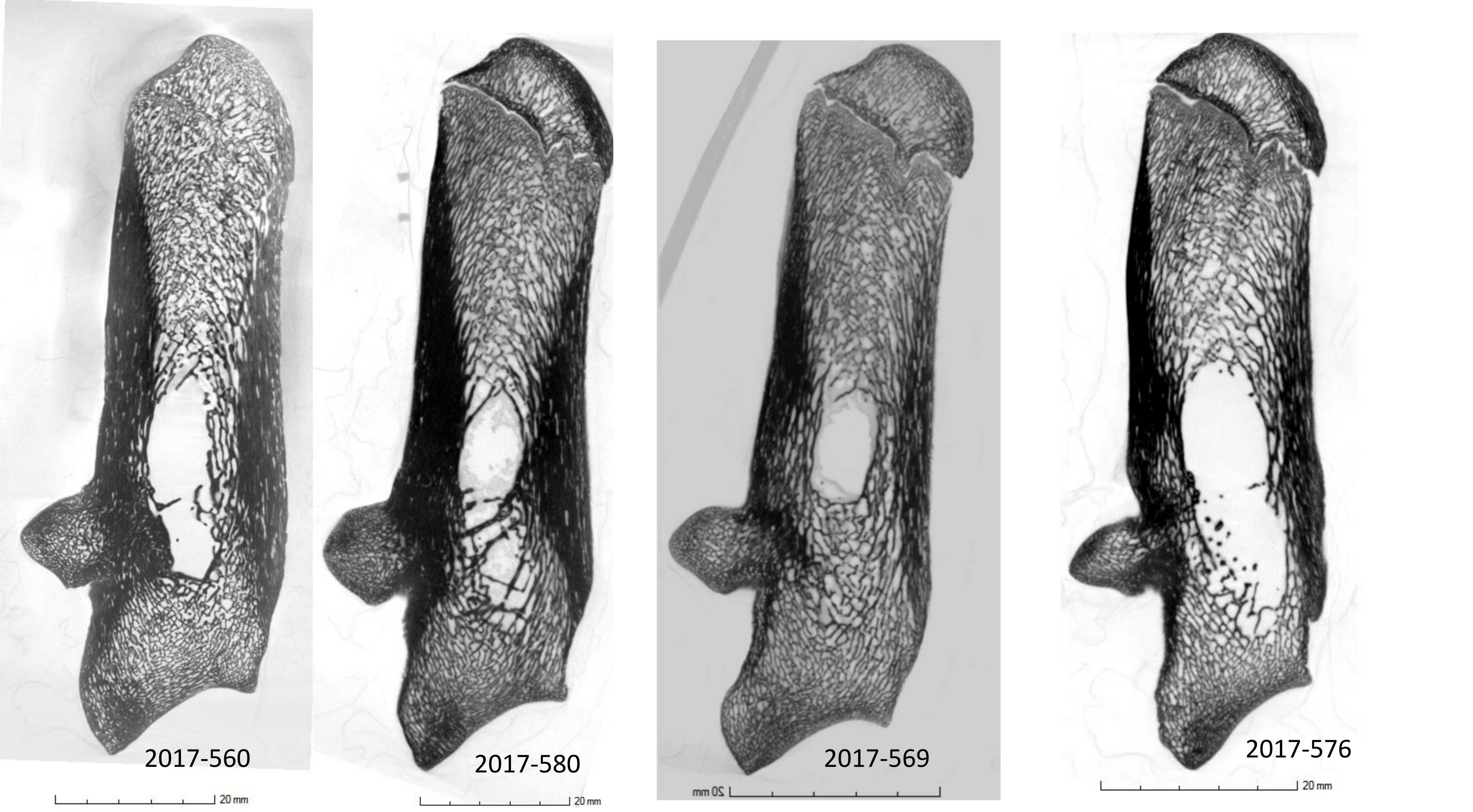
2017-560
20 mm
2017-580
20 mm
2017-569
2017-576
20 mm

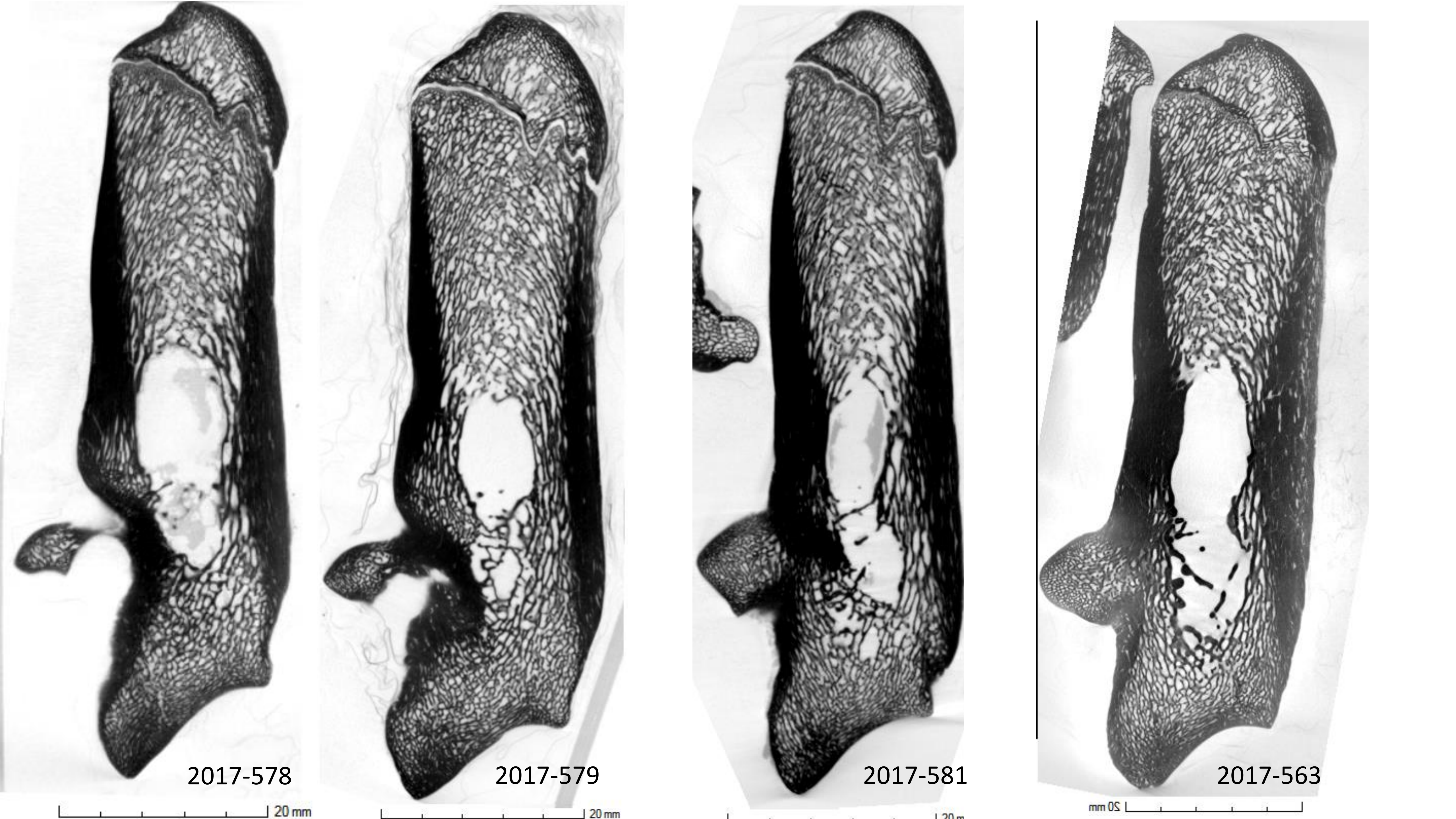
2017-578
20 mm
2017-579
20 mm
2017-581
2017-563

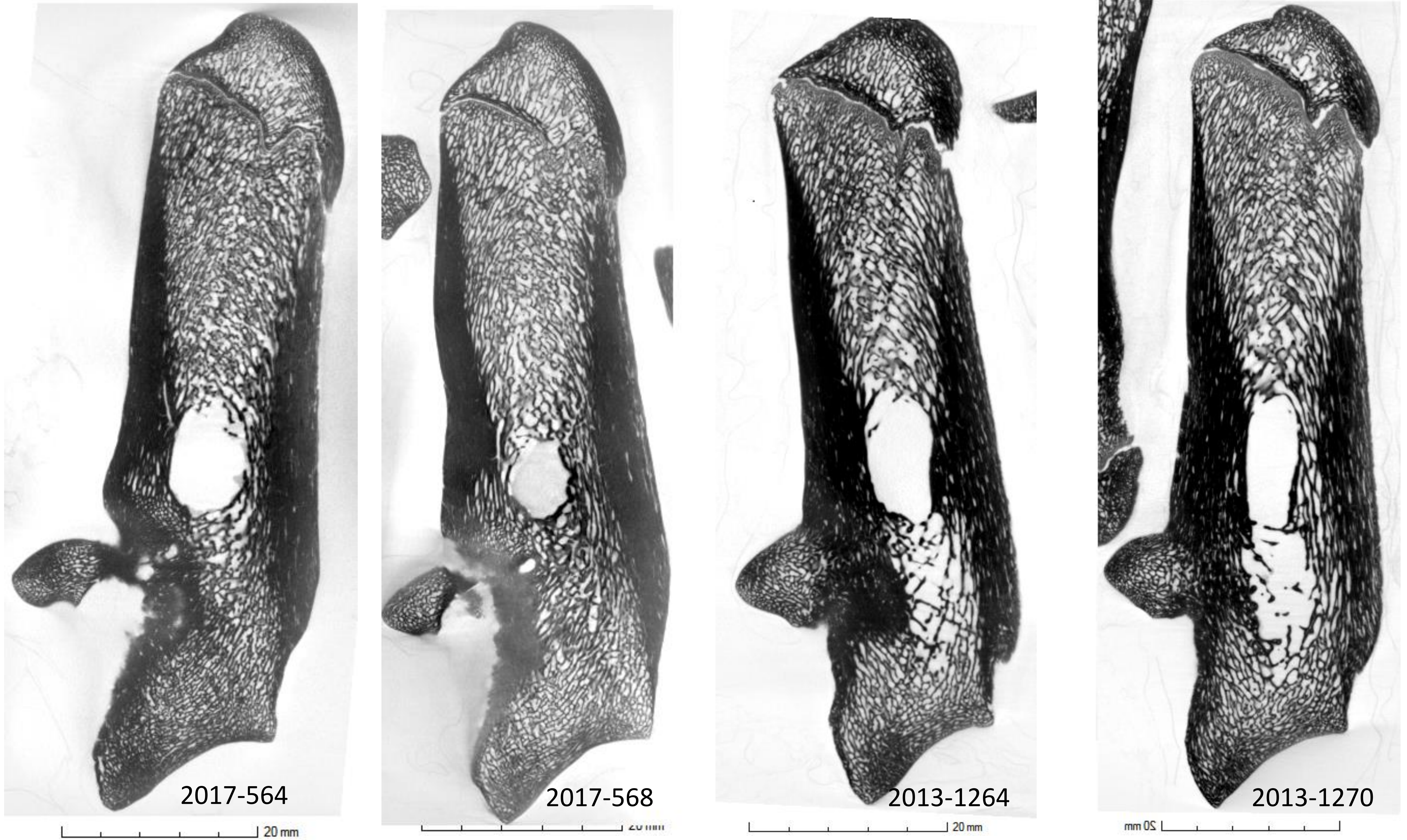
2017-564
20 mm
2017-568
2013-1264
20 mm
2013-1270
20 mm

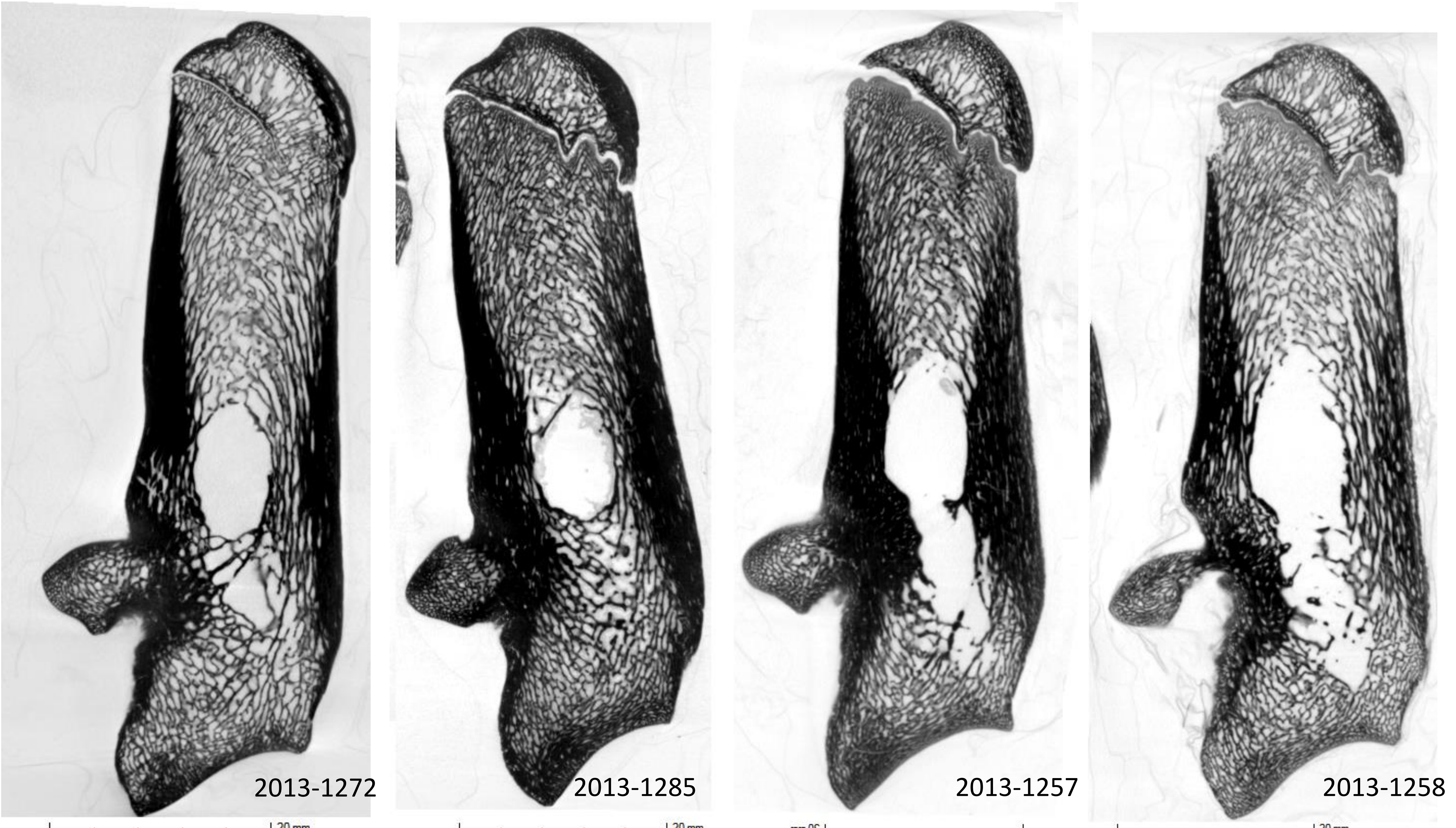
2013-1272
2013-1285
2013-1257
2013-1258

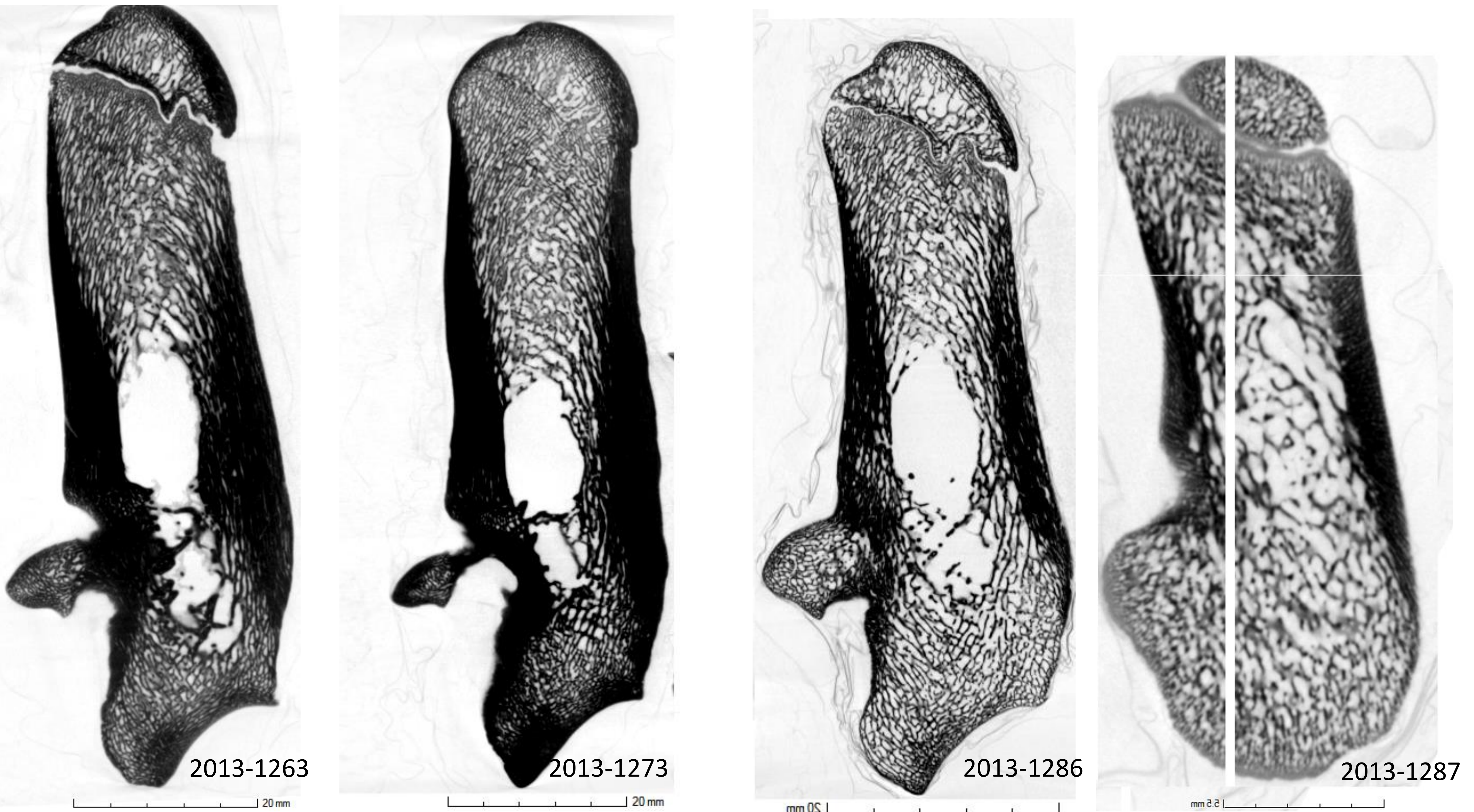
2013-1263
20 mm
2013-1273
20 mm
2013-1286
2013-1287

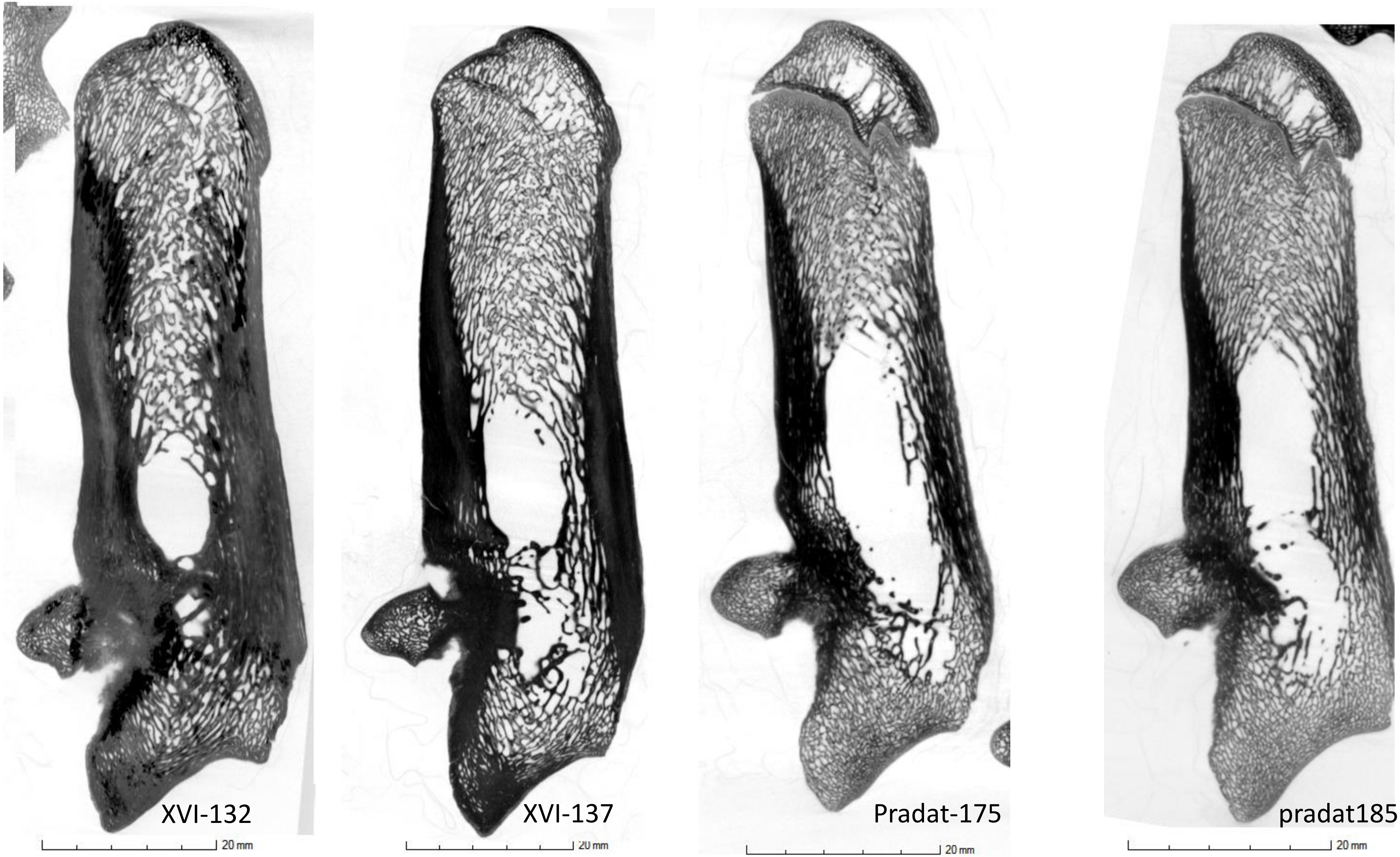
XVI-132
20 mm
XVI-137
20 mm
Pradat-175
20 mm
pradat185
20 mm

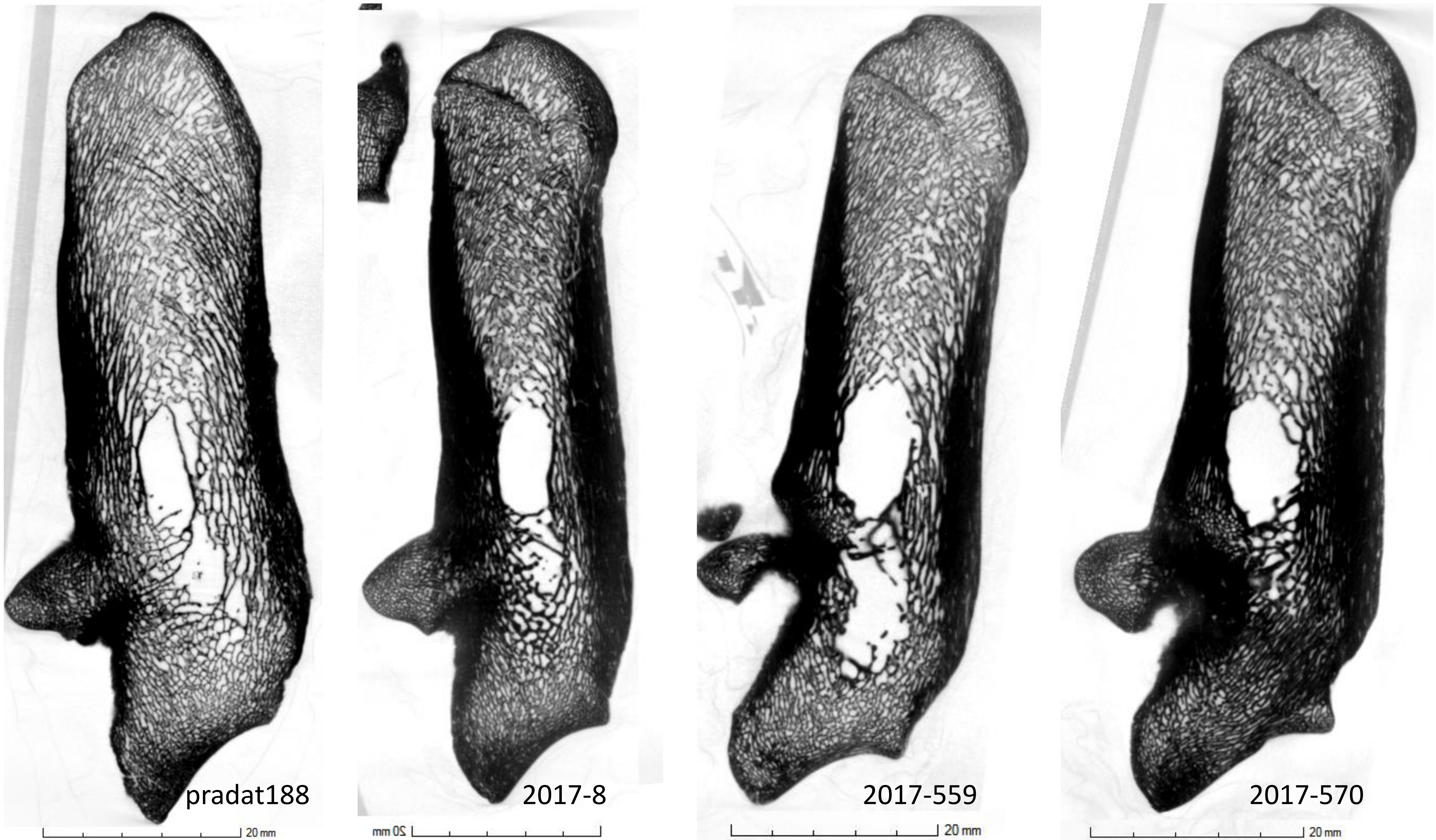
pradat188
20 mm
2017-8
20 mm
2017-559
20 mm
2017-570
20 mm

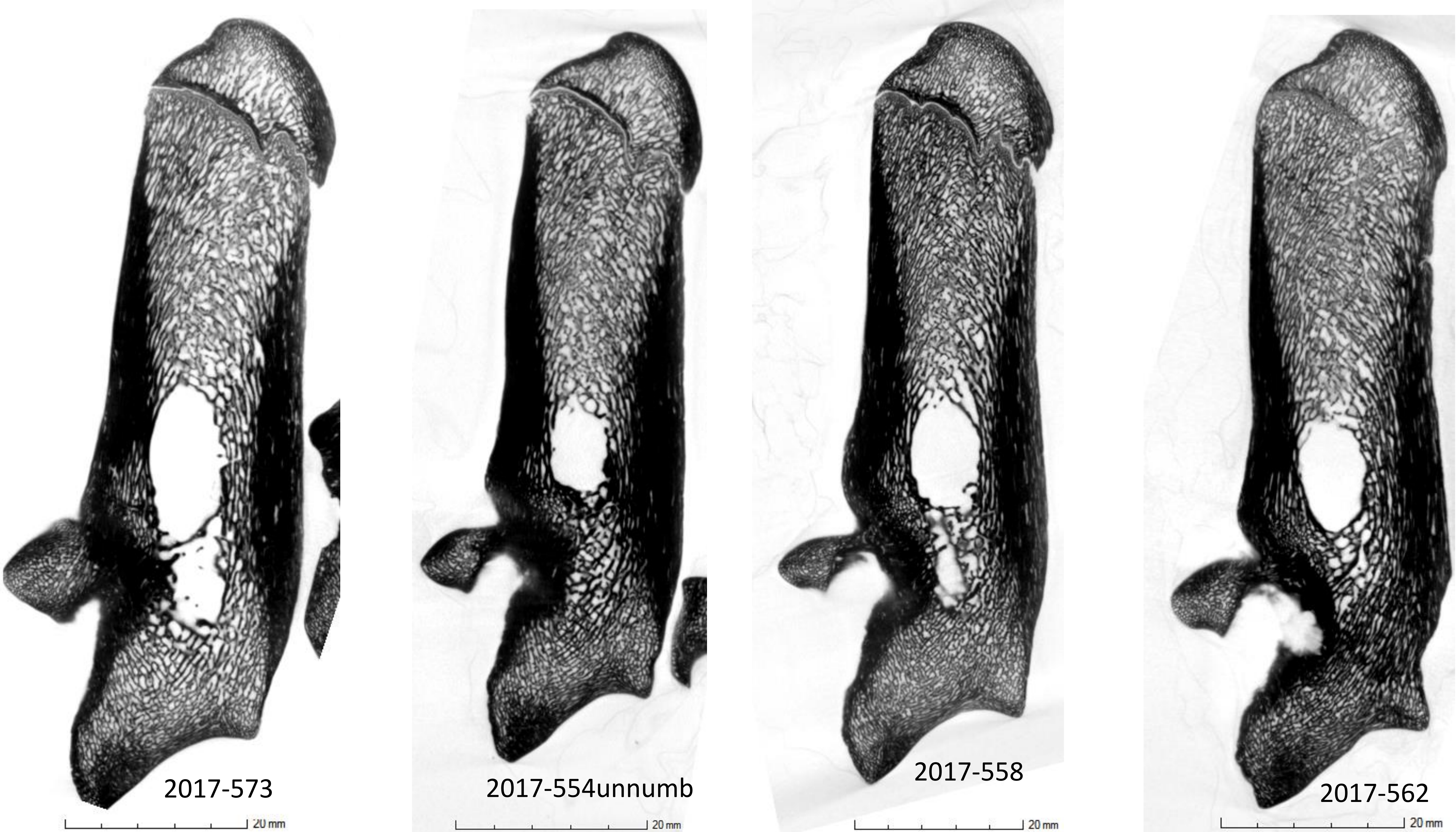
2017-573
20 mm
2017-554unnumb
20 mm
2017-558
20 mm
2017-562
20 mm

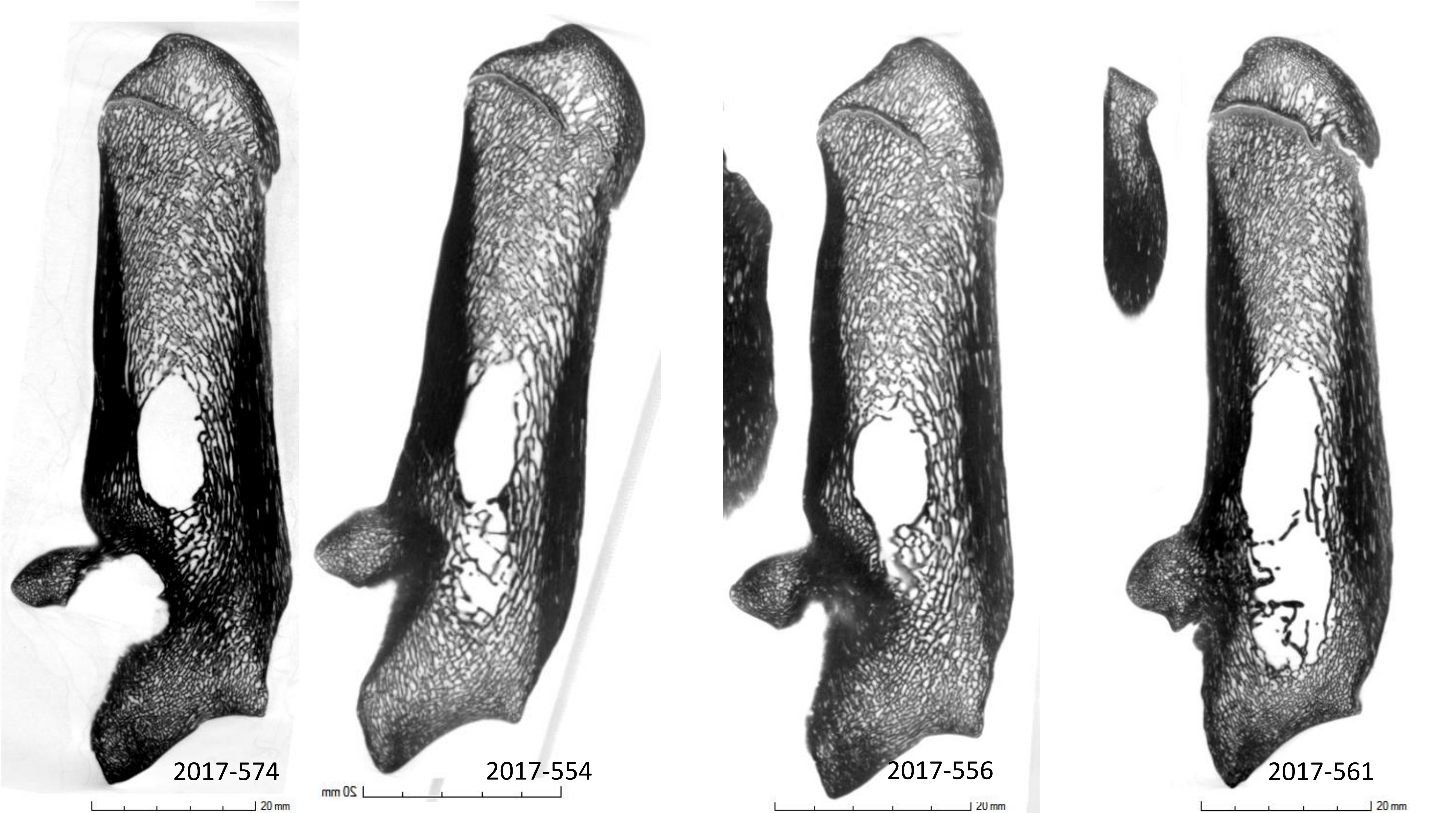
2017-574
20 mm
2017-554
20 mm
2017-556
20 mm
2017-561
20 mm

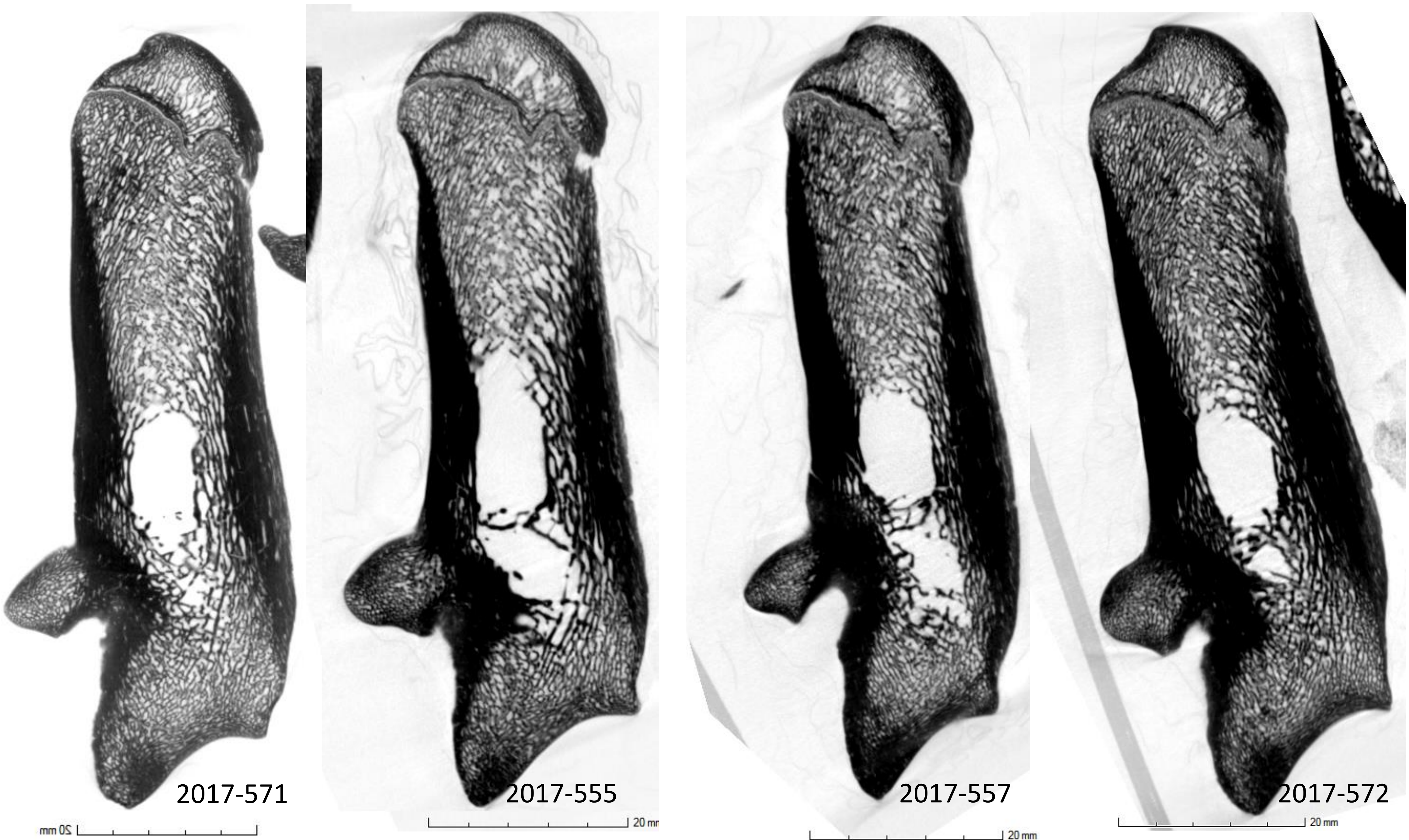

2017-571
2017-555
2017-557
2017-572
20 mm
20 mm

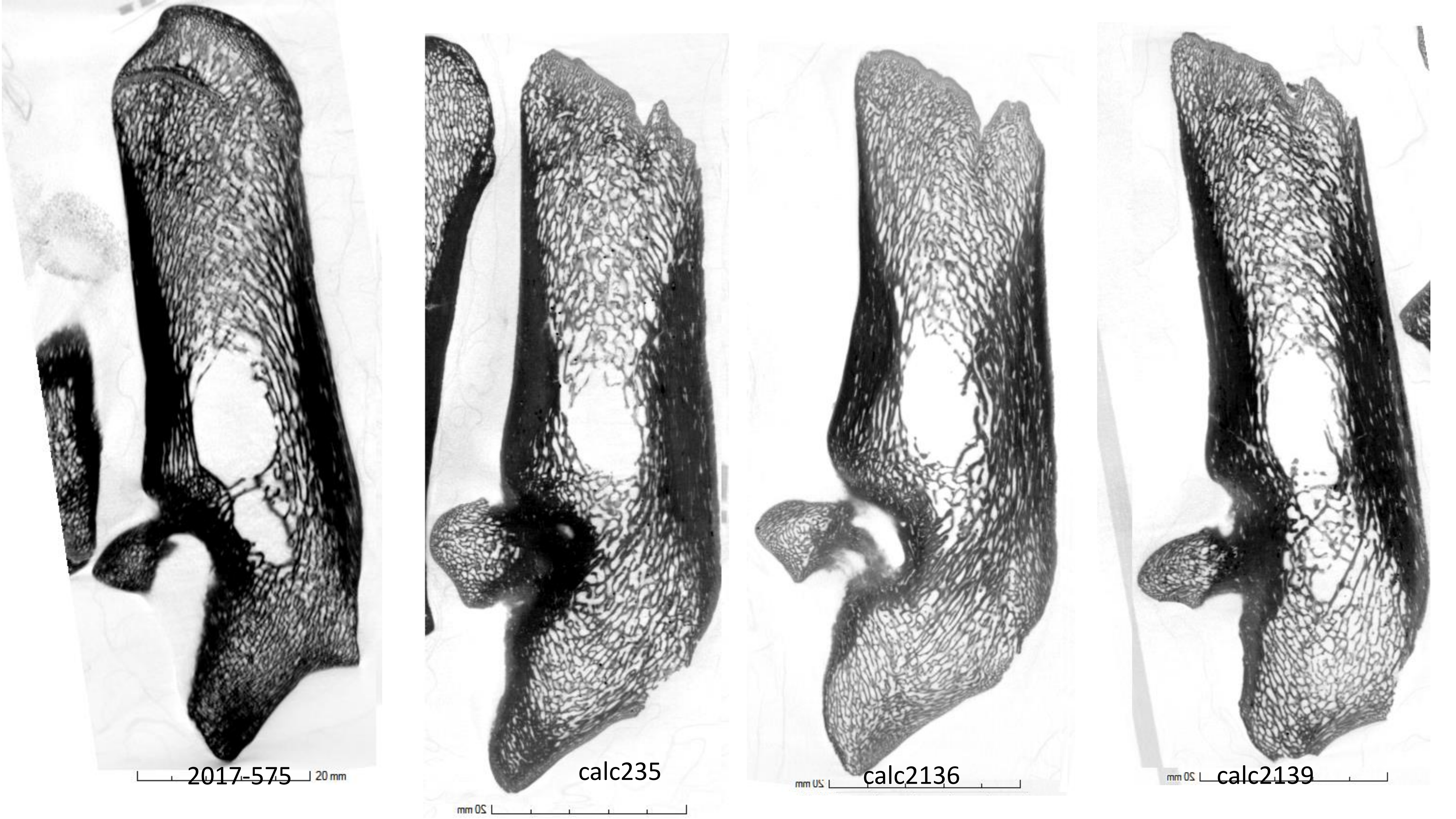
2017-575
20 mm
calc235
calc2136
calc2139

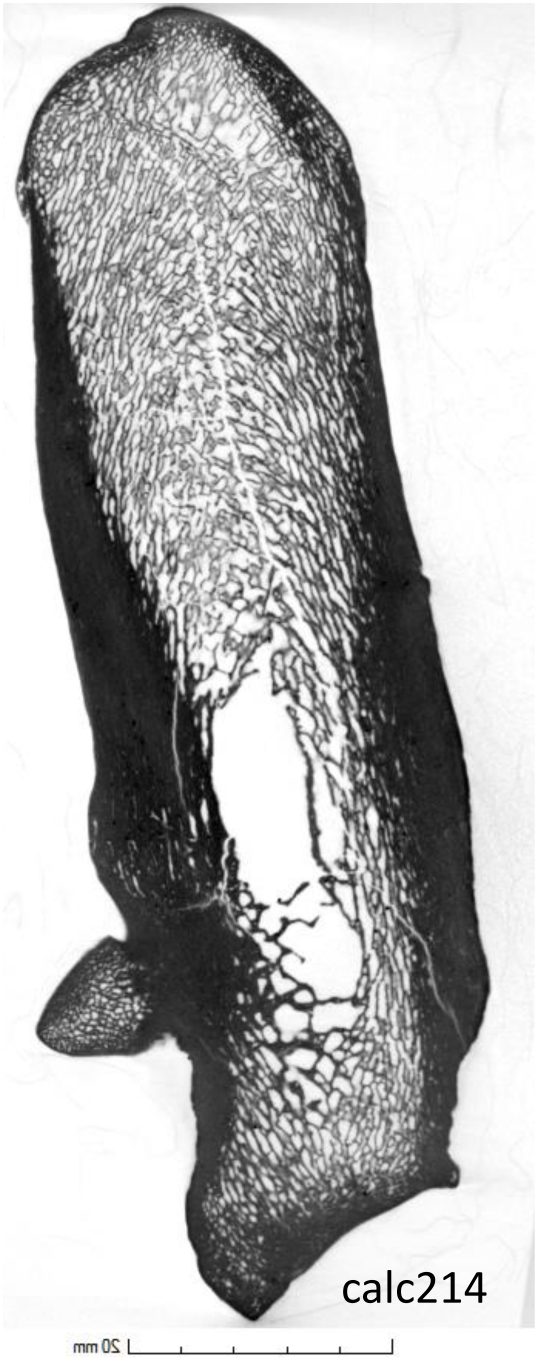
calc214

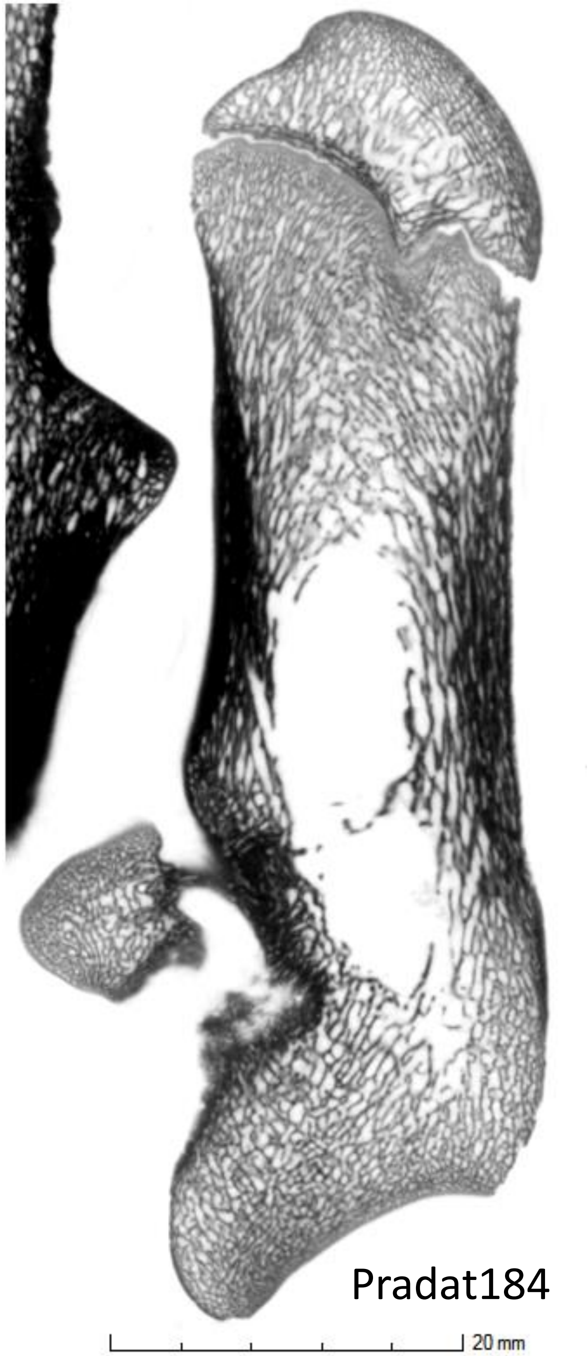
Pradat184
20 mm

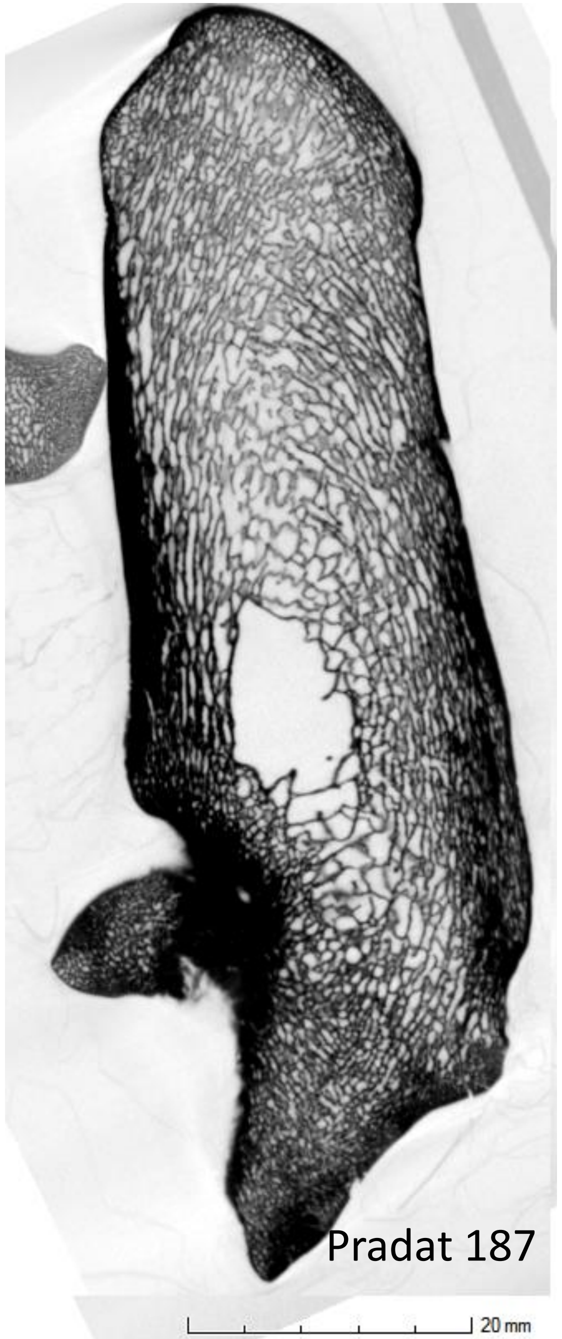
Pradat 187
20 mm
