## Supplementary figures and images for "Can growth in captivity alter the calcaneal microanatomy of a wild ungulate?"

### Appendix 3 transversal sections

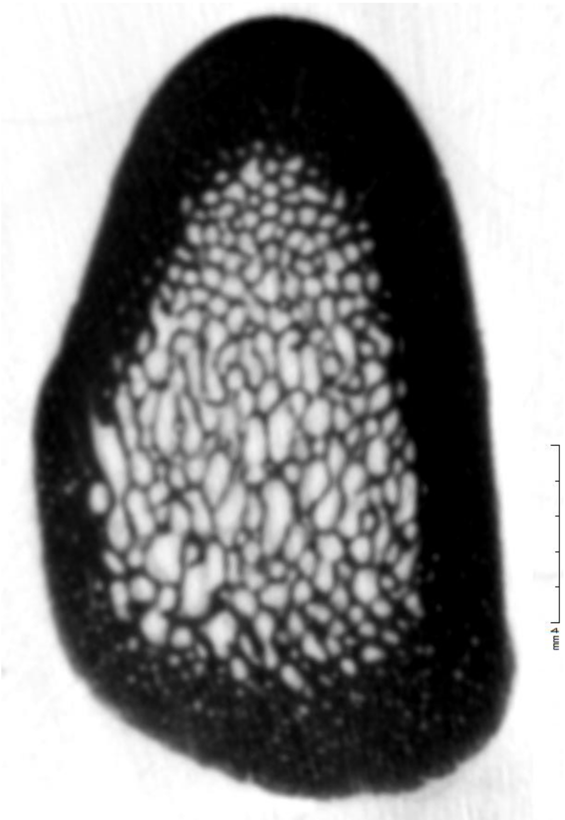

2017-578

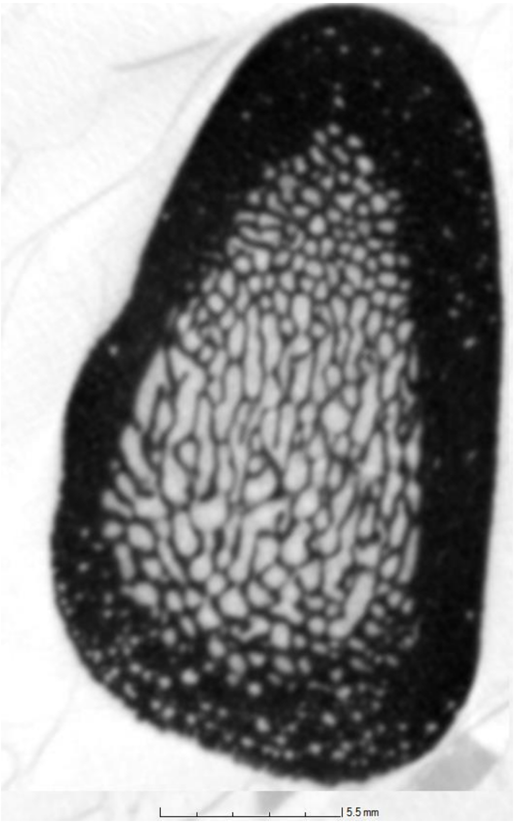


2017-580

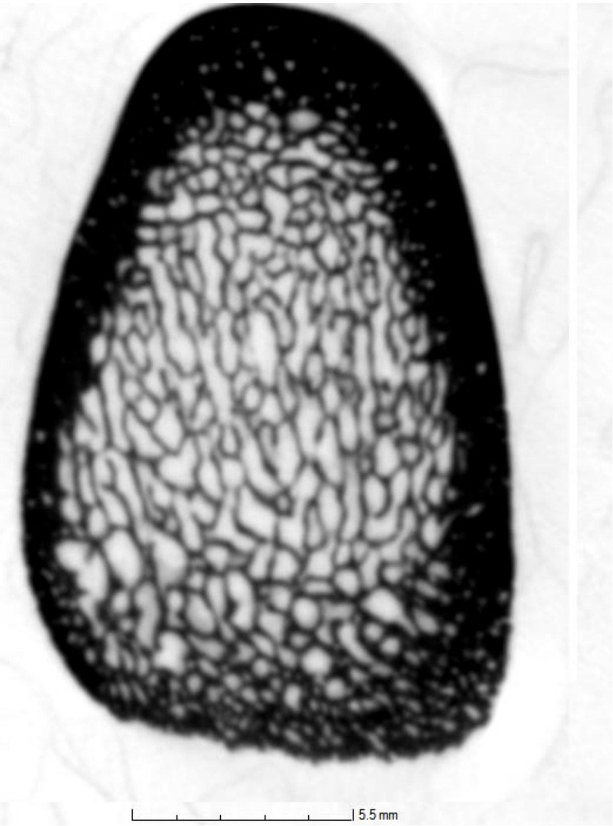


2017-576

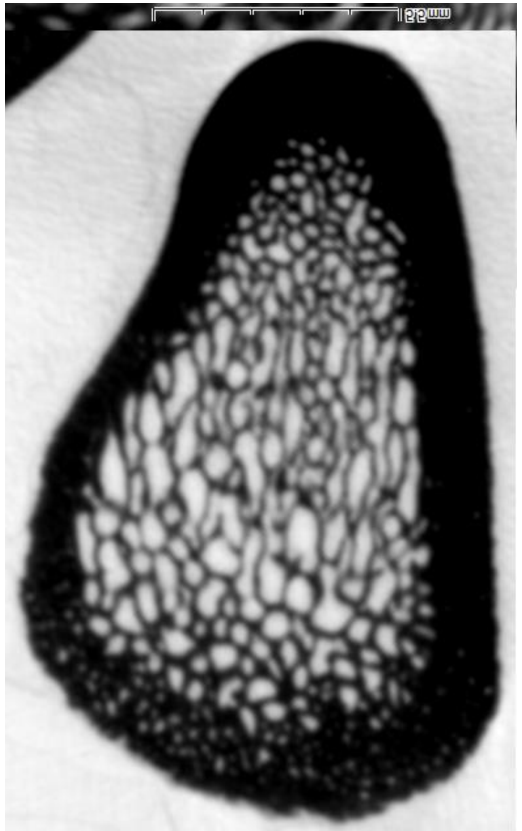

2017-581

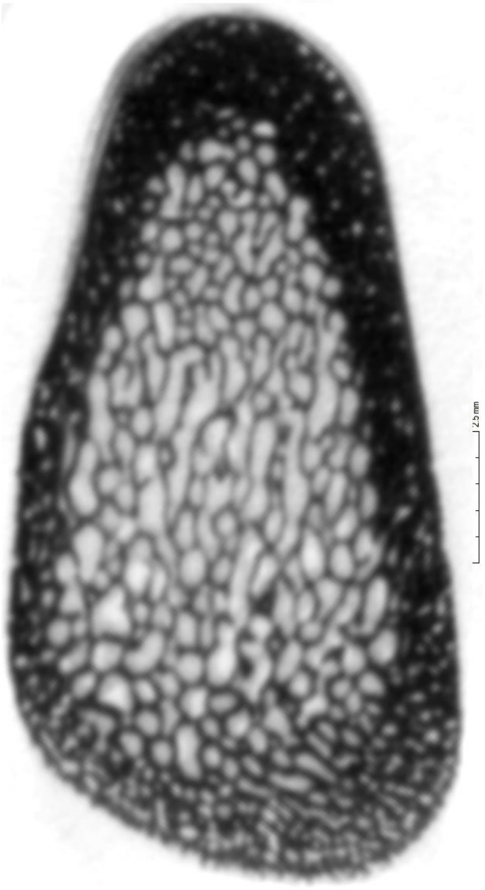

2017-569

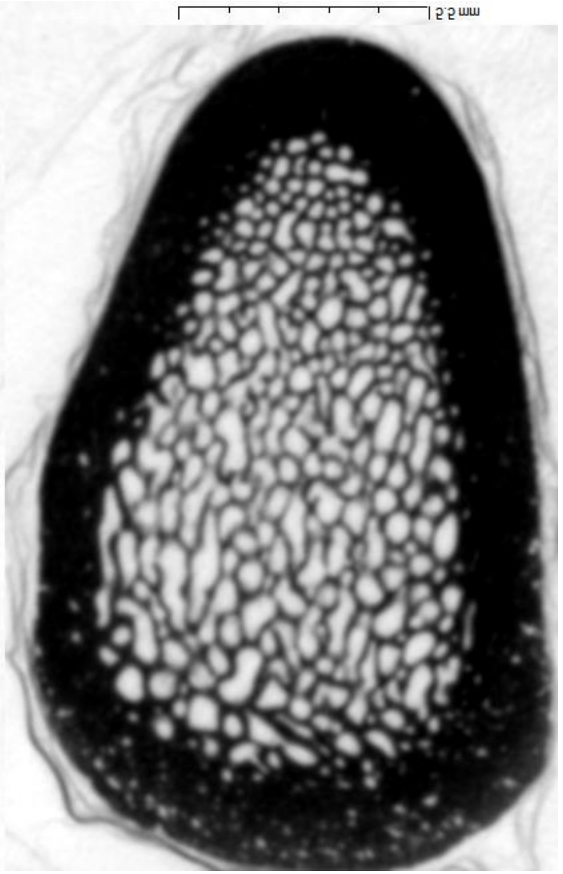

2017-579

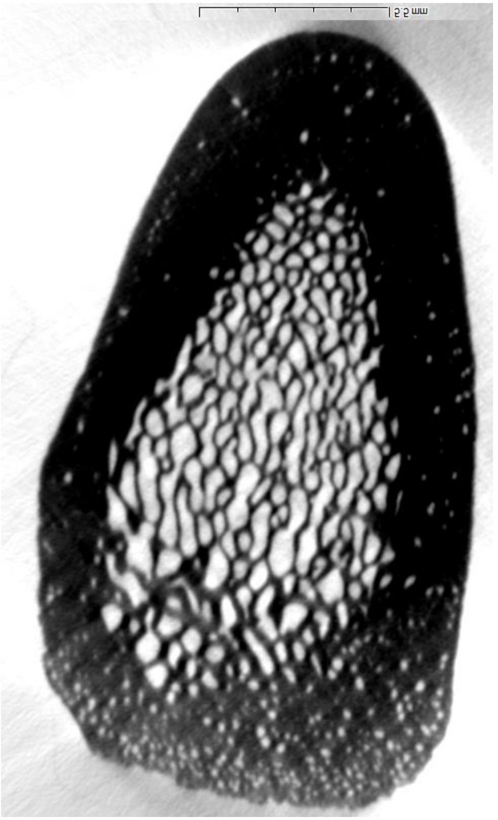

2017-560

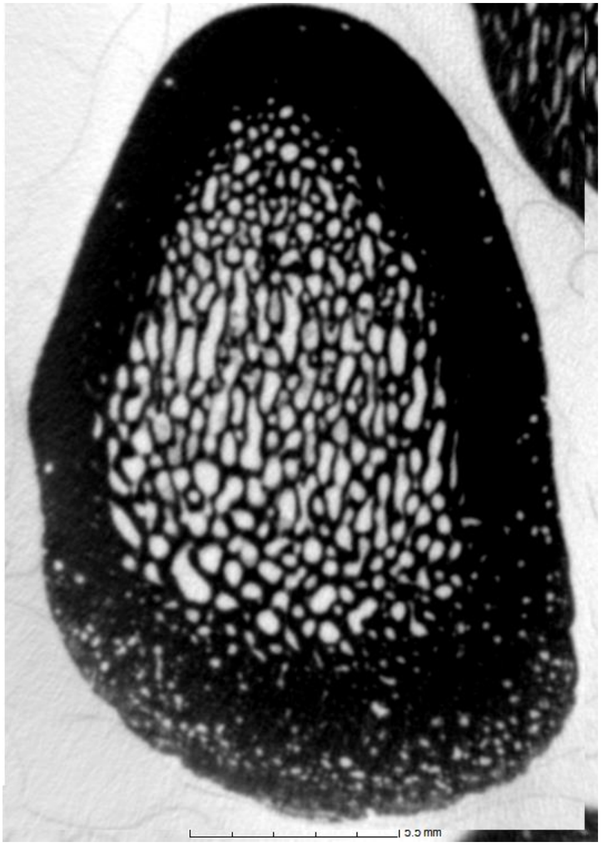

2017-563

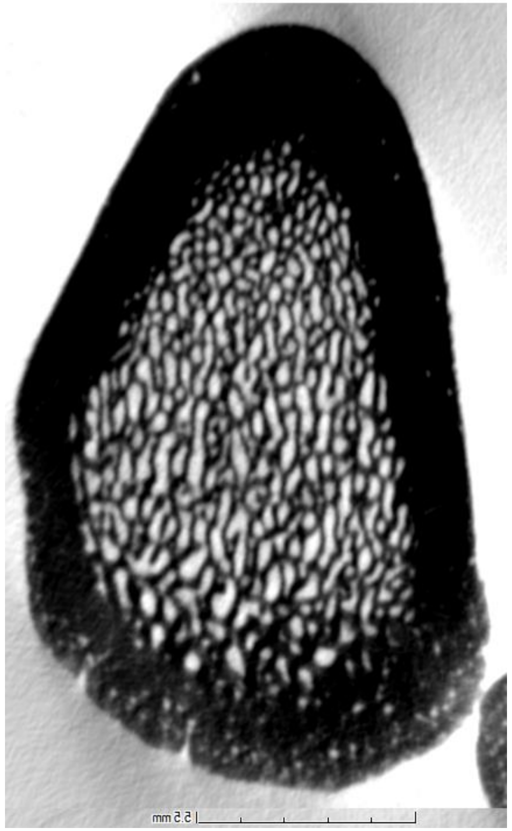

2017-564

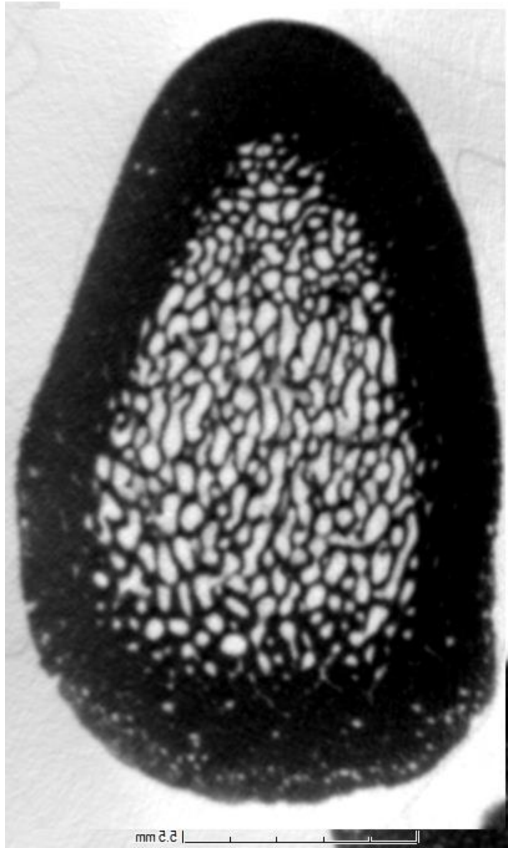

2017-568

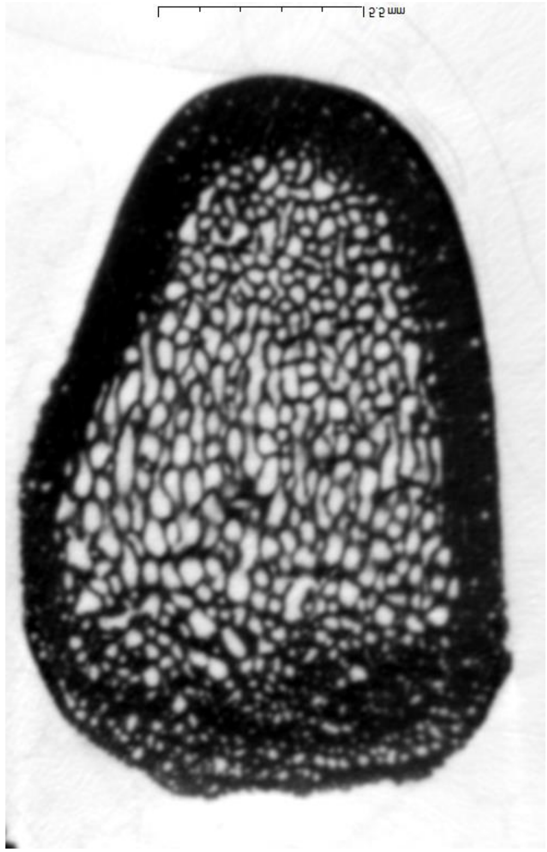

2013-1264

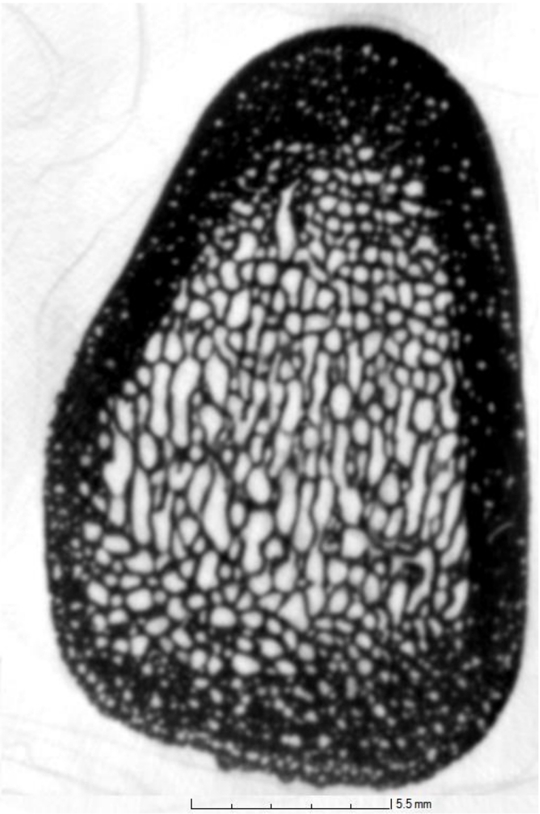


2013-1270

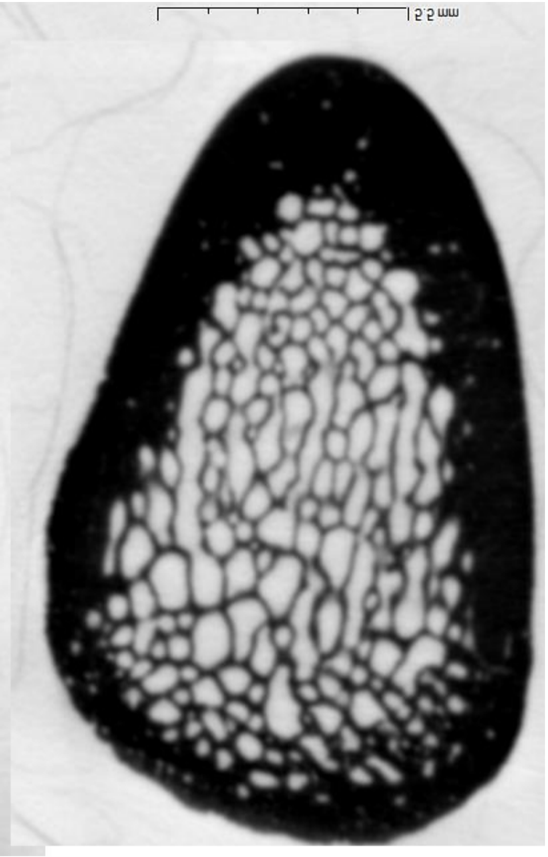

2013-1272

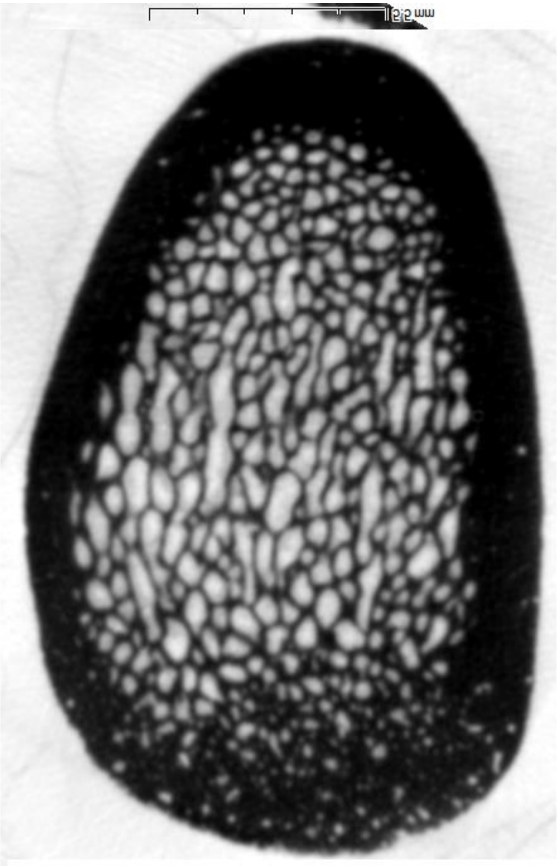

2013-1285

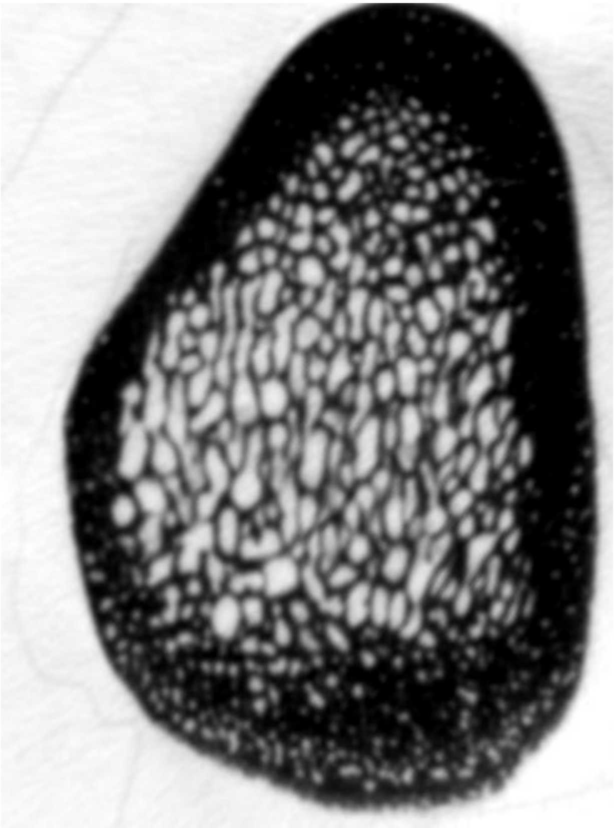

2013-1257

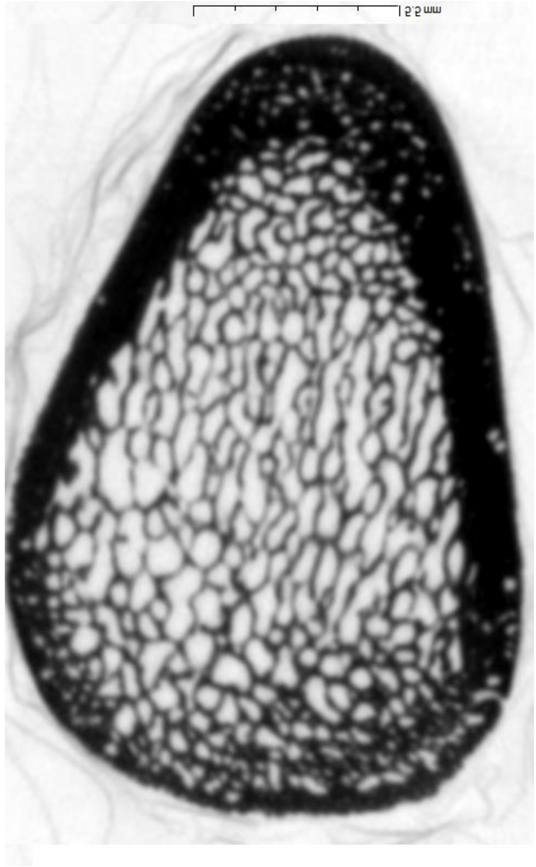

2013-1258

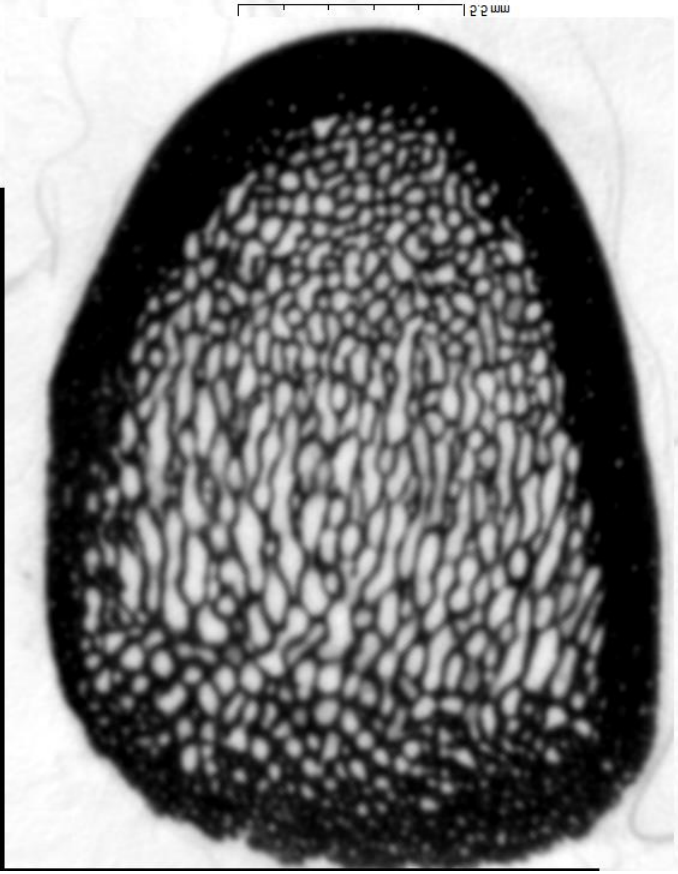

2013-1263

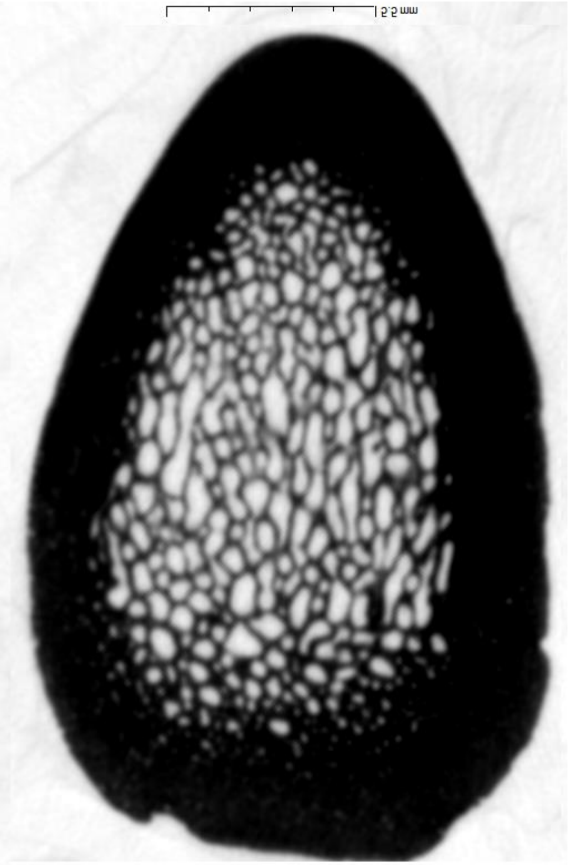

2013-1273

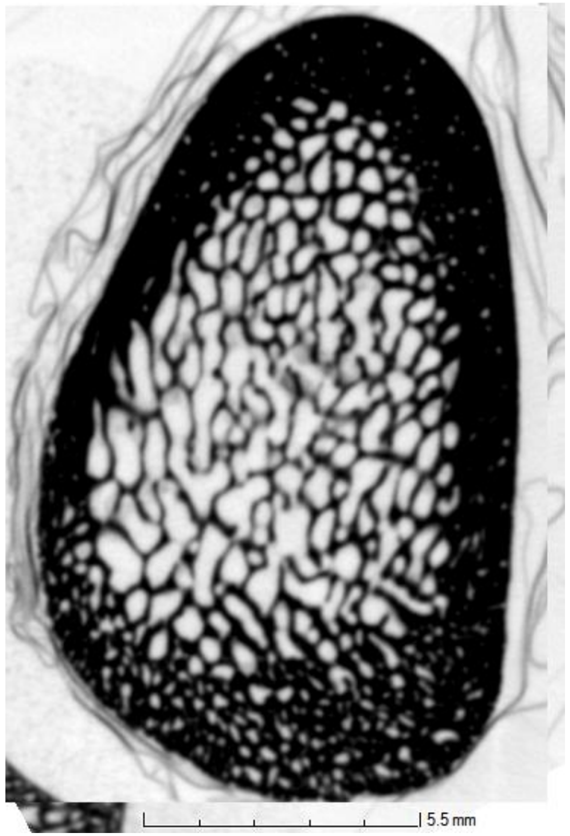


2013-1286

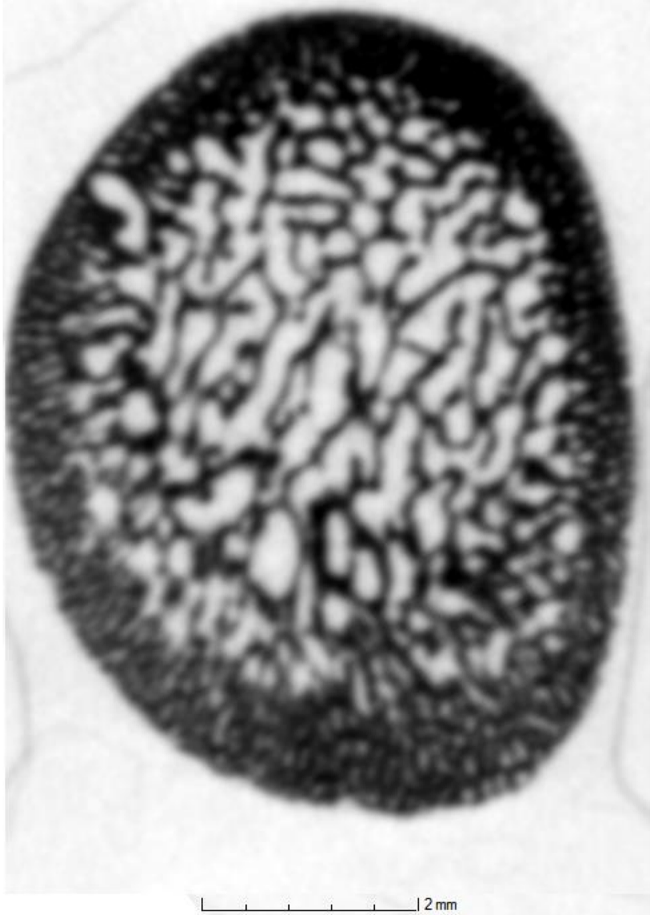


2013-1287

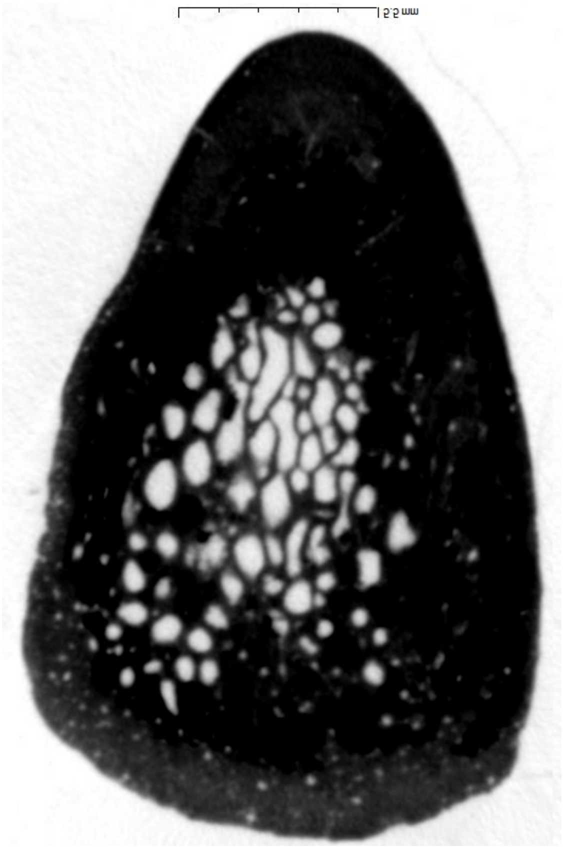

XVI-132

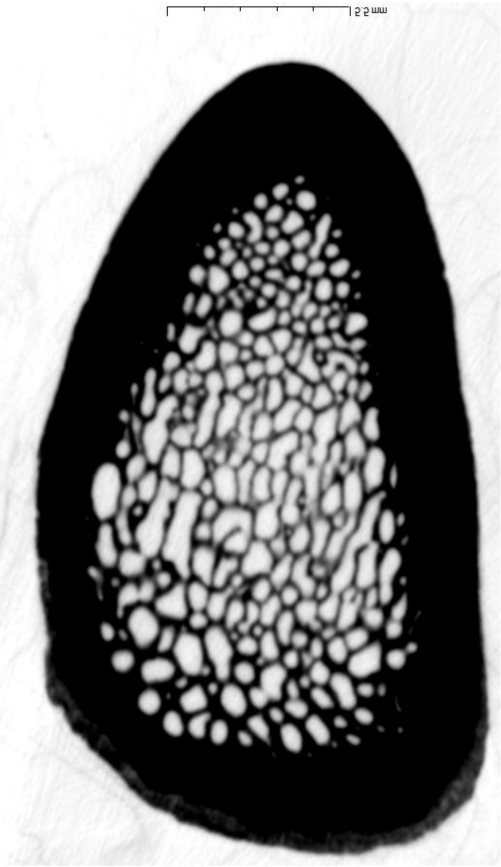

XVI-137

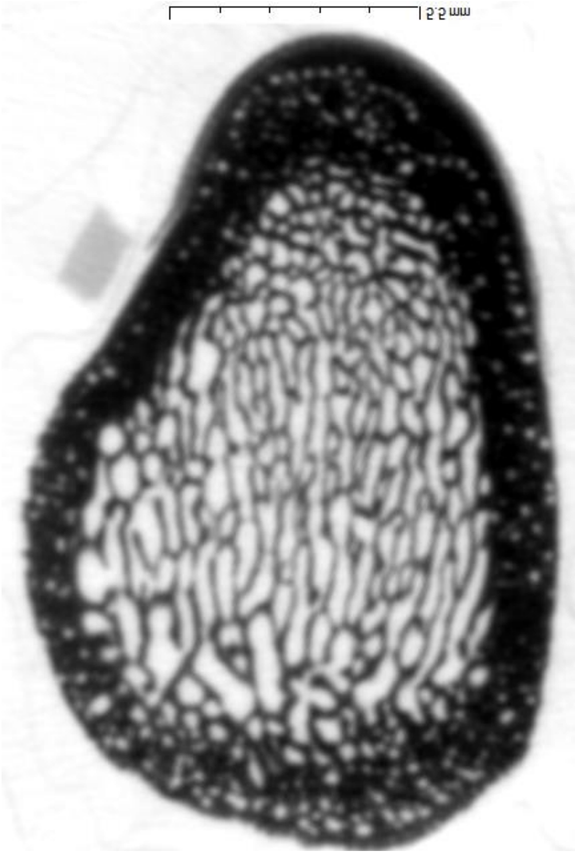

Pradat-175

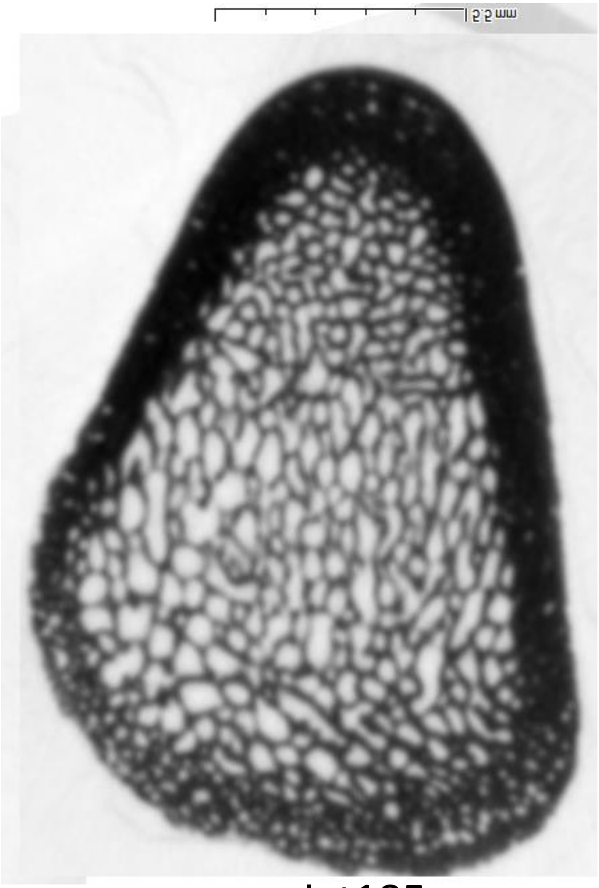

pradat185

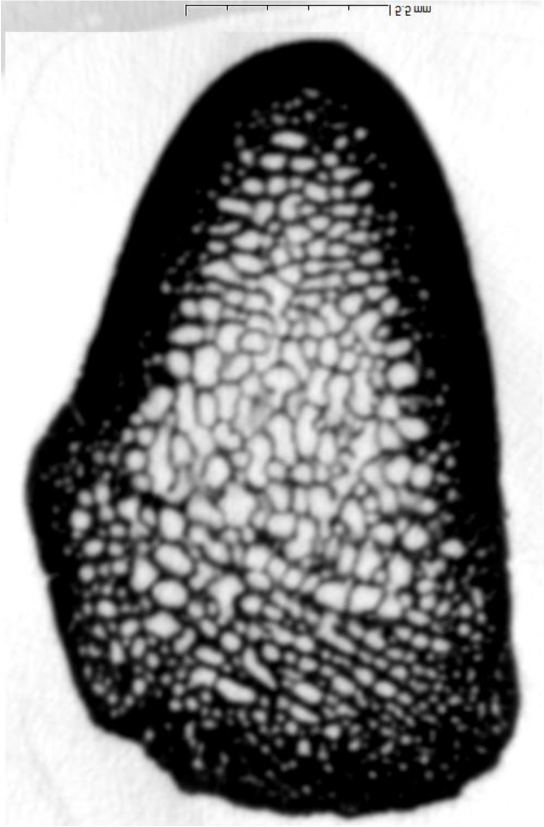

pradat188

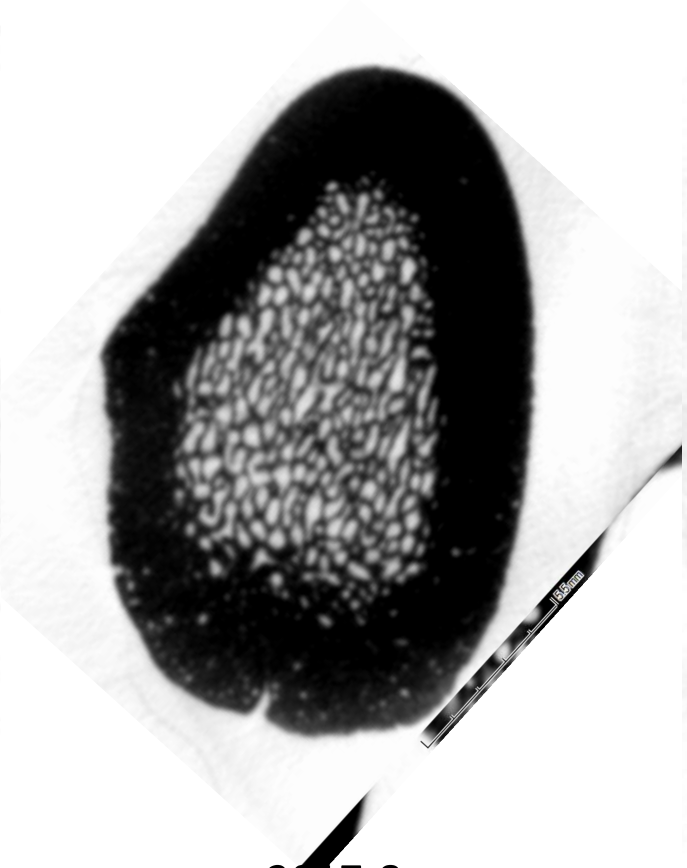

2017-8

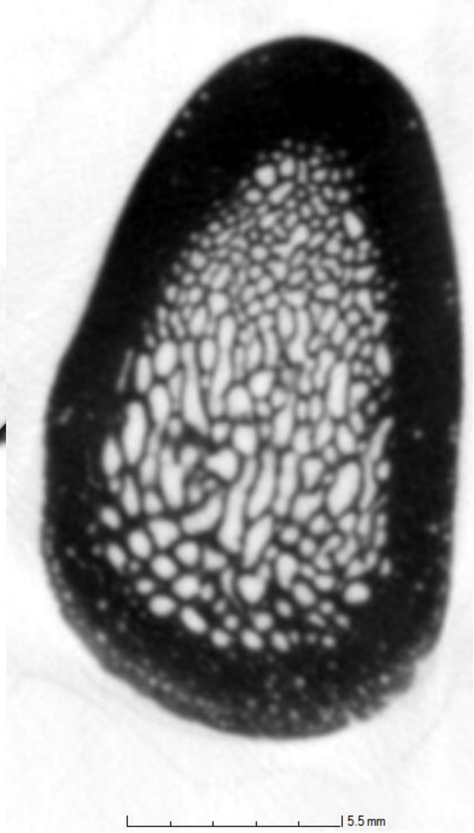


2017-559

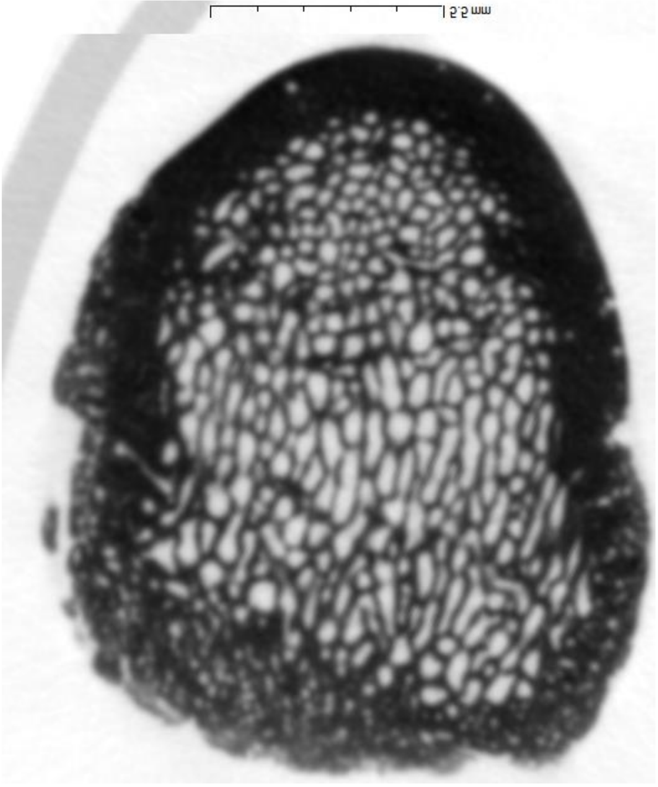

2017-570

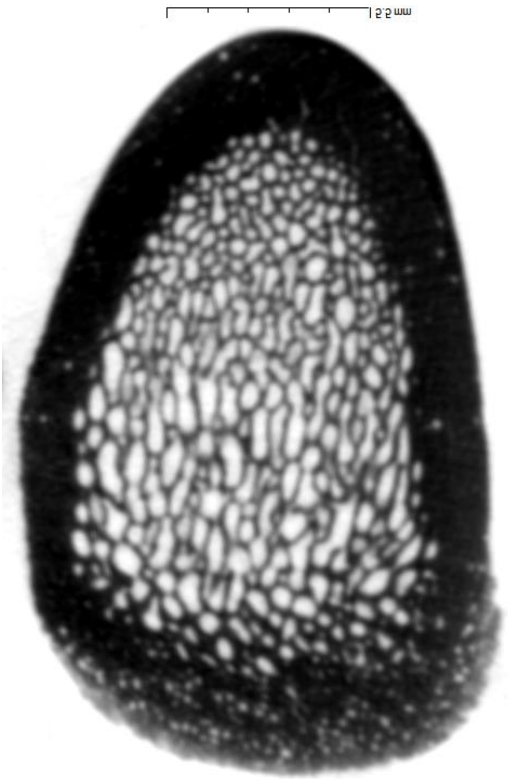

2017-573

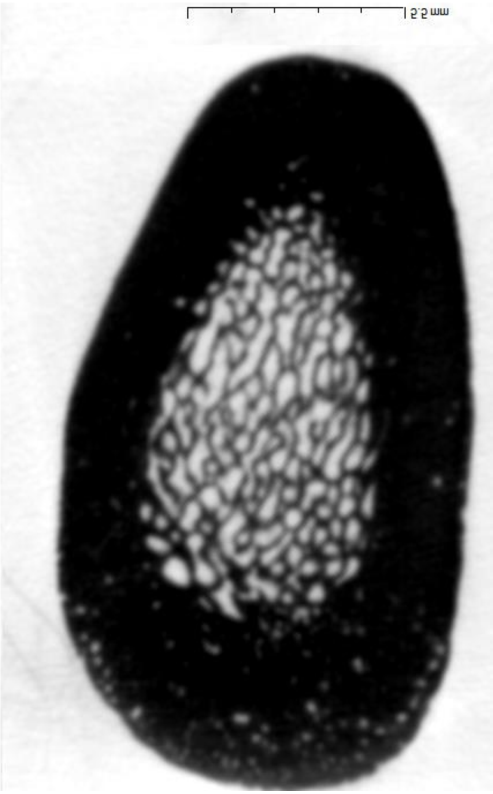

2017-554unnumb

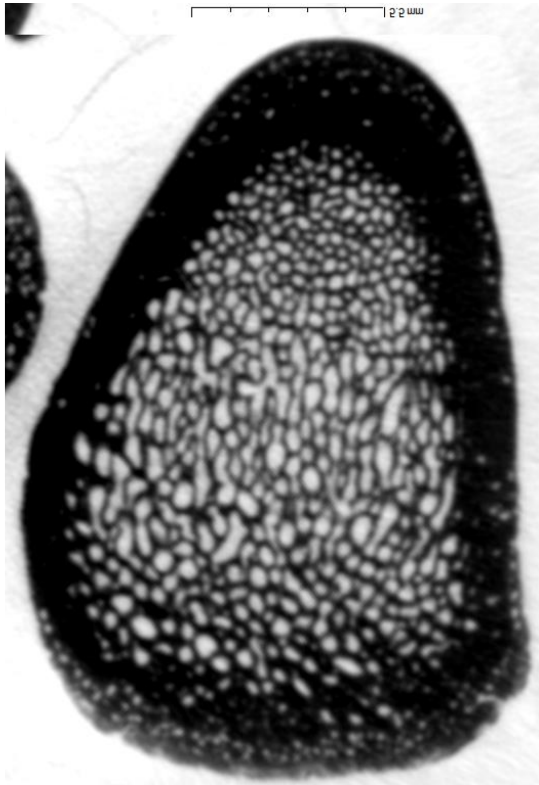

2017-558

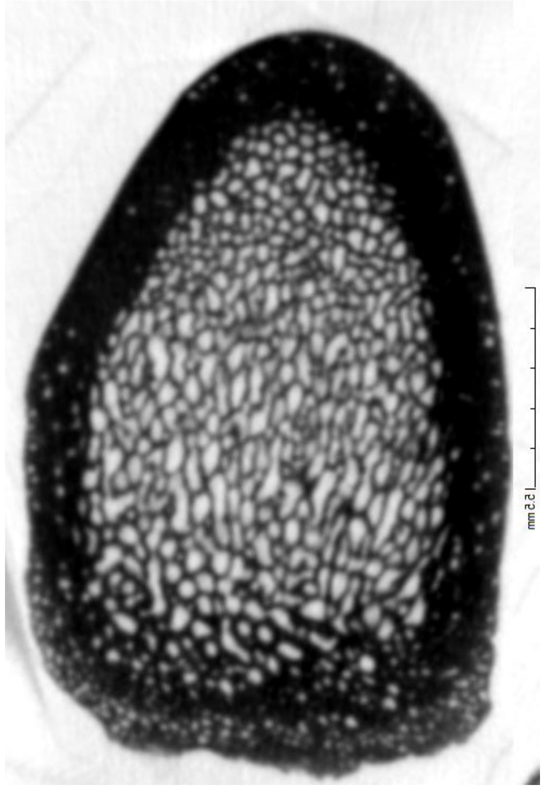

2017-562

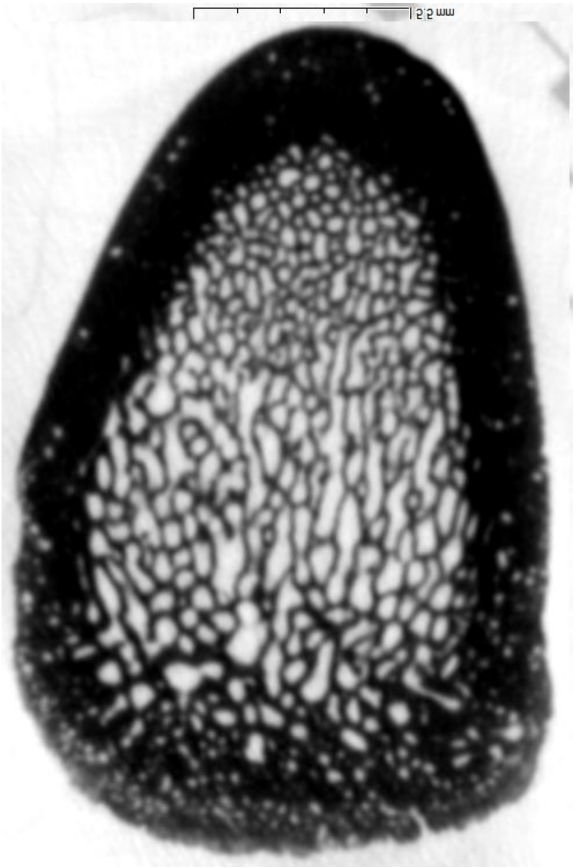

2017-574

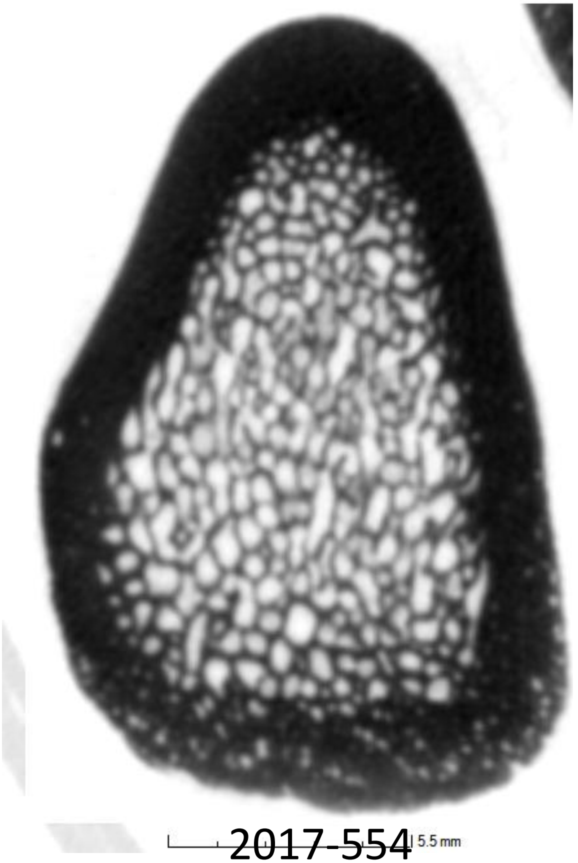

2017-554

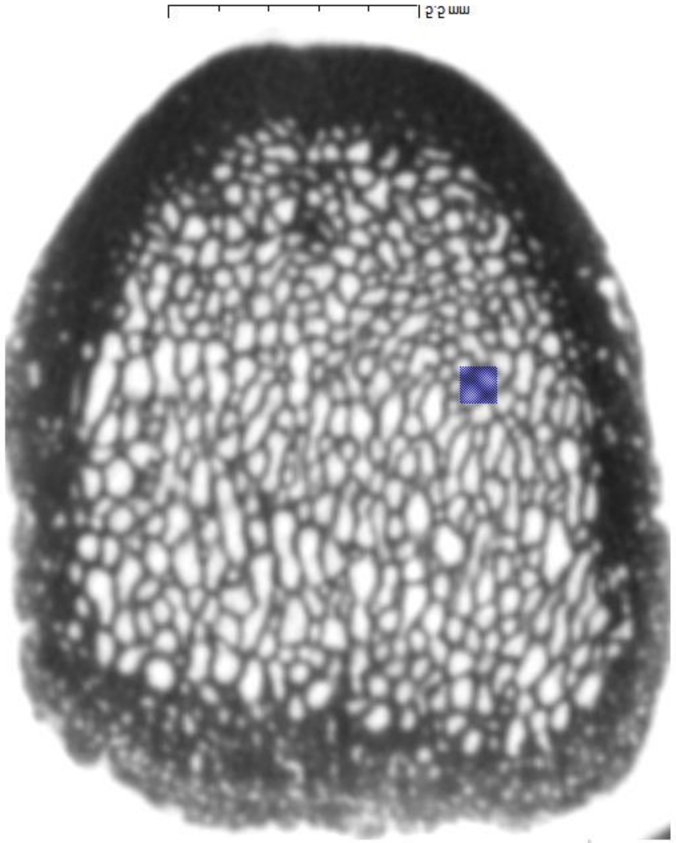

2017-556

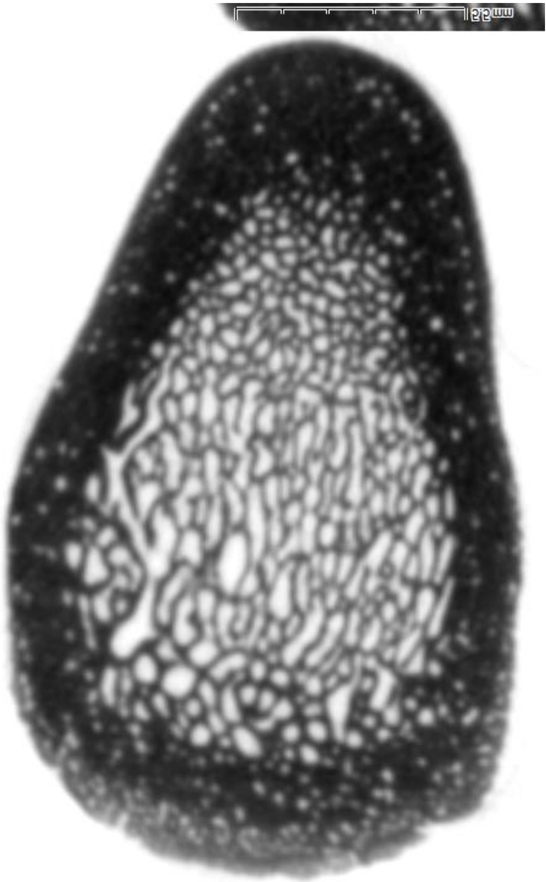

2017-561

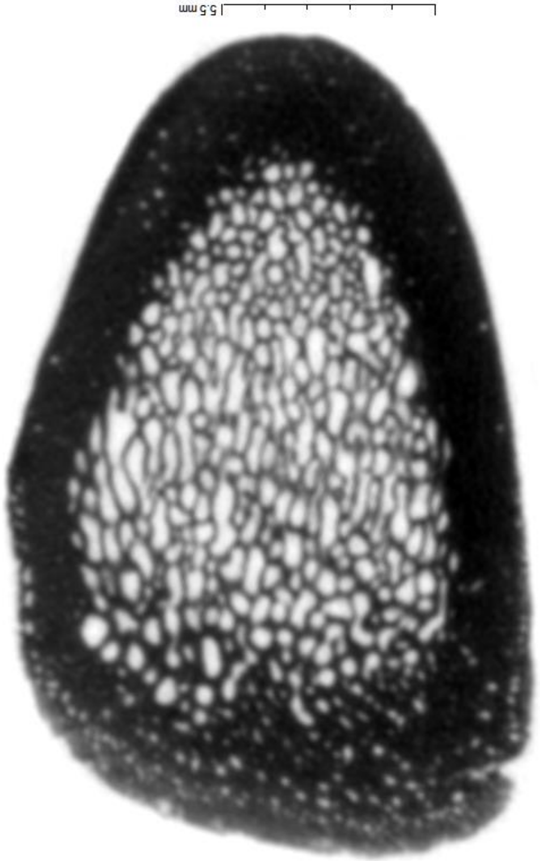

2017-571

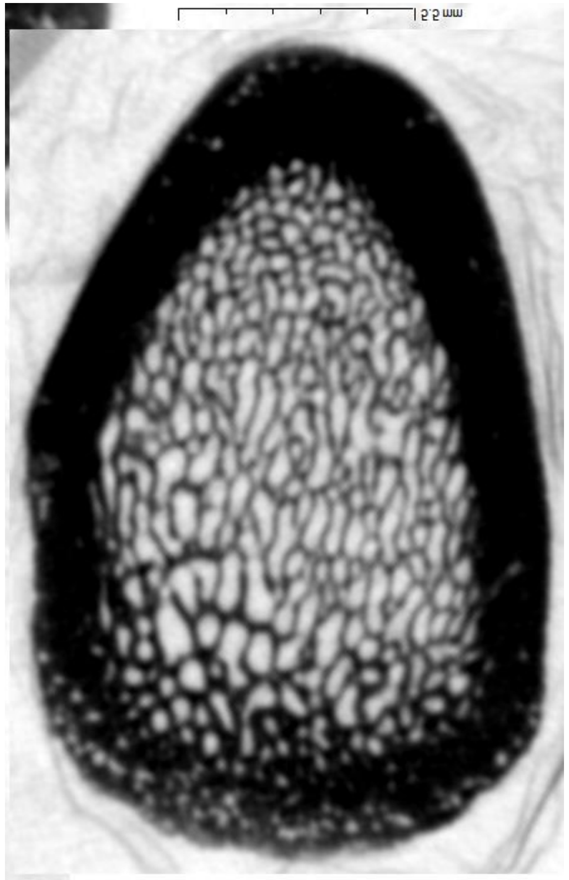


2017-555

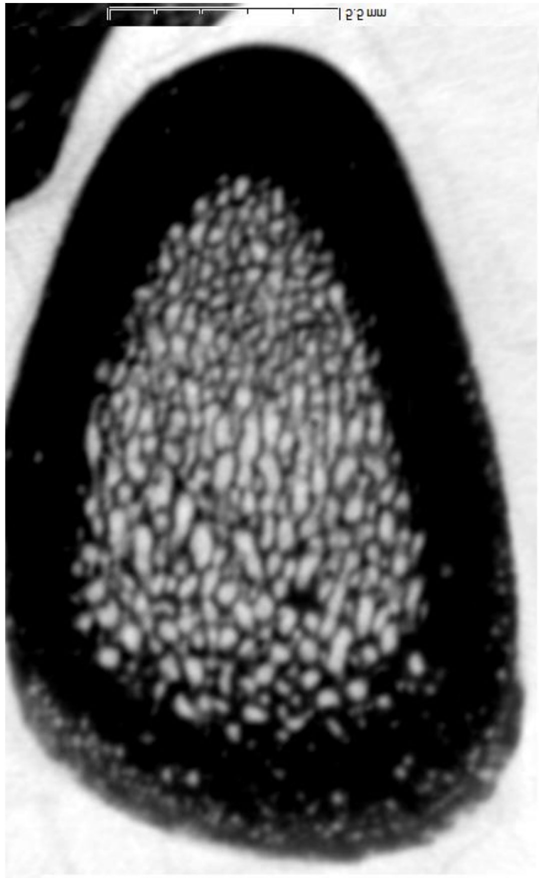


2017-557

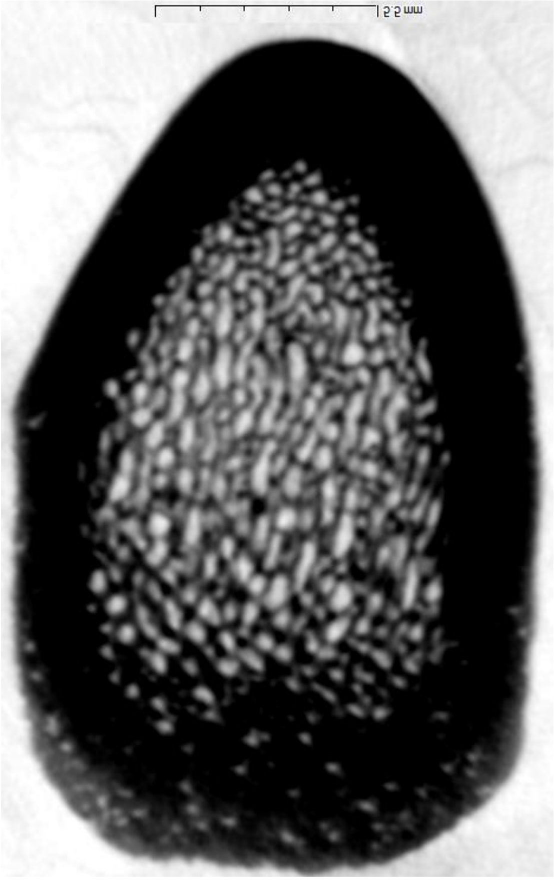


2017-572

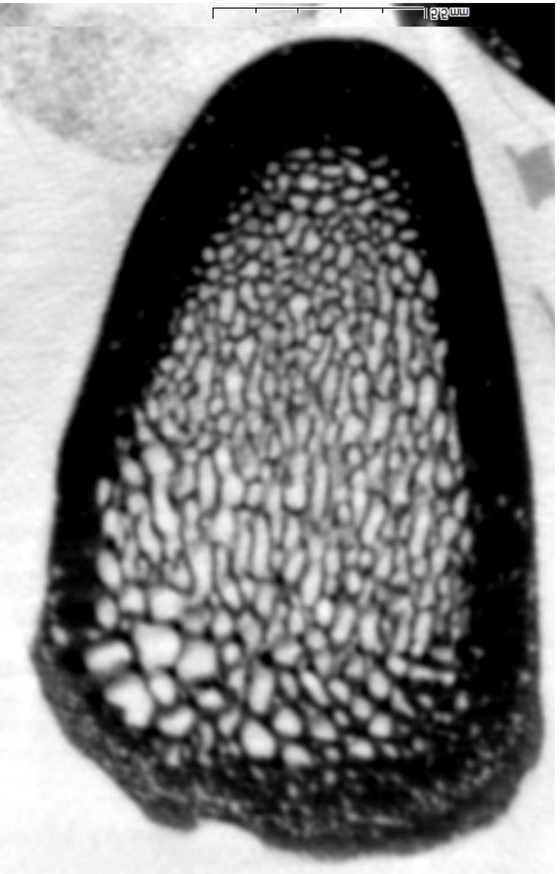

2017-575

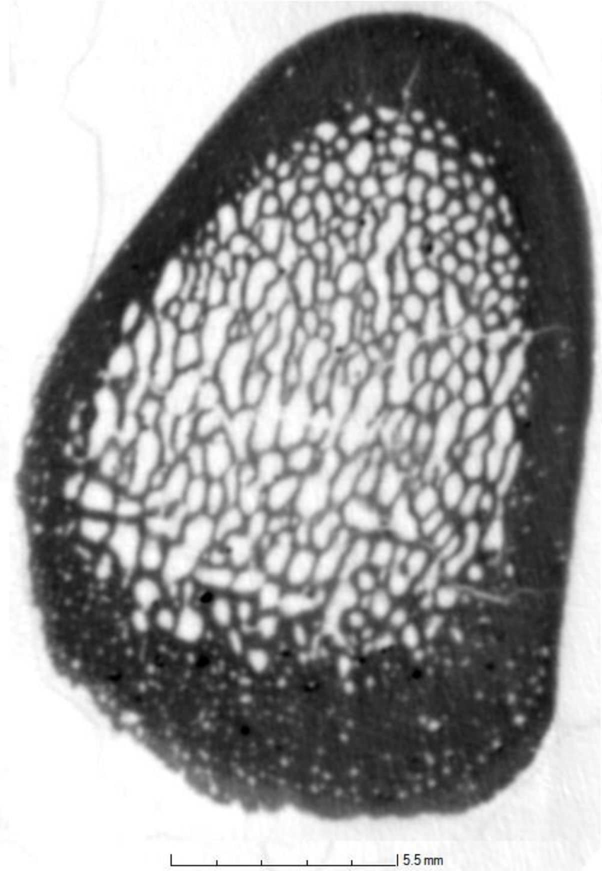


Calc 235

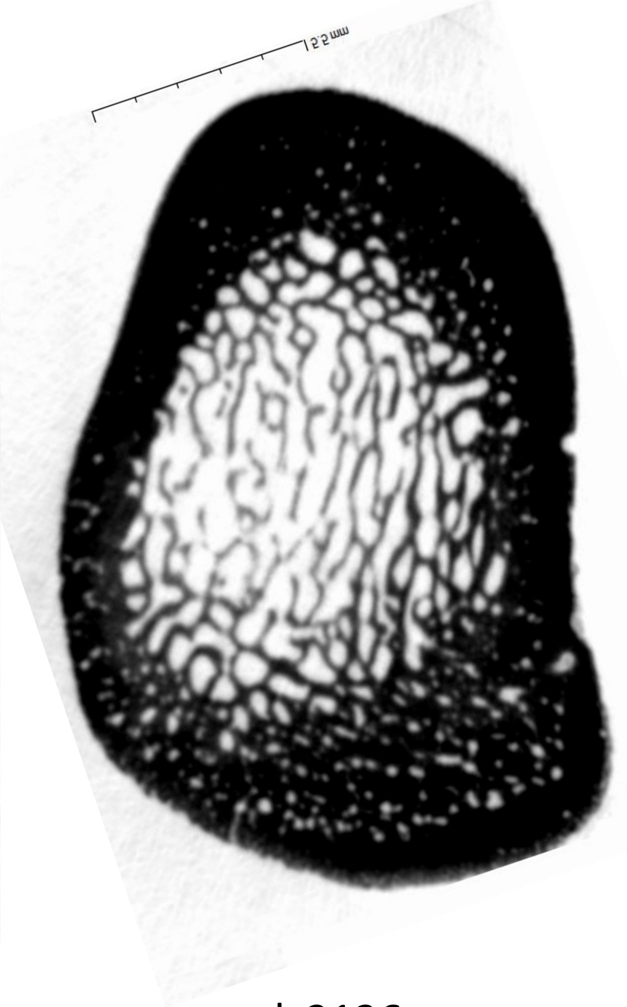


calc2136

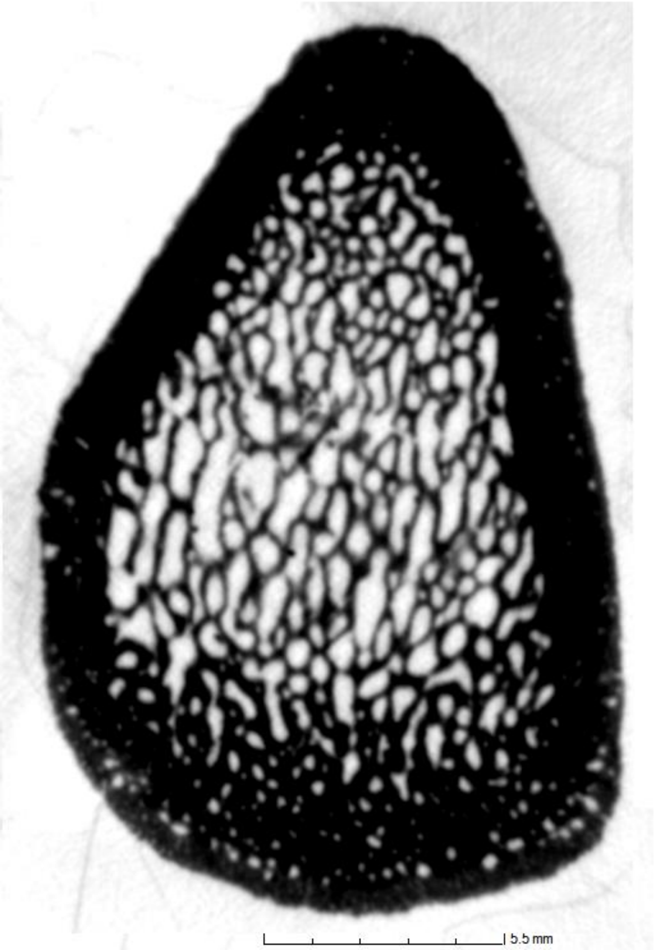


calc2139

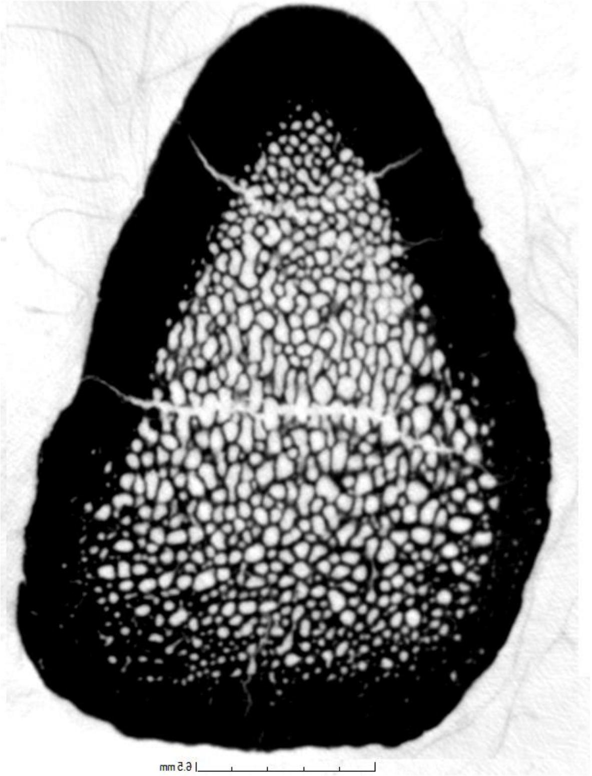

Calc214

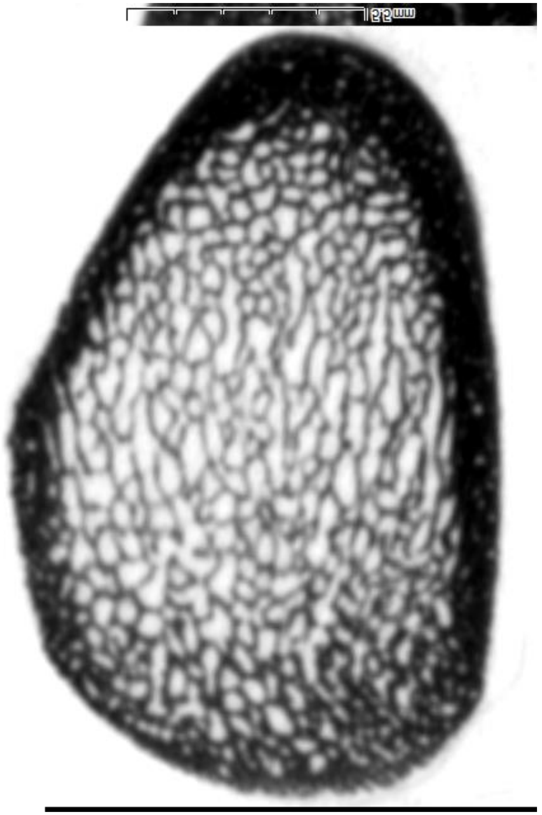

Pradat184

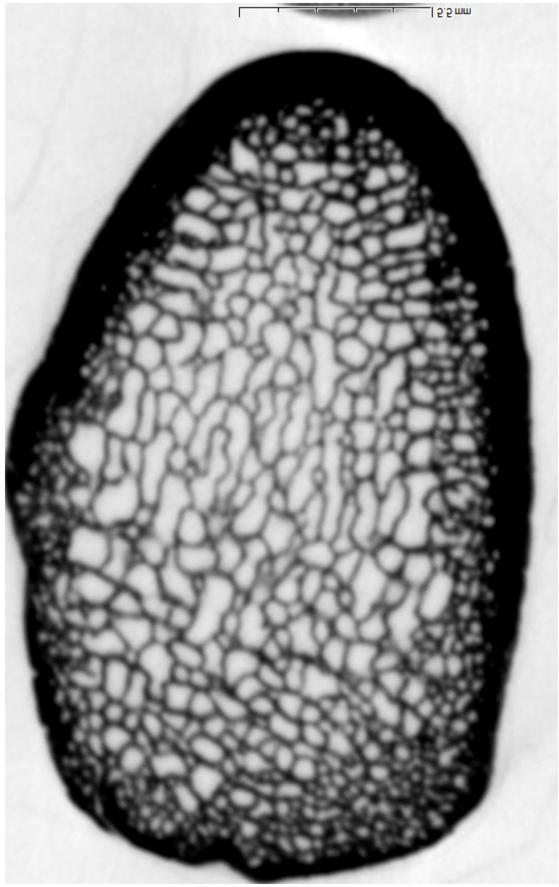

Pradat 187
