## Appendix 4 3D mapps of the cortical thickness for "Can growth in captivity alter the calcaneal microanatomy of a wild ungulate?"

2017-578

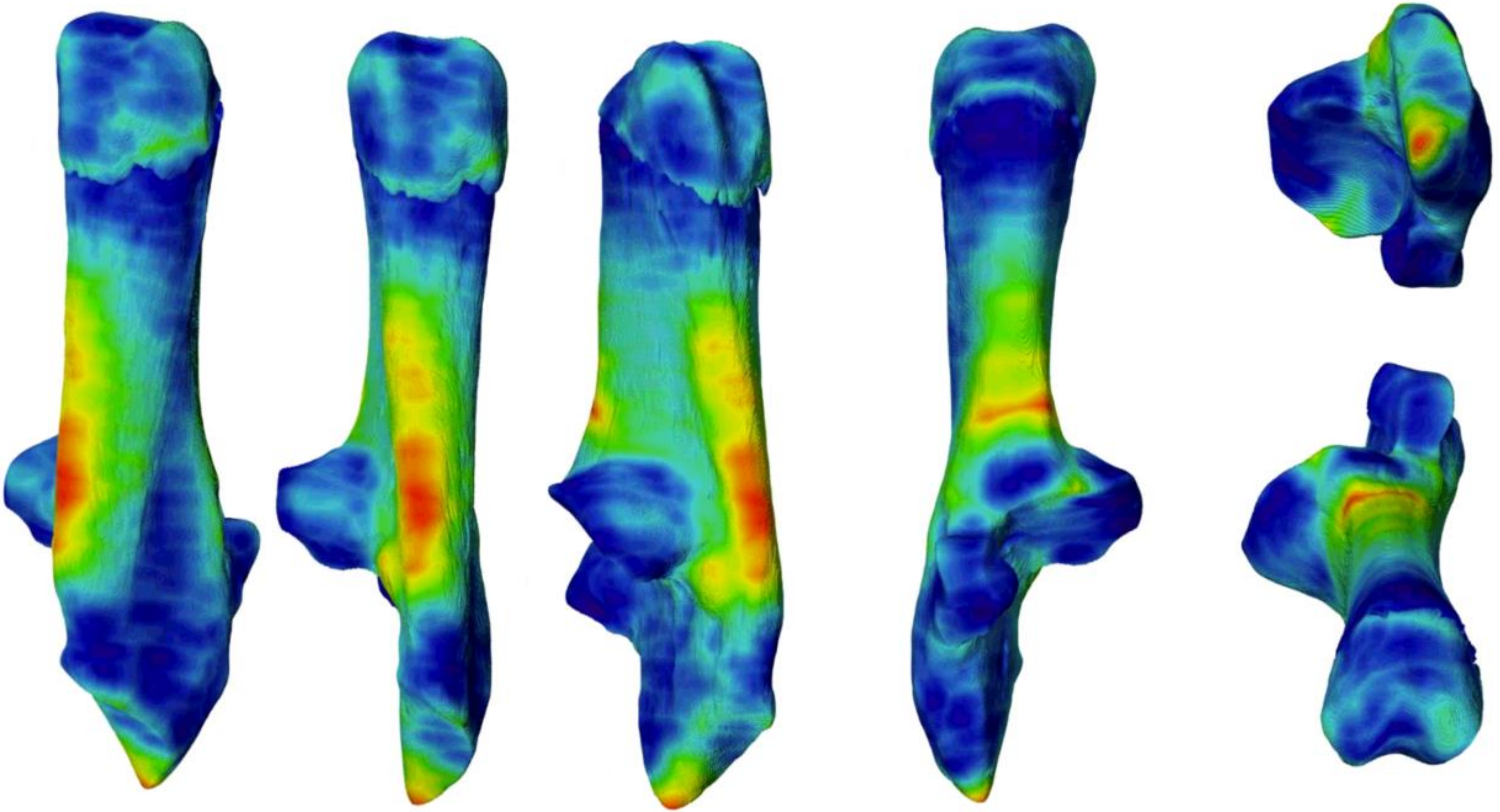

2017-580

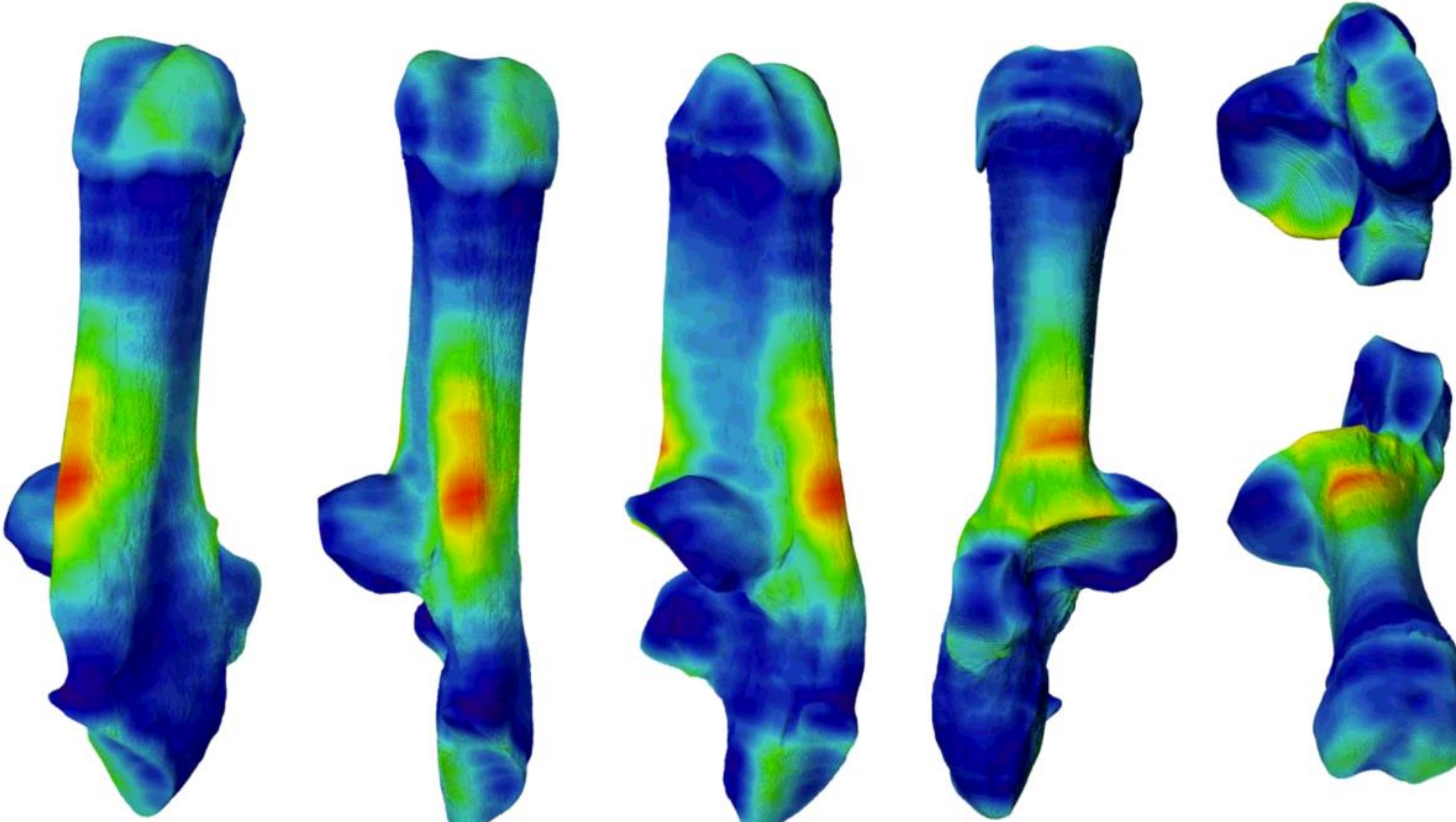

2017-576

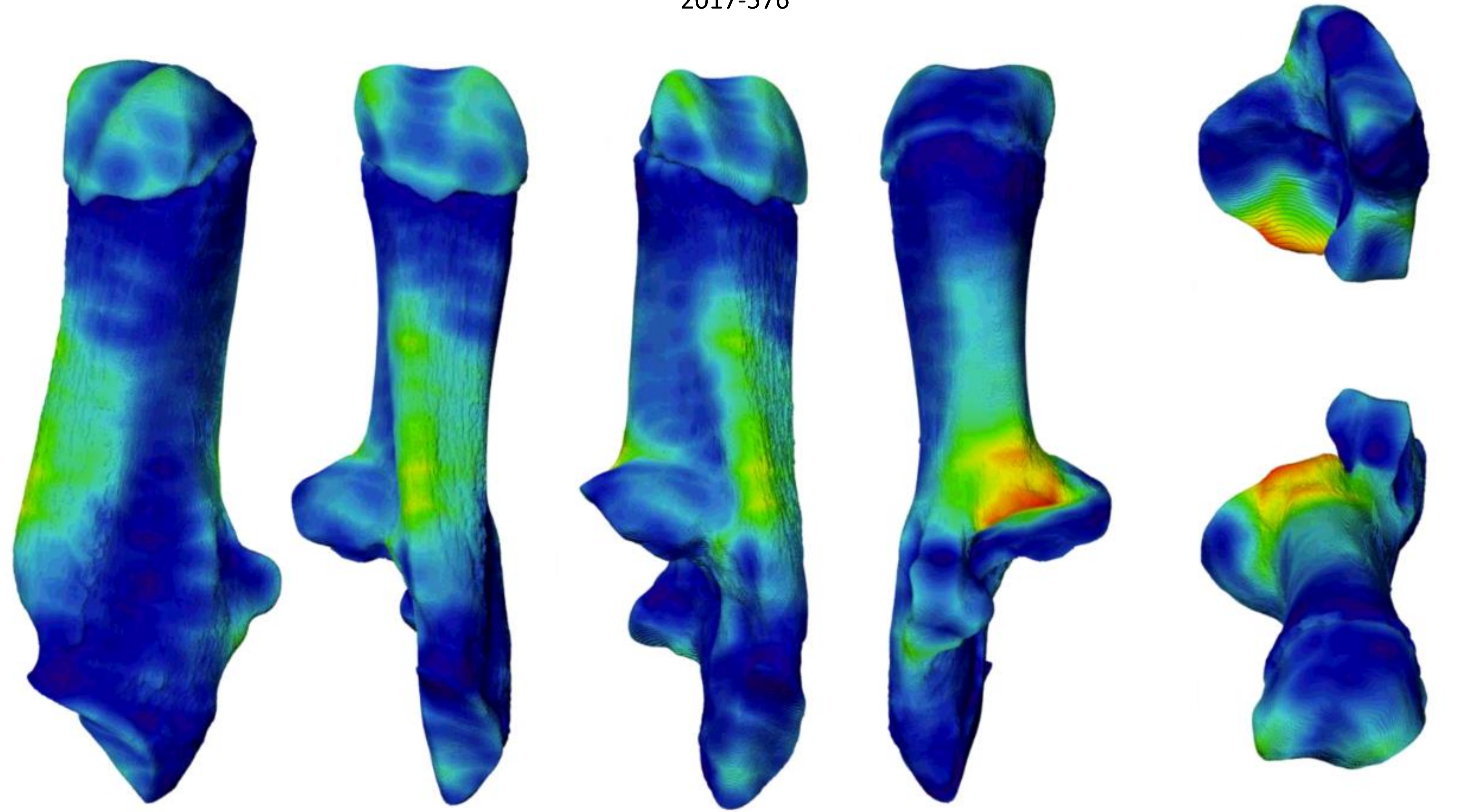

2017-581

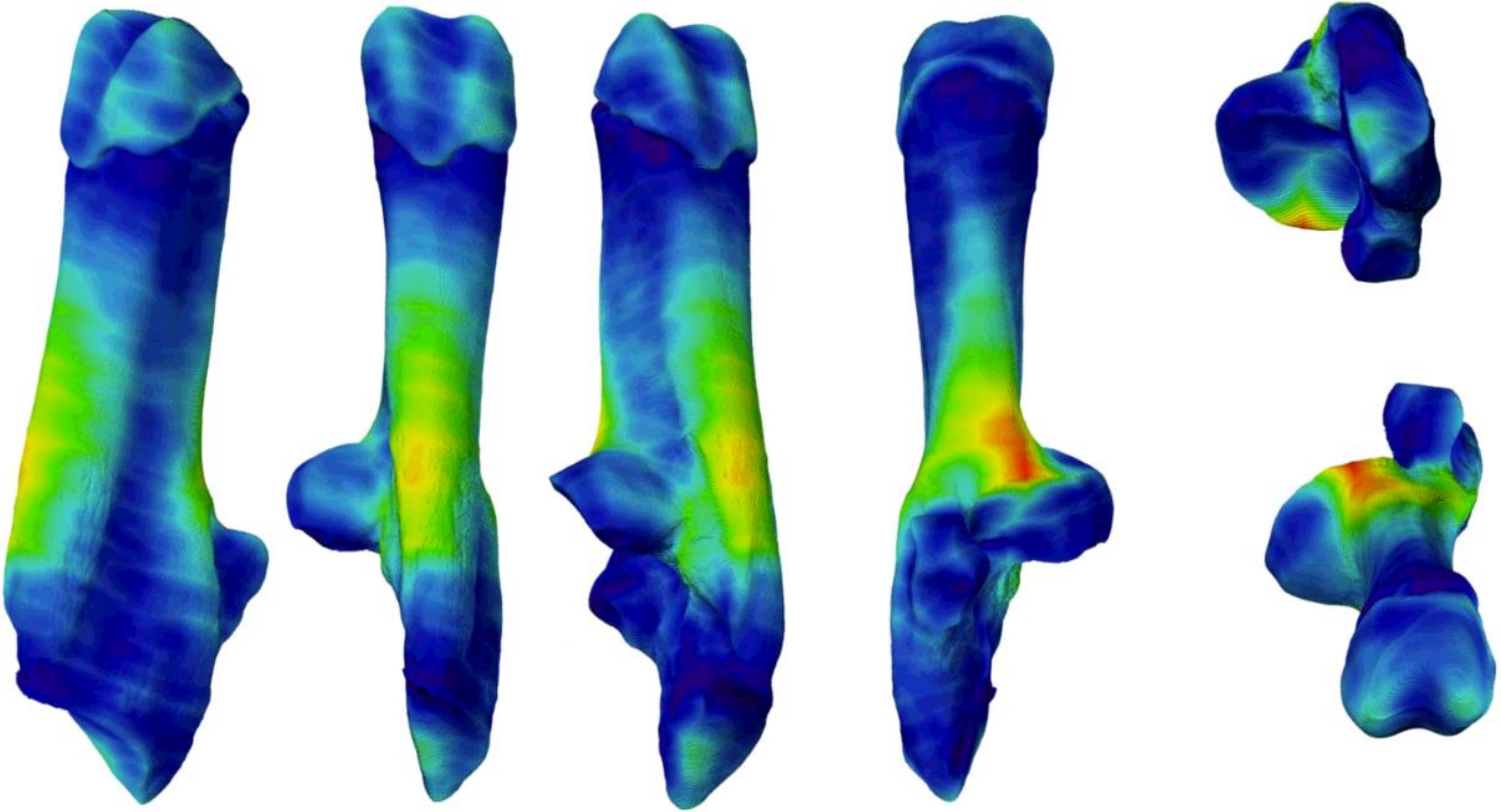

2017-569

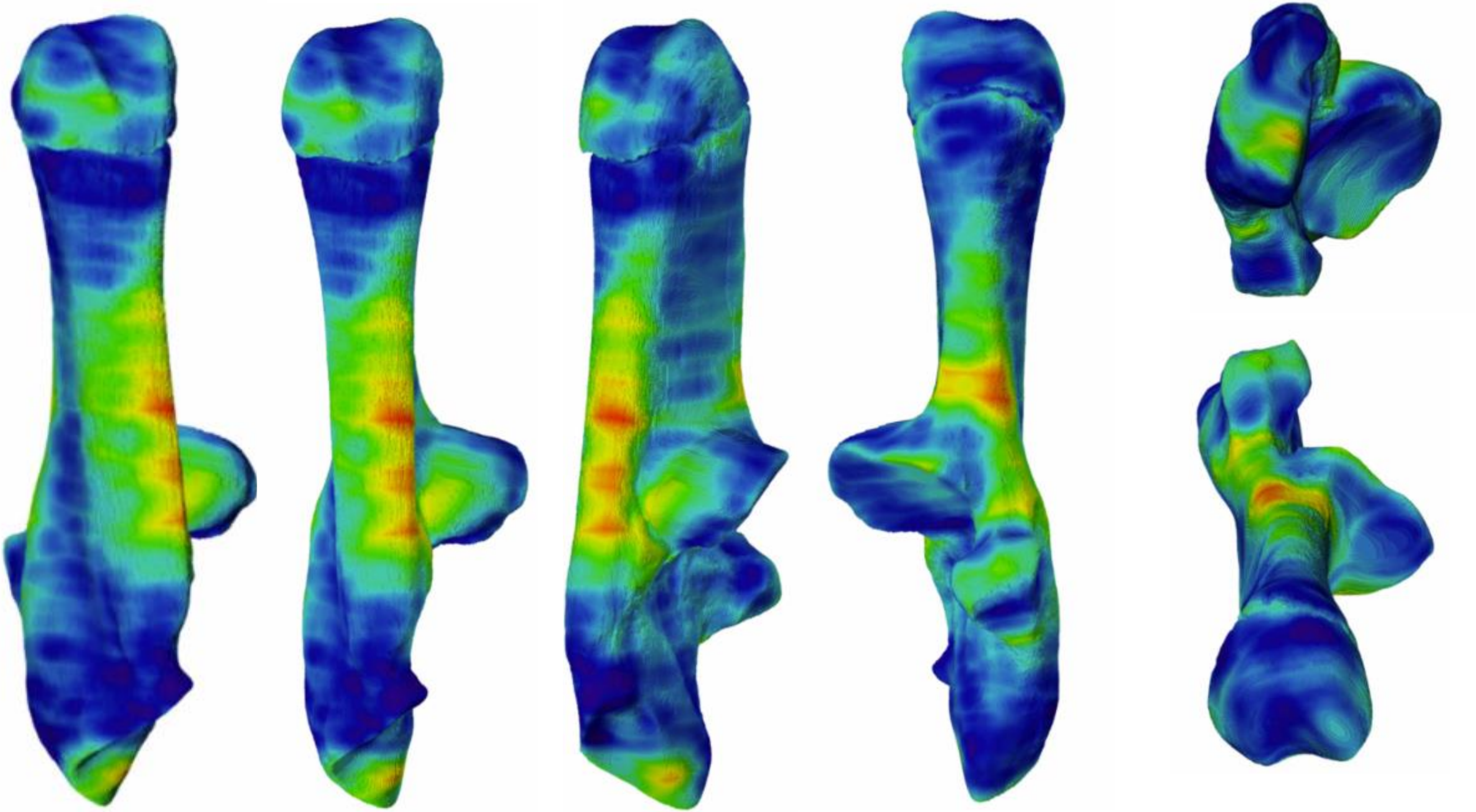

2017-579

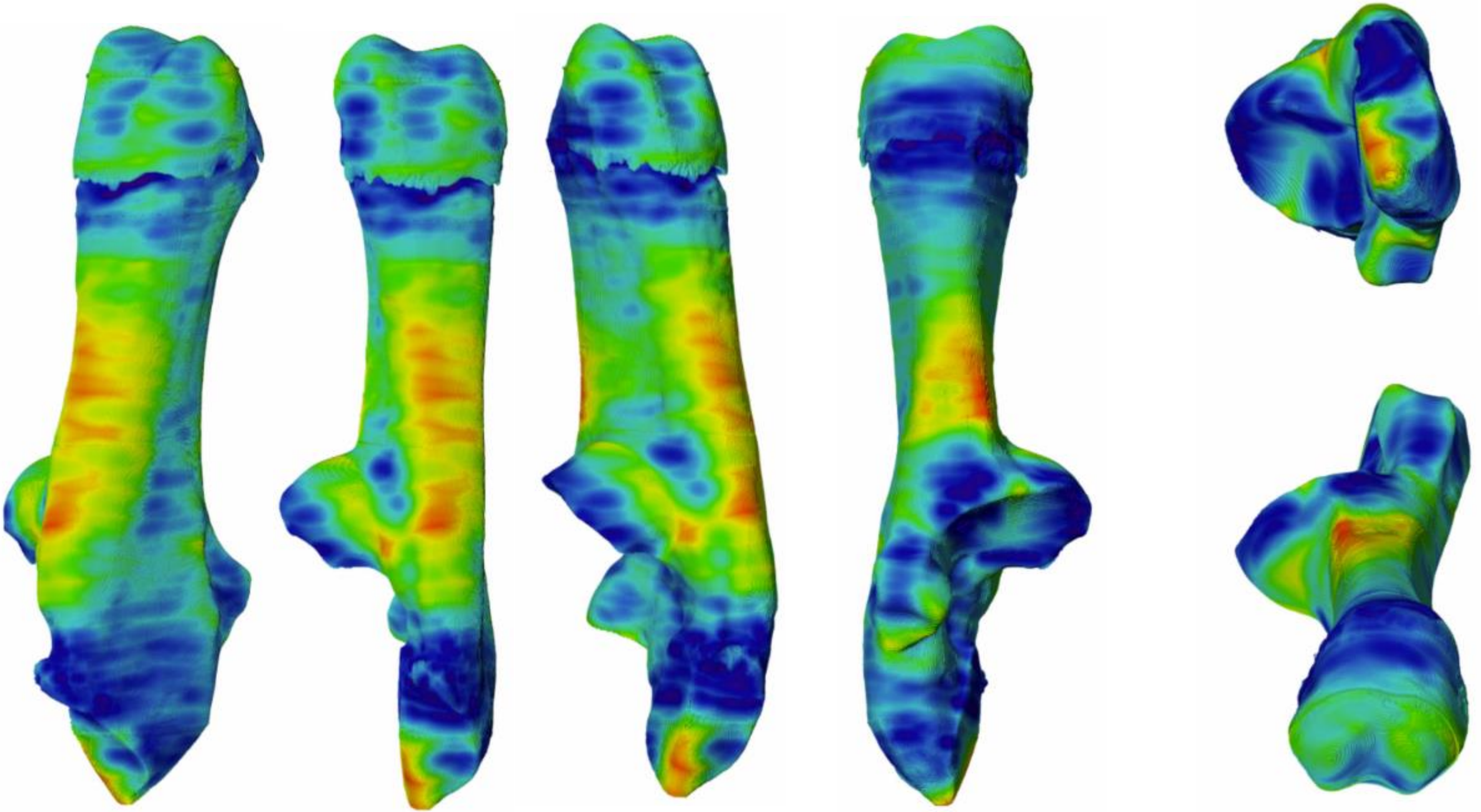

2017-8

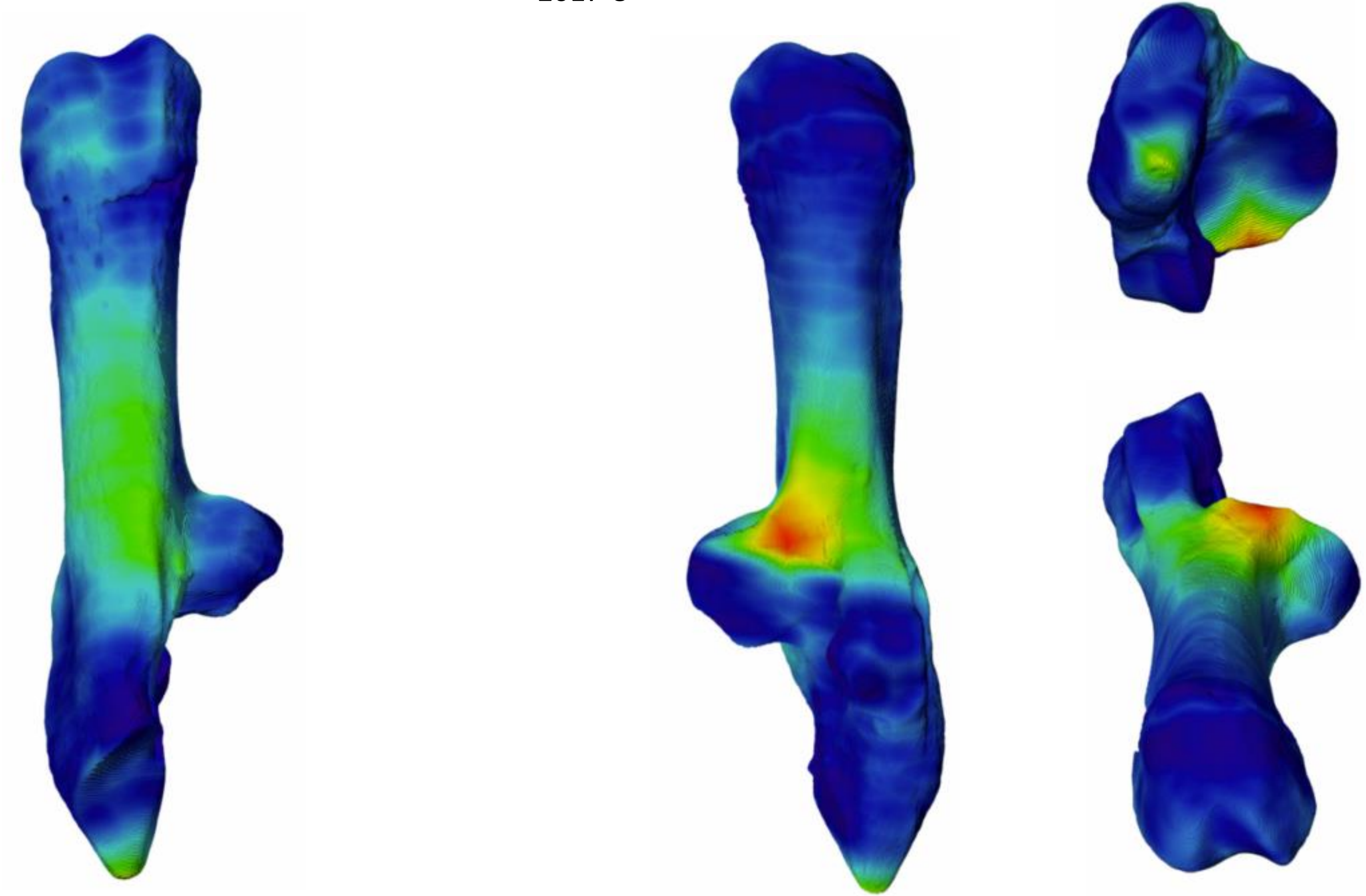

2017-559

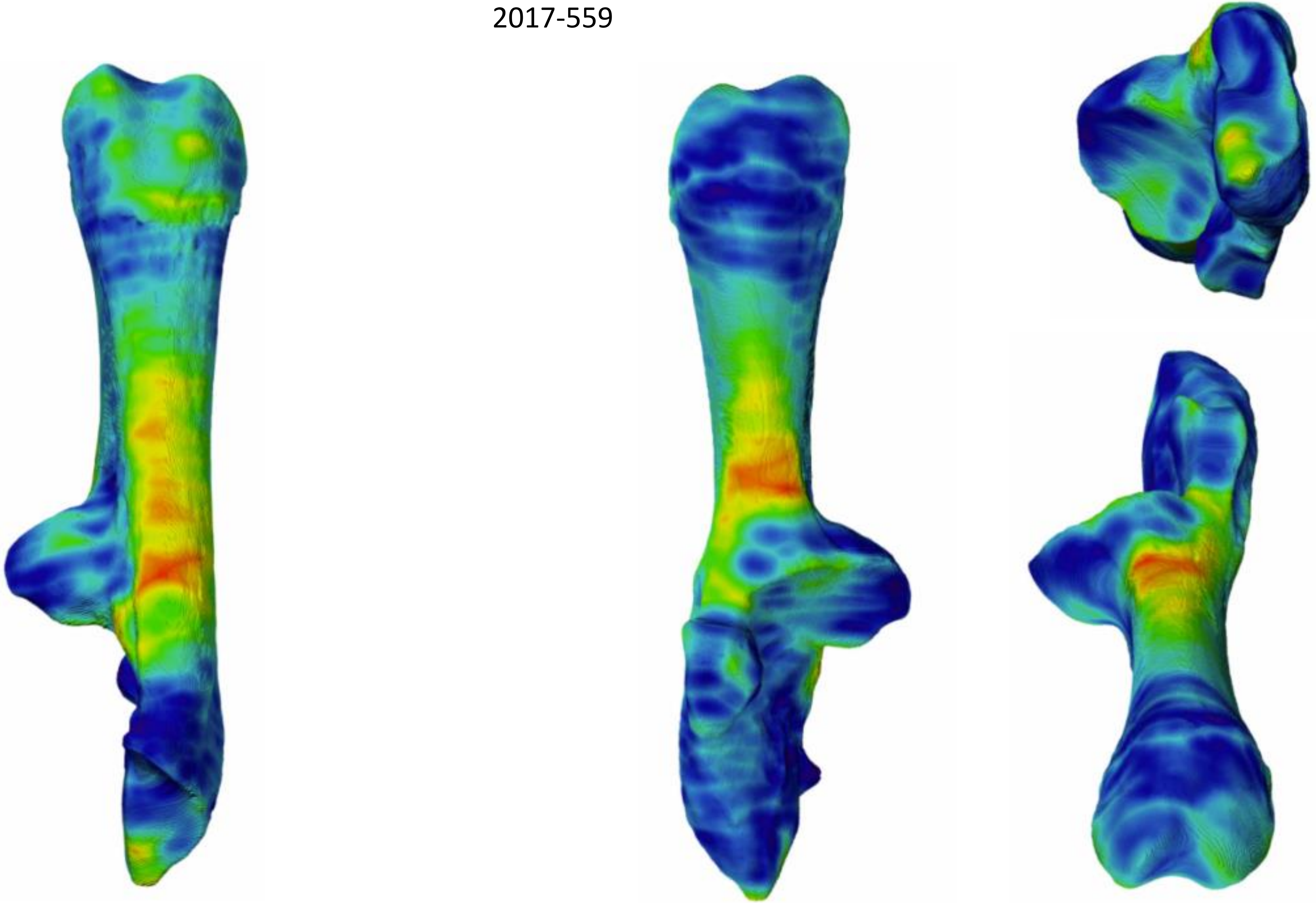

2017-570

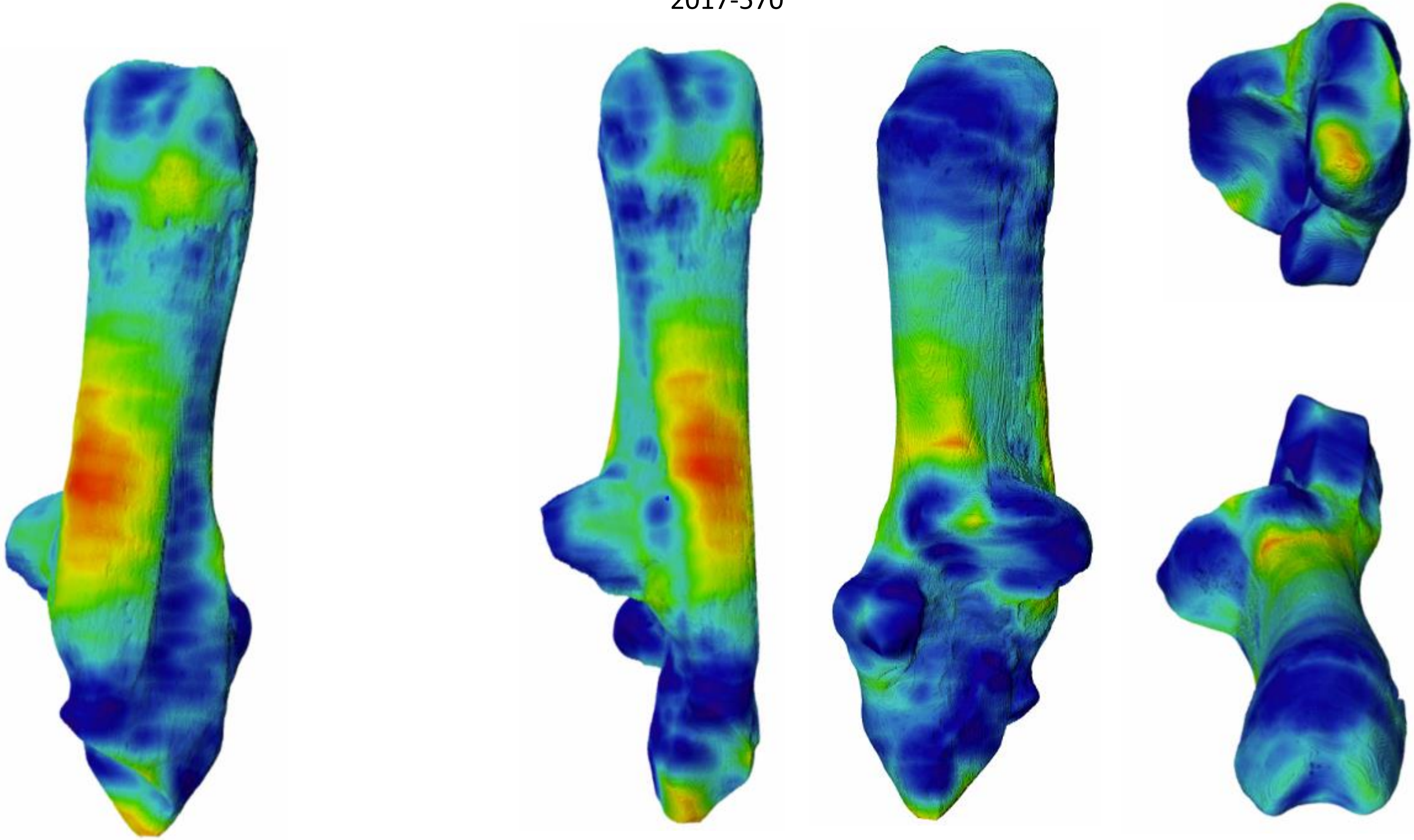

2017-573

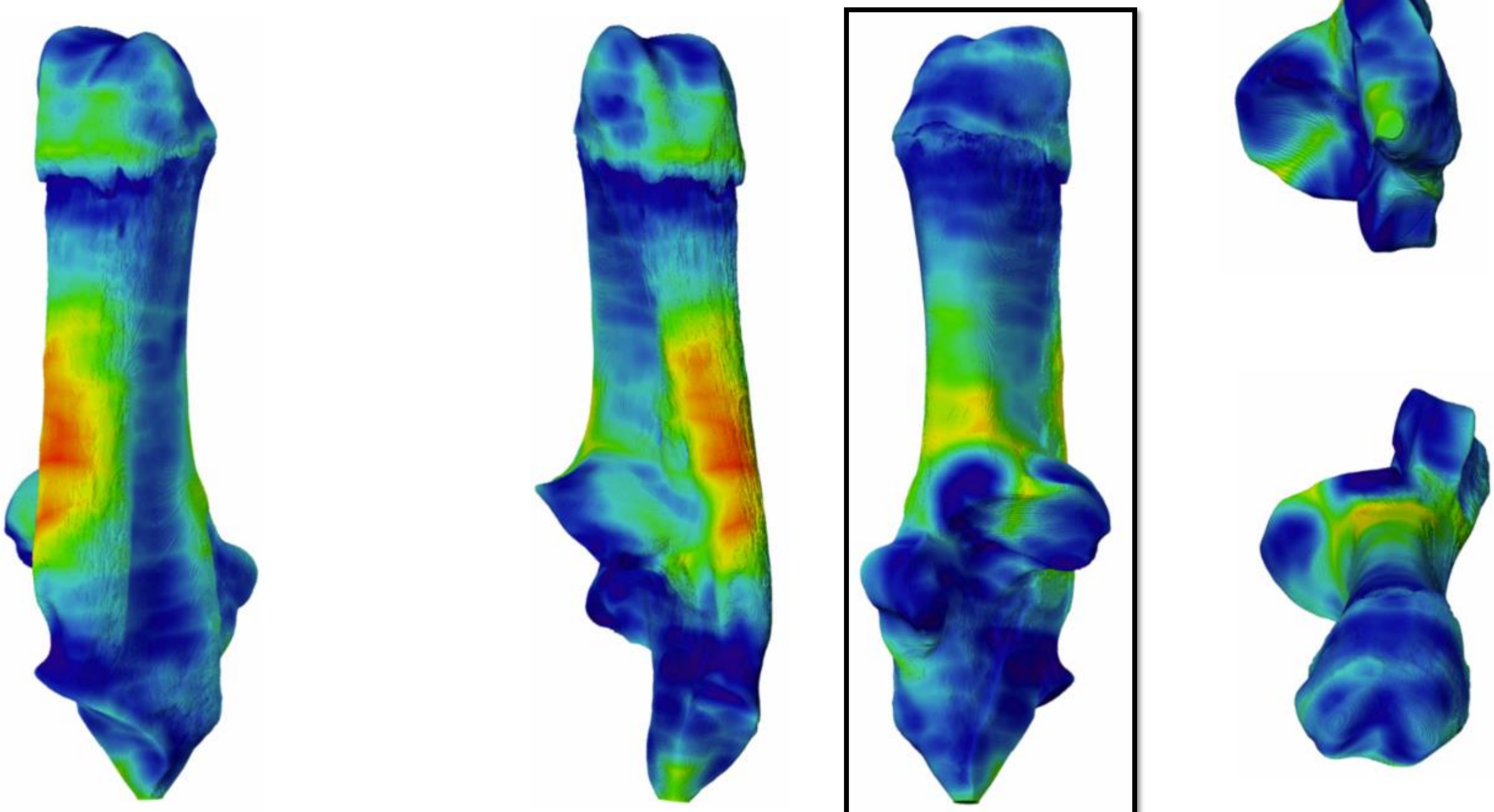

2017-554unnumb

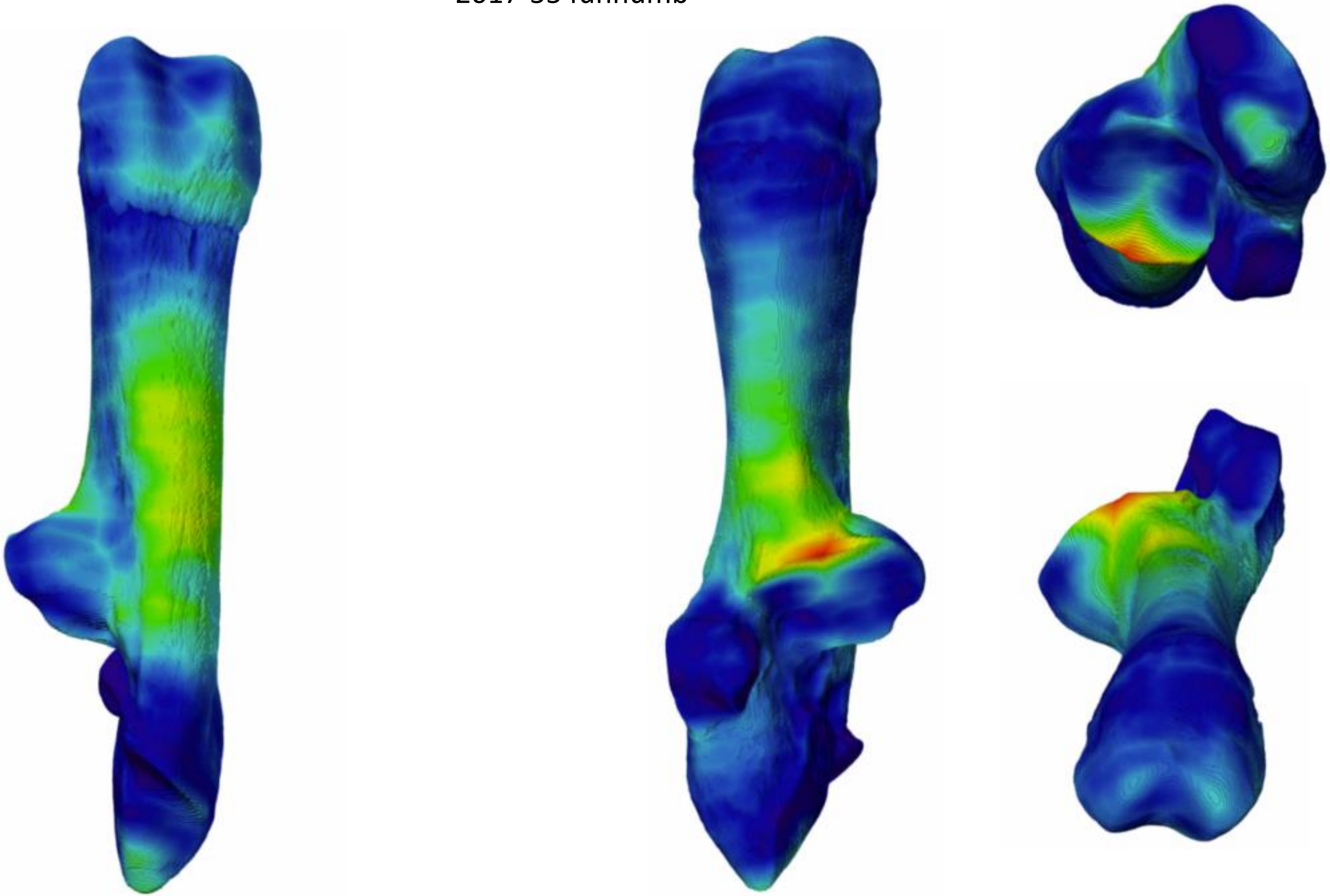

2017-558

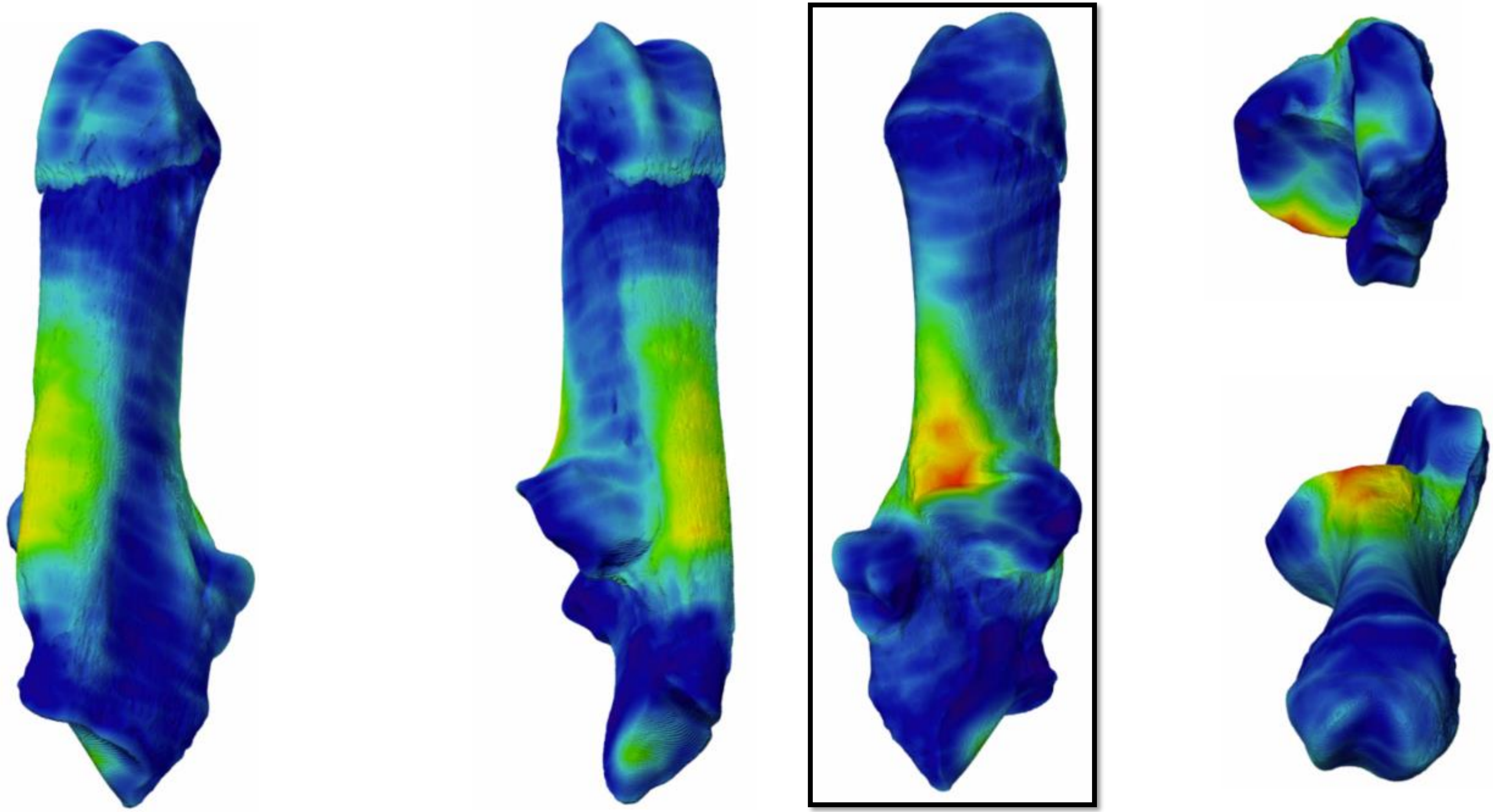

2017-562

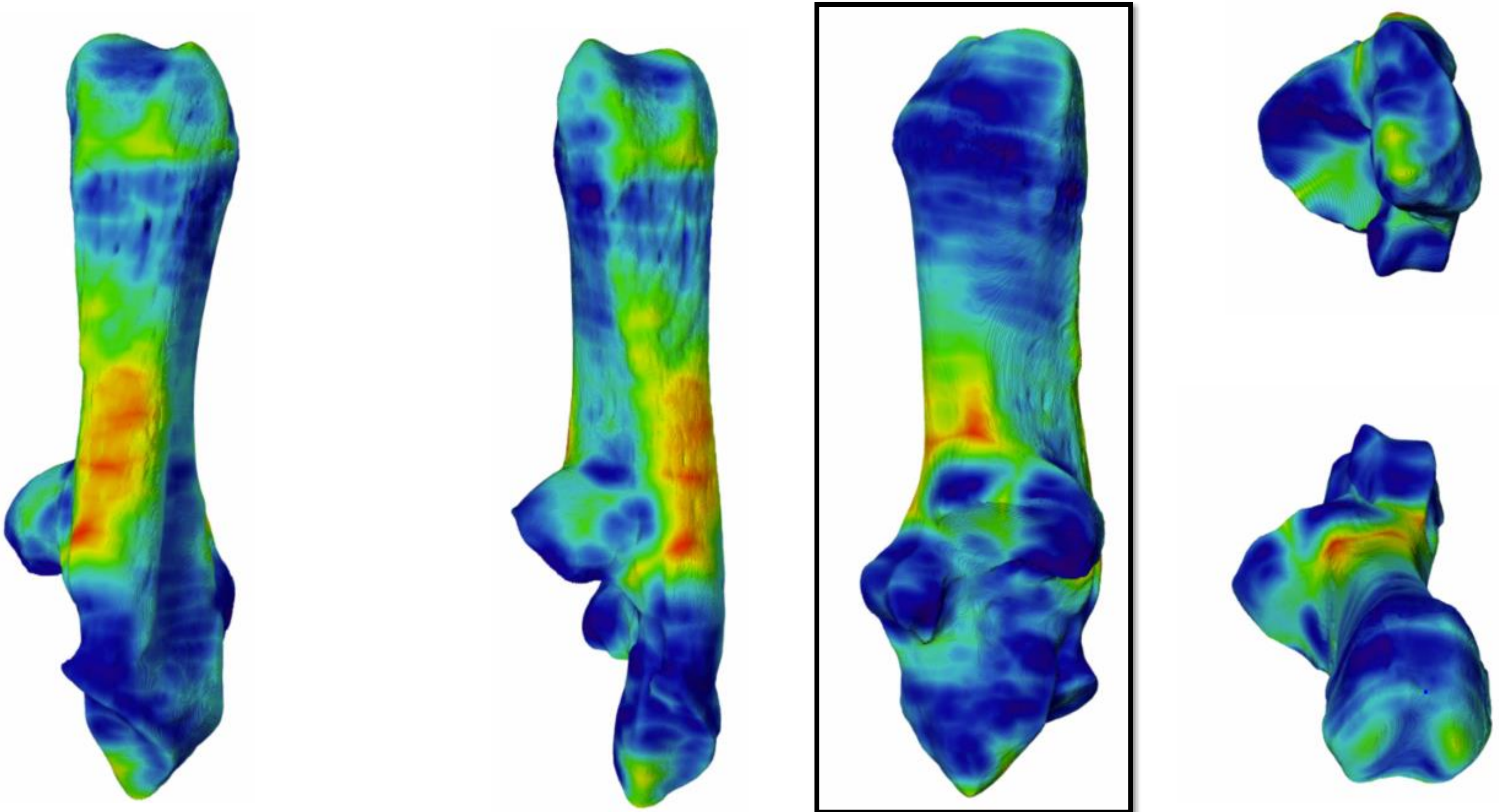

2017-574

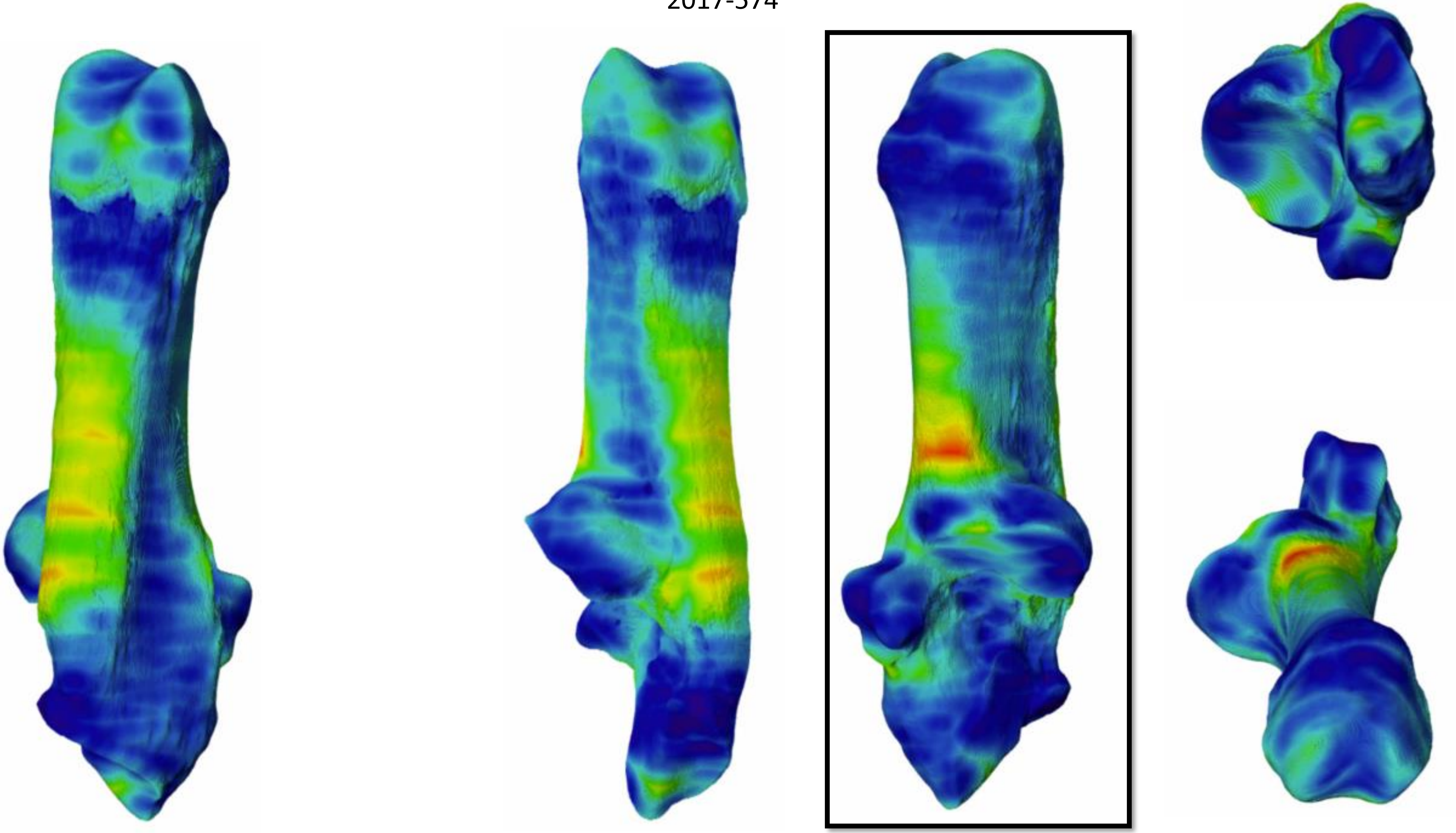

2017-554

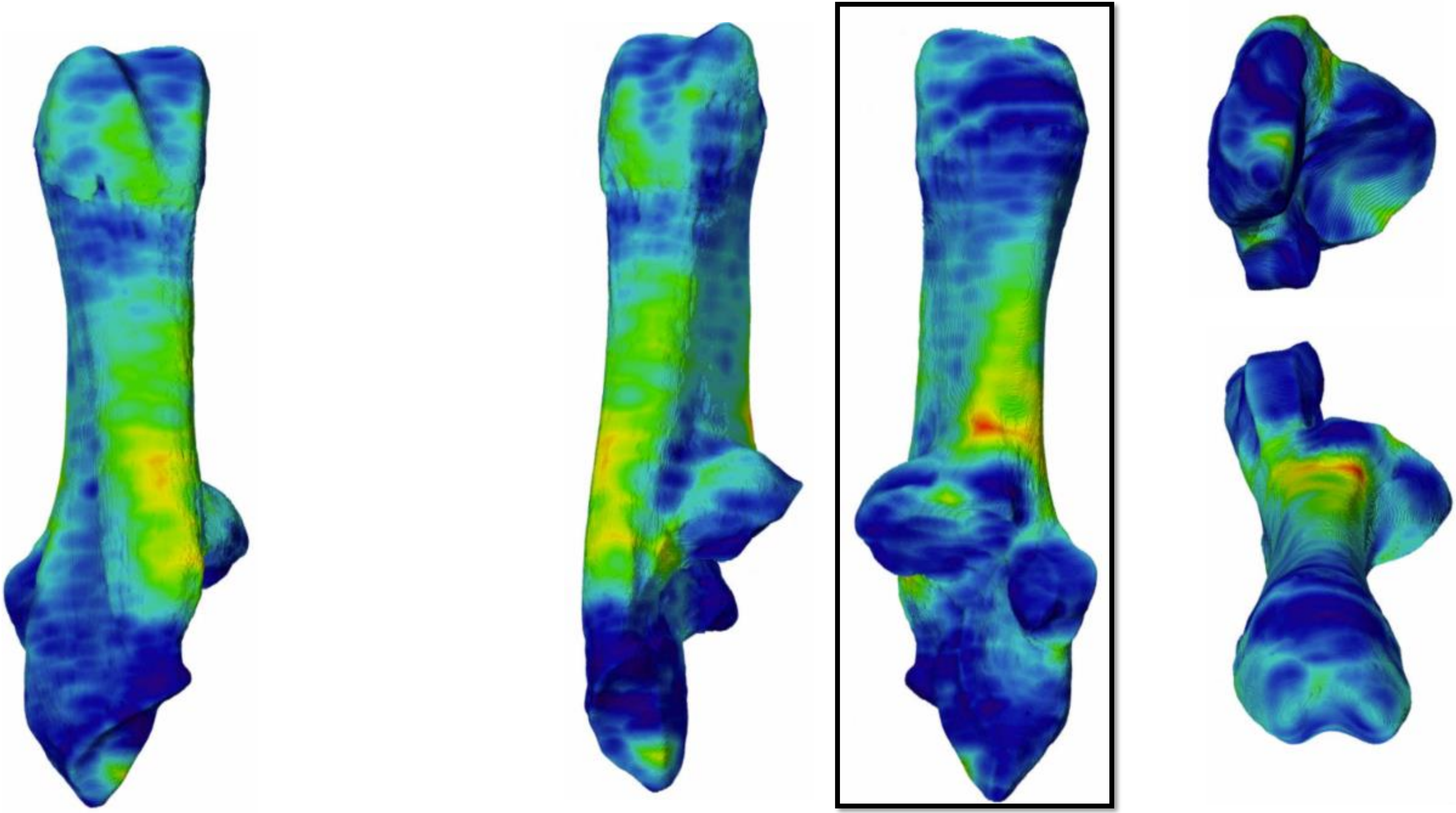

2017-556

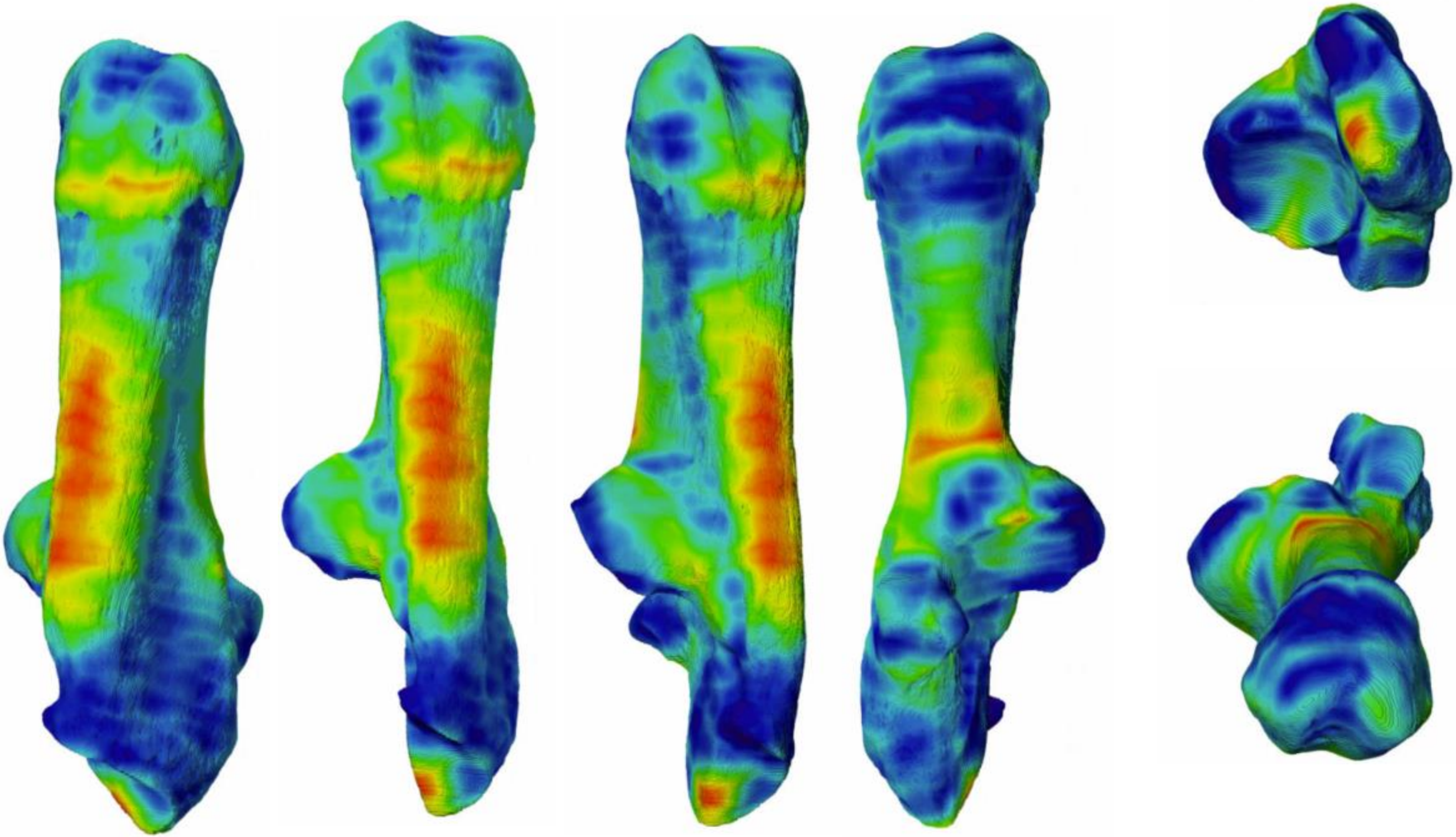

2017-561

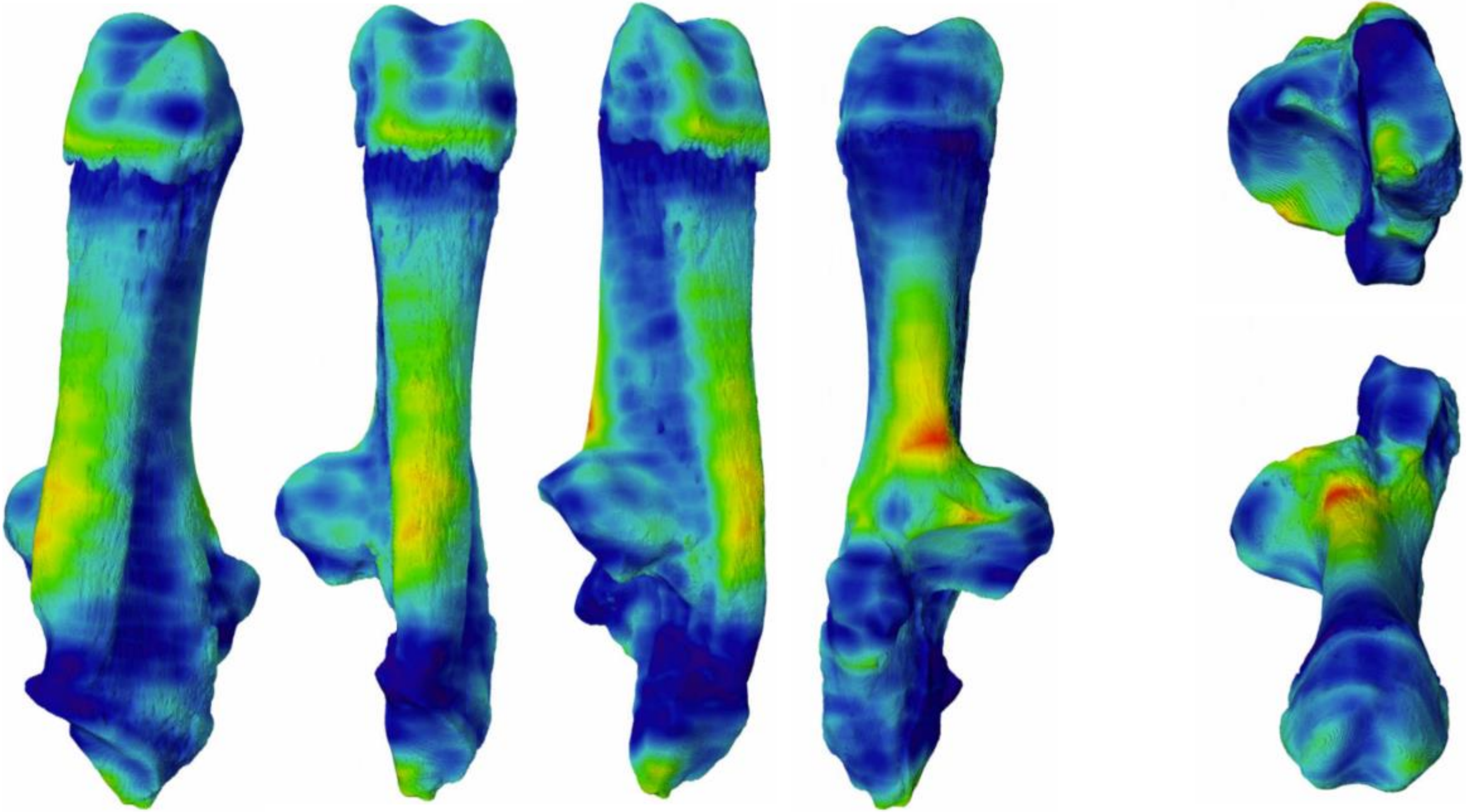

2017-571

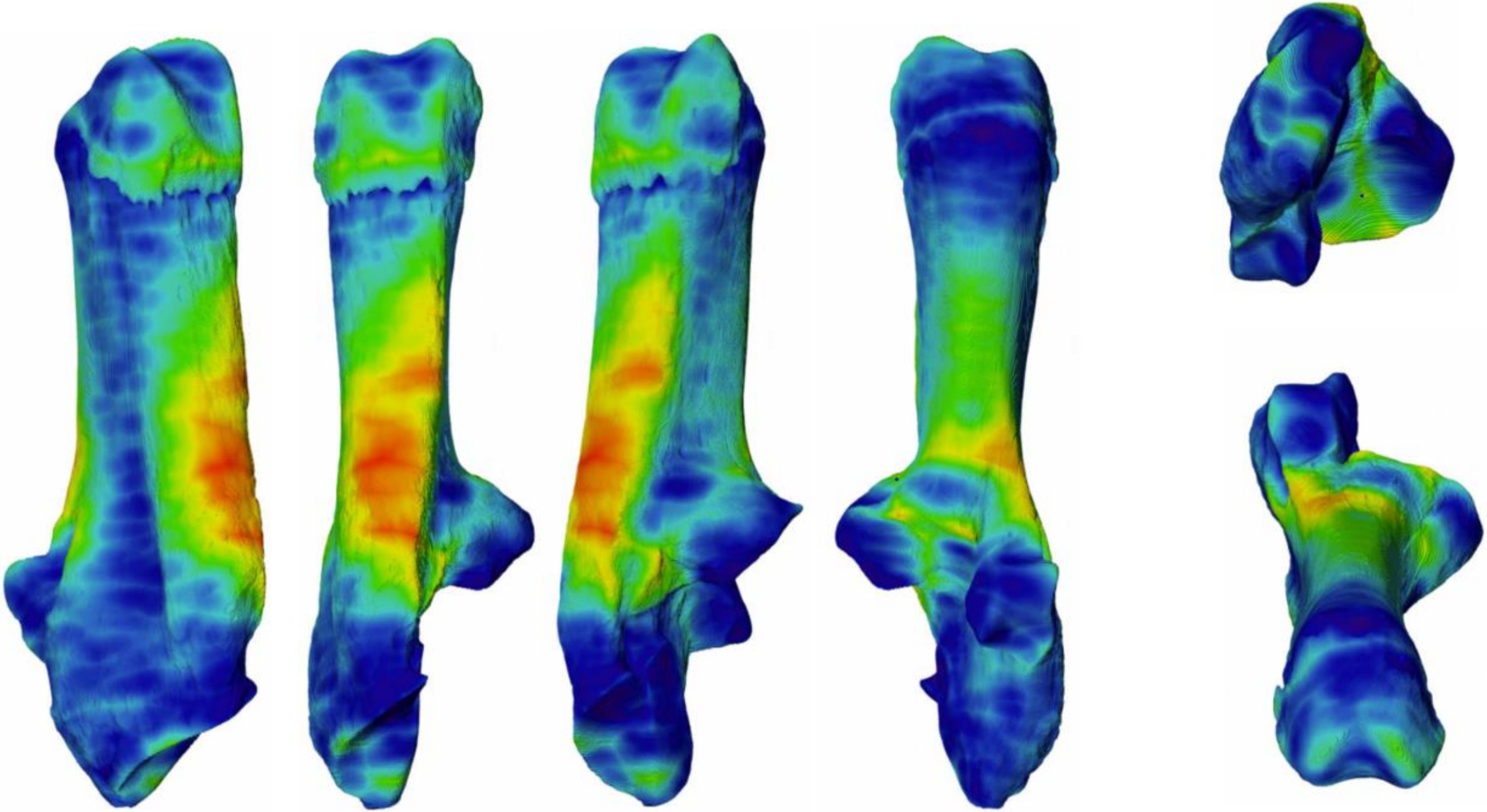

2017-555

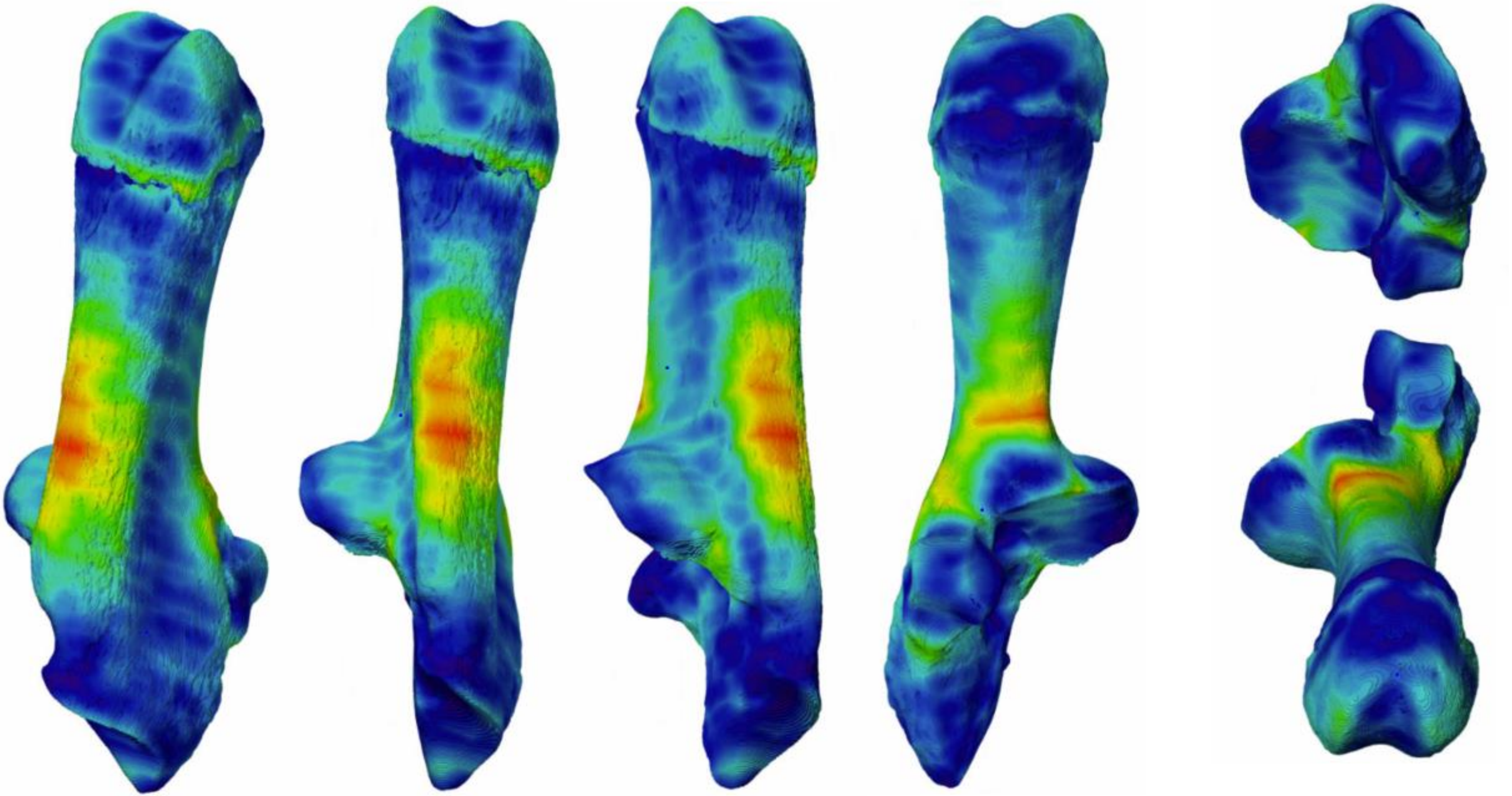

2017-557

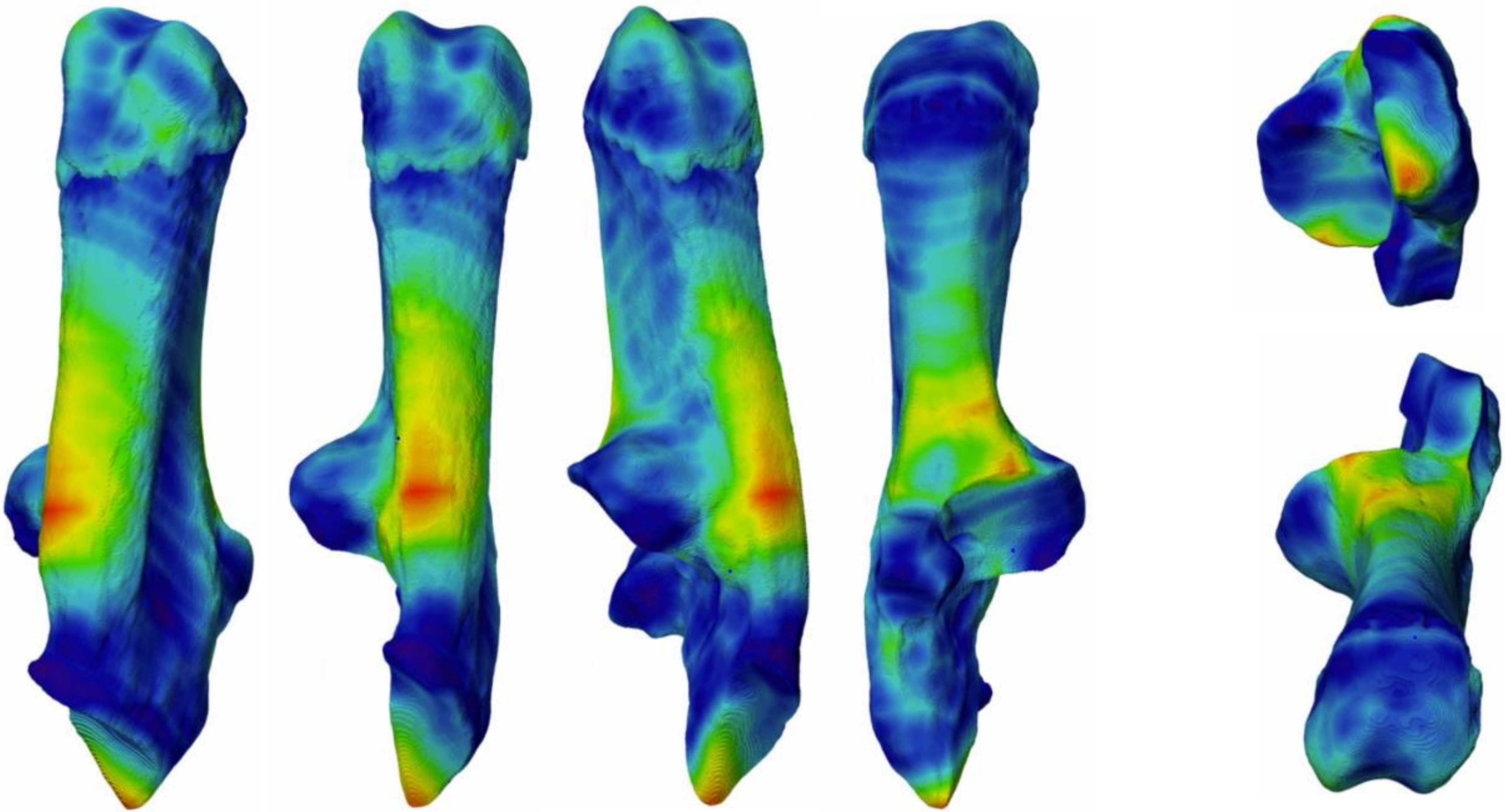

2017-572

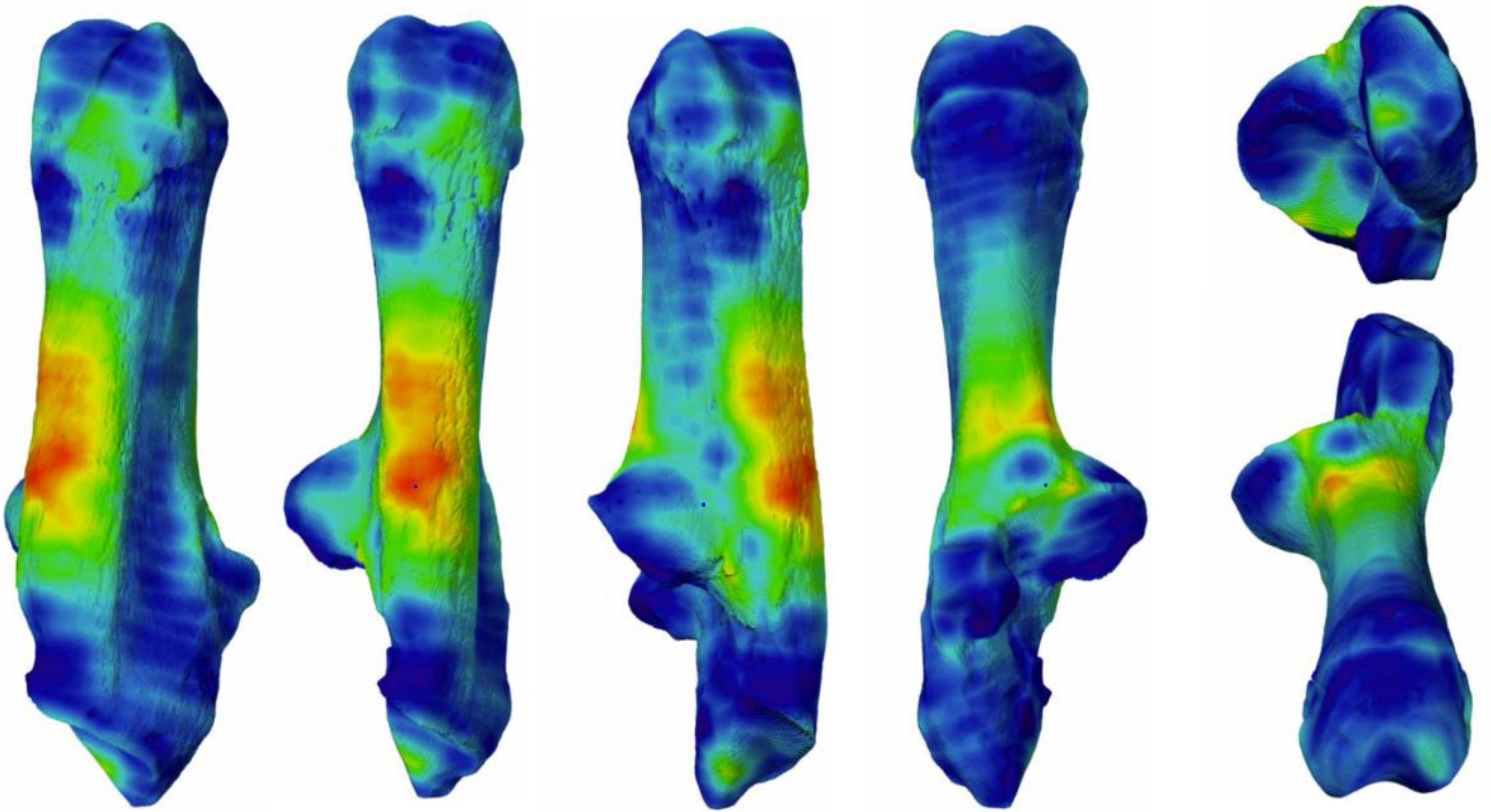

2017-575

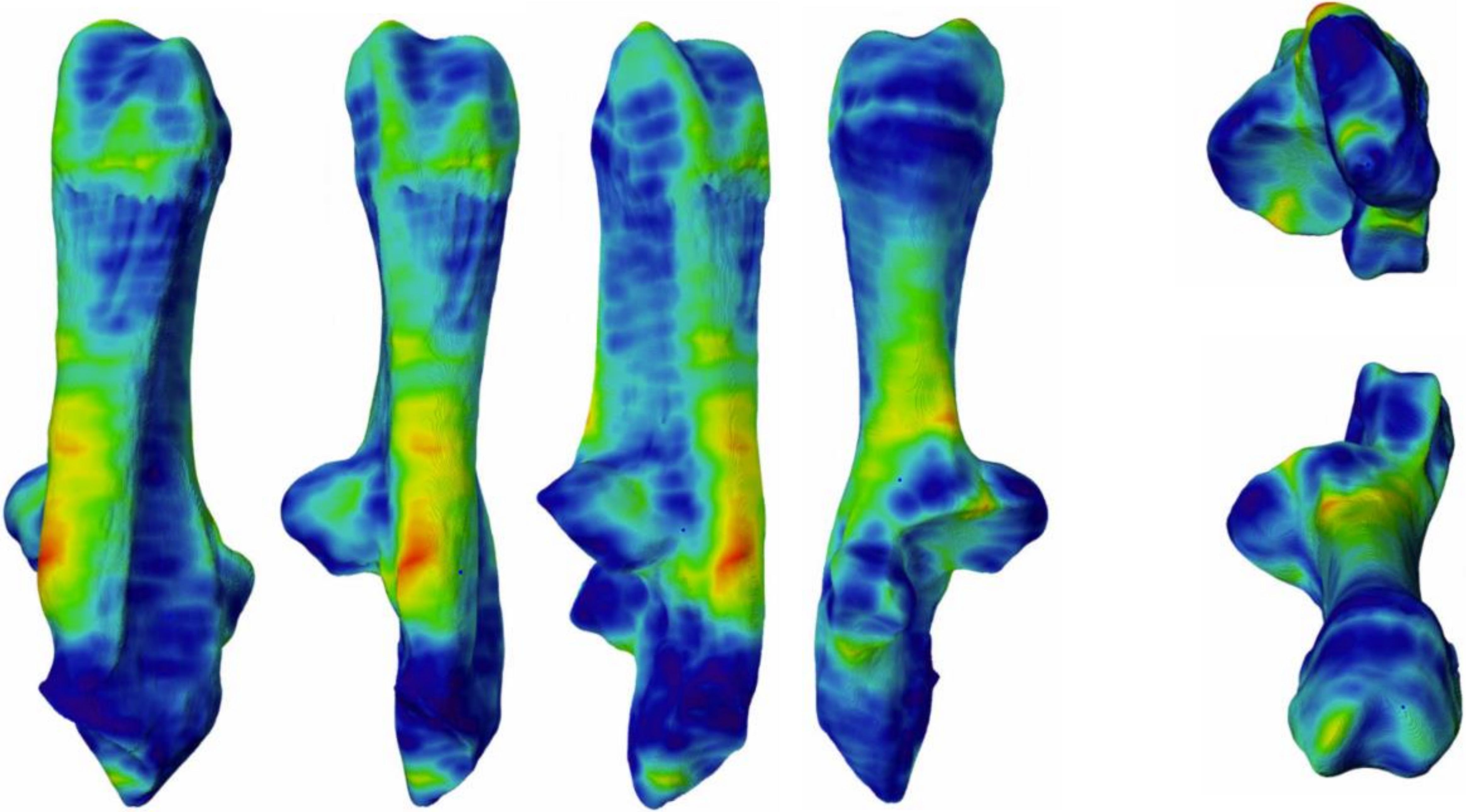

2017-c235

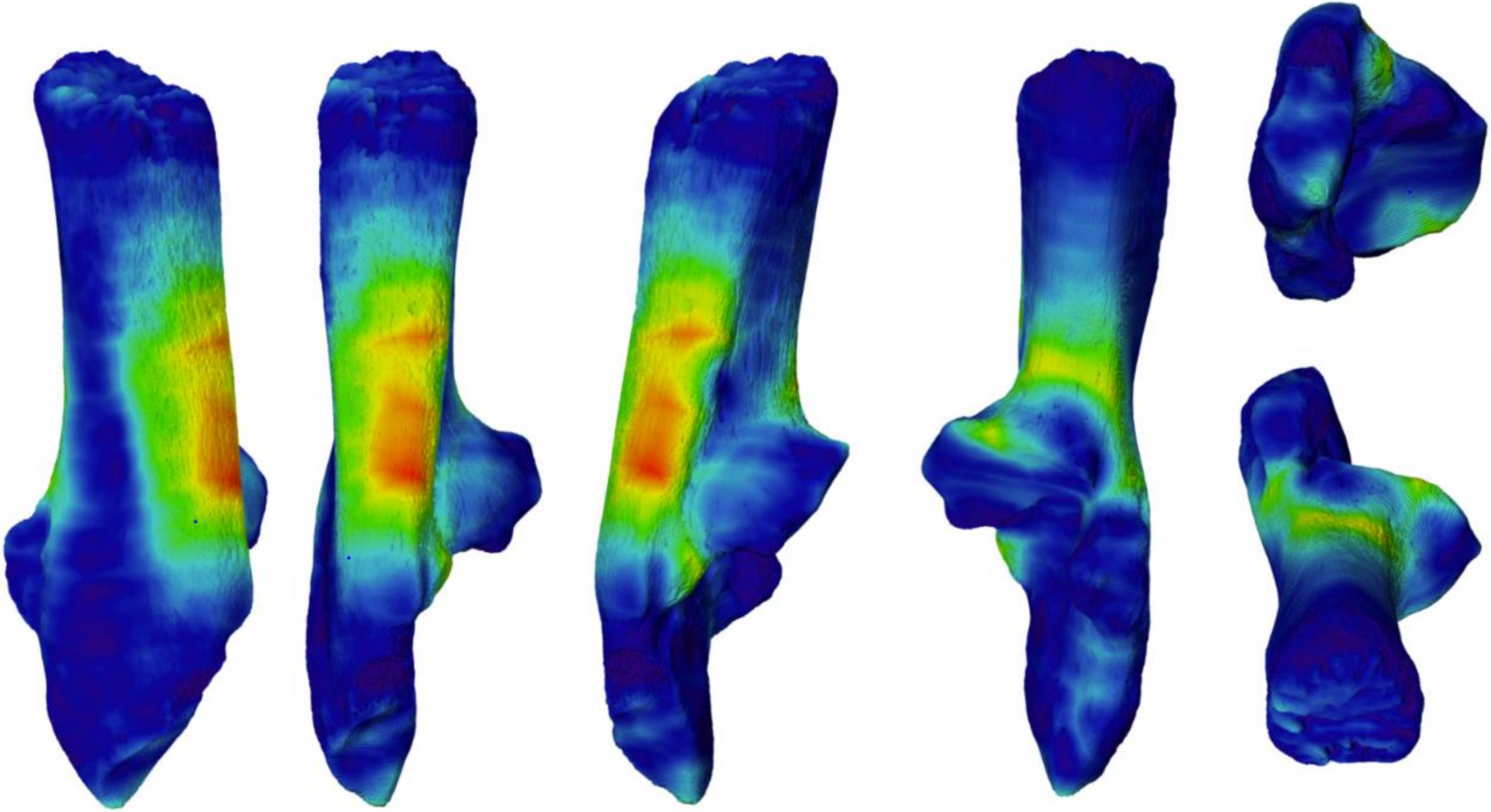

2017-2136

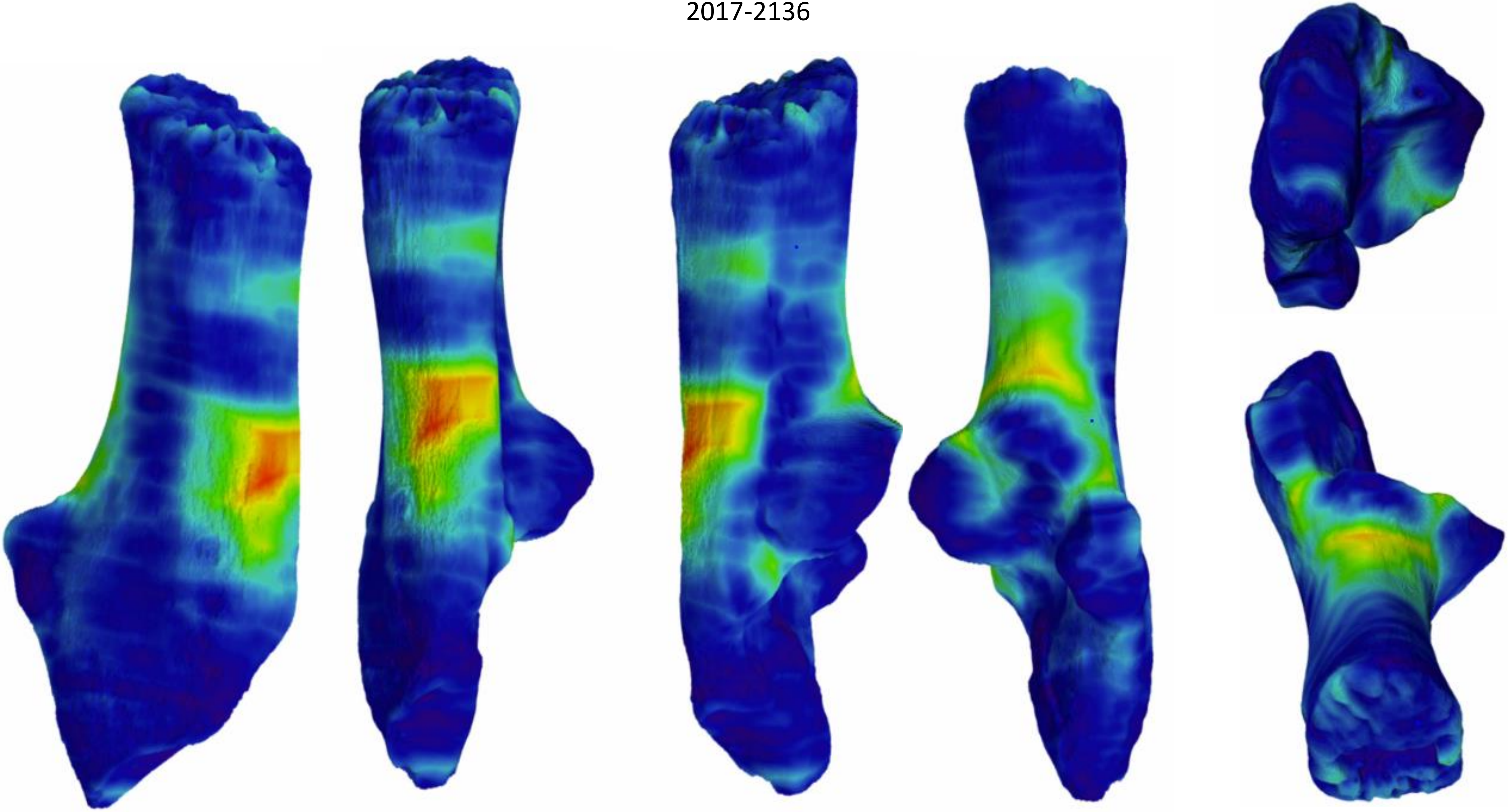

2017-c2139

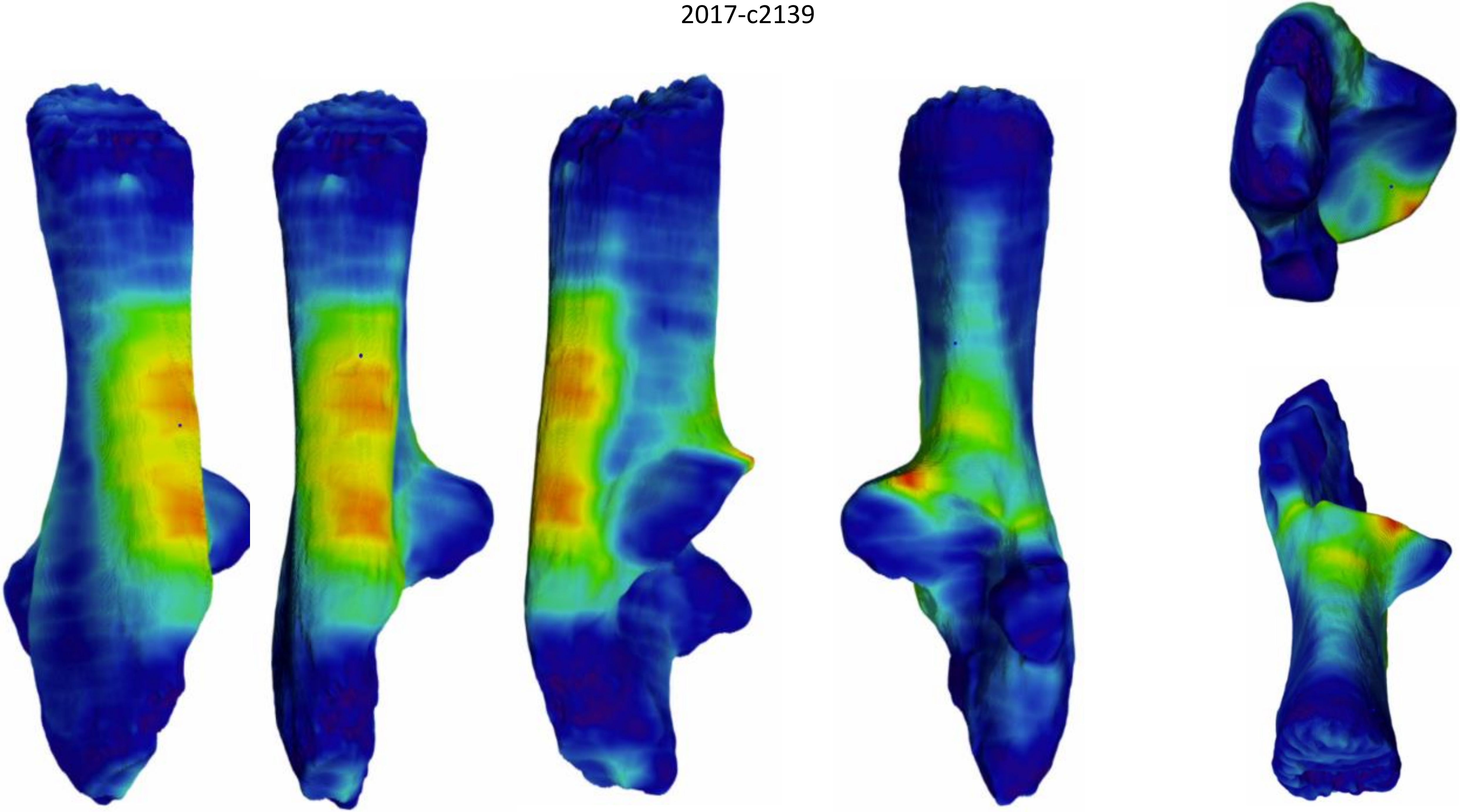

2017-c214

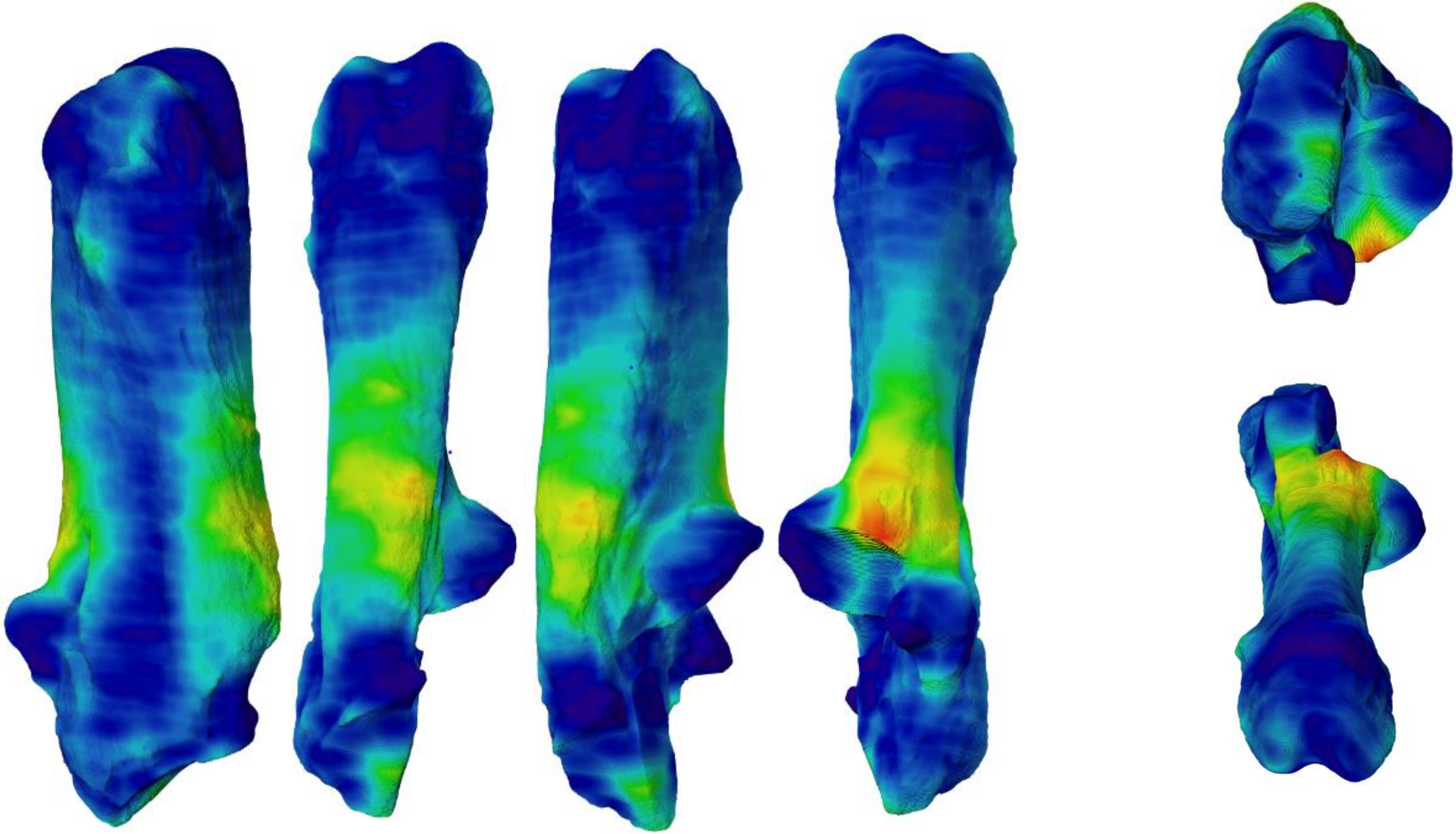

2017-Pradat184

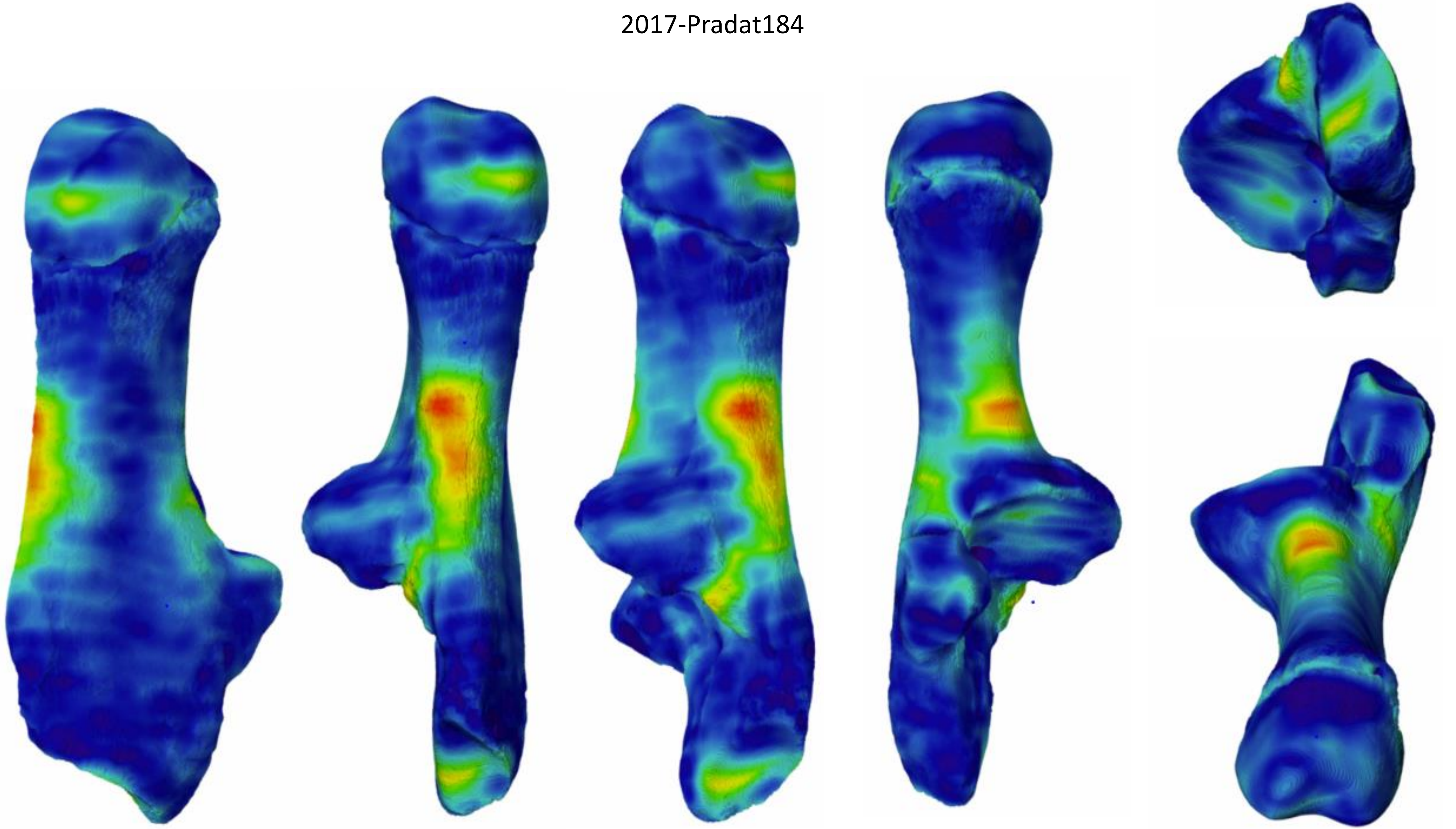

2017-Pradat187

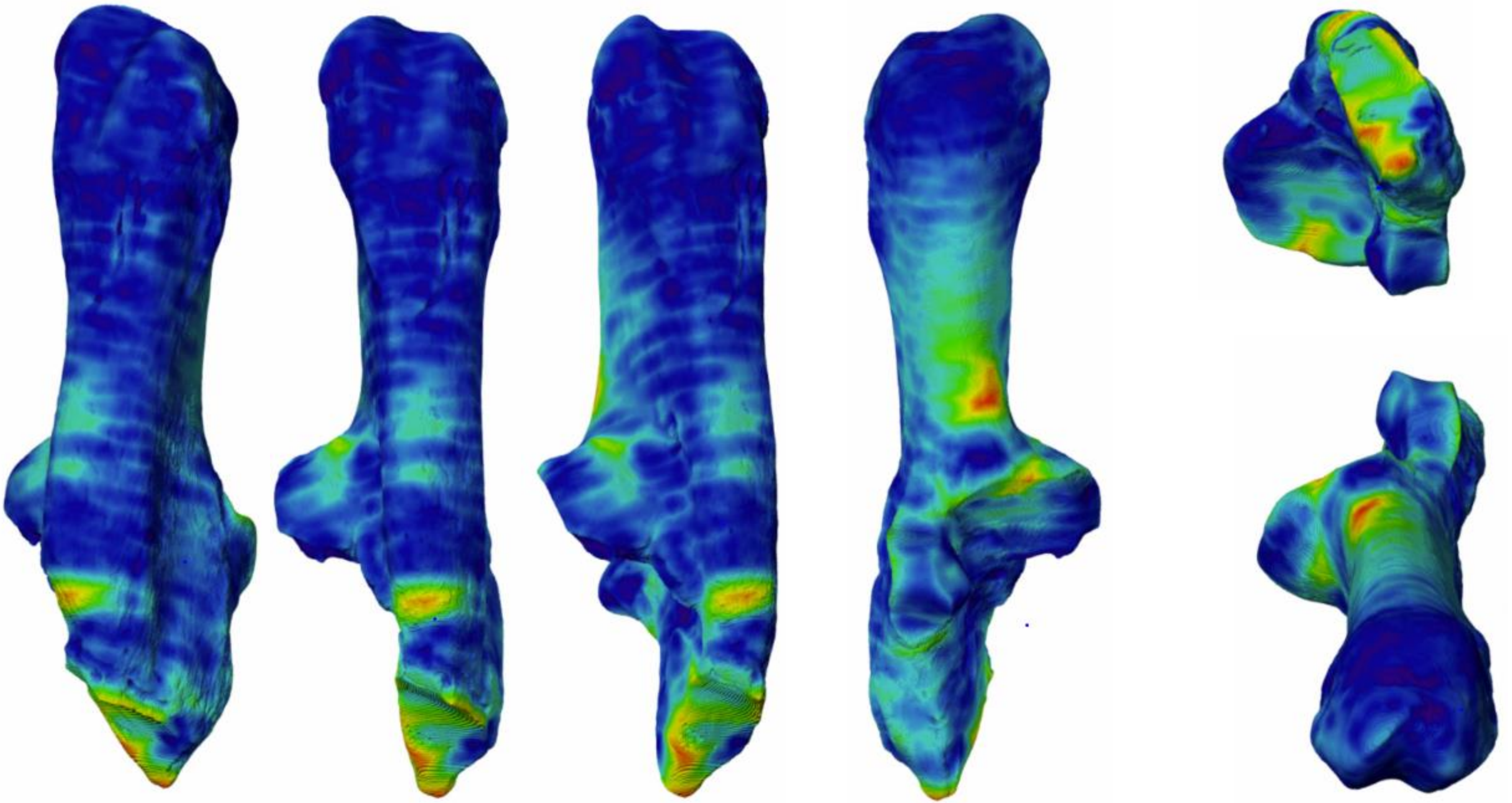

2017-560

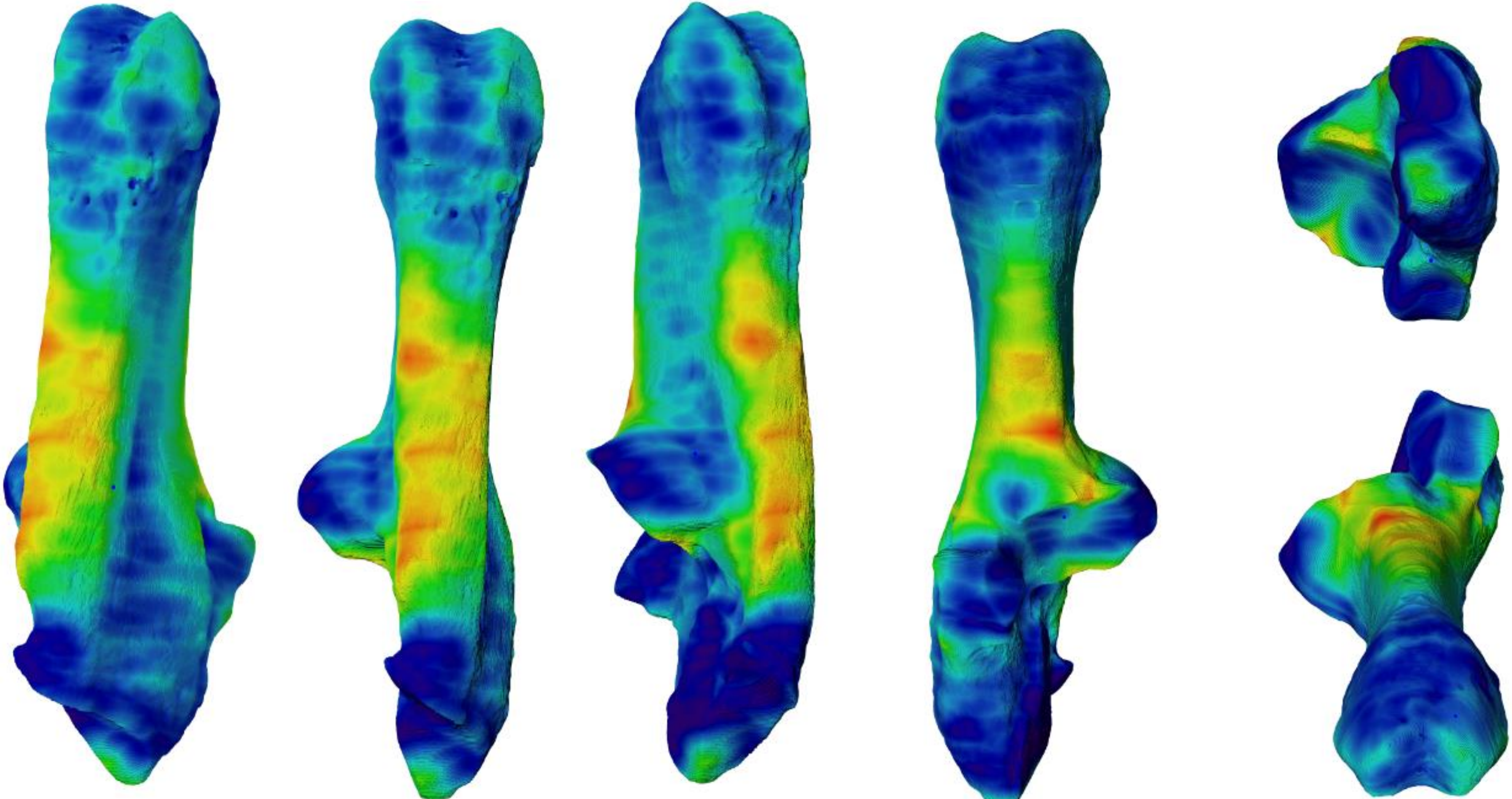

2017-563

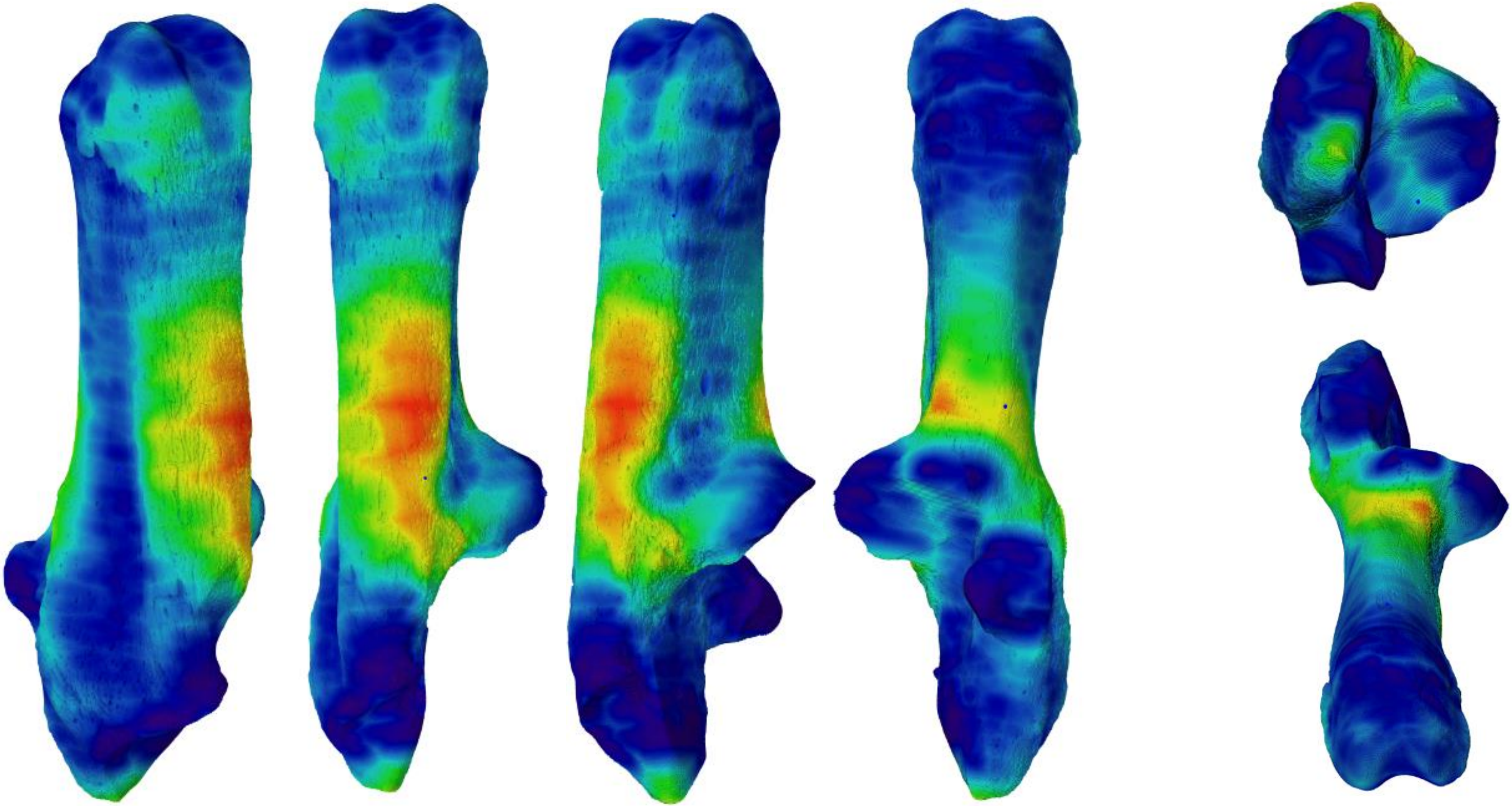

2017-564

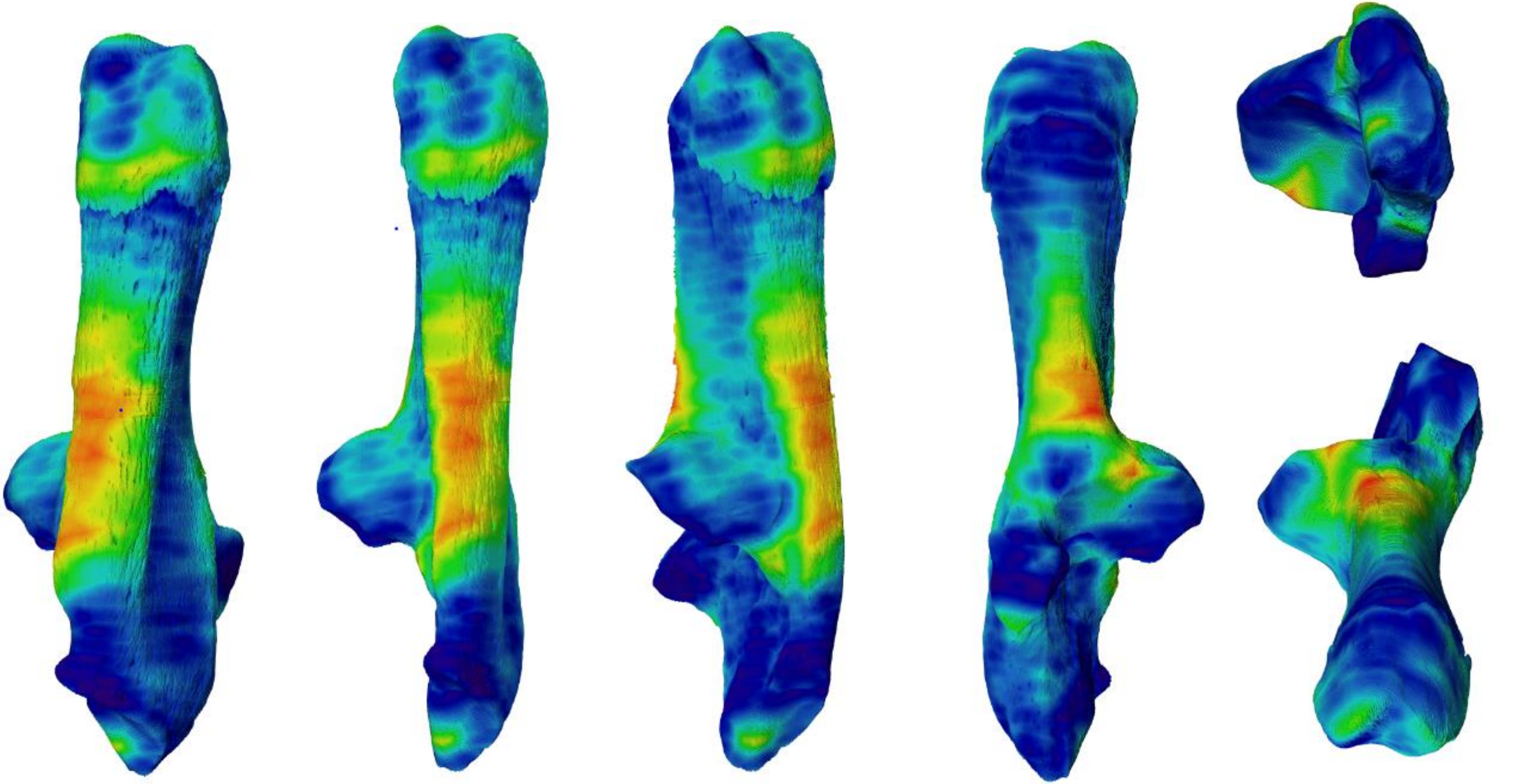

2017-568

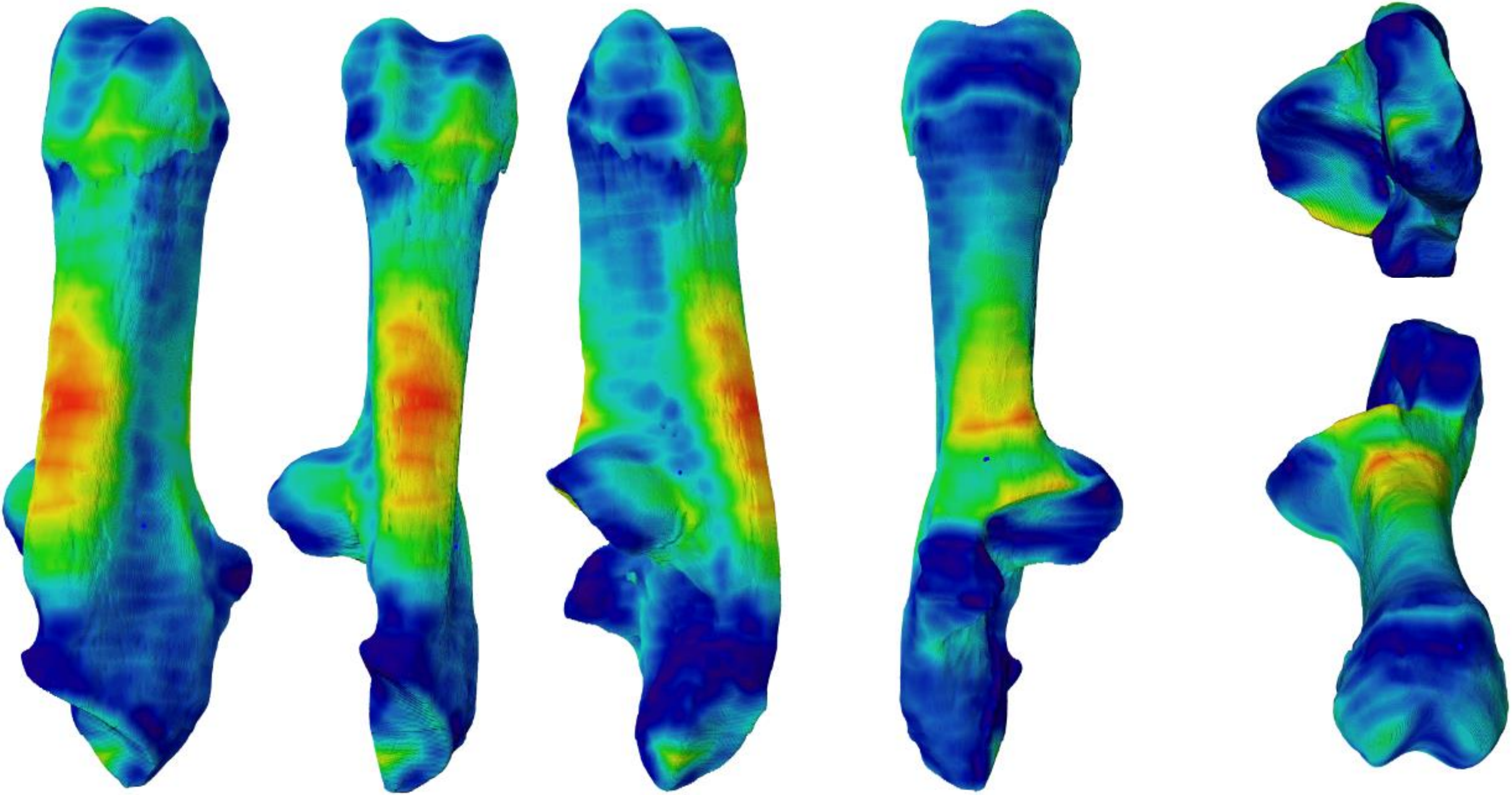

COMP1OS1

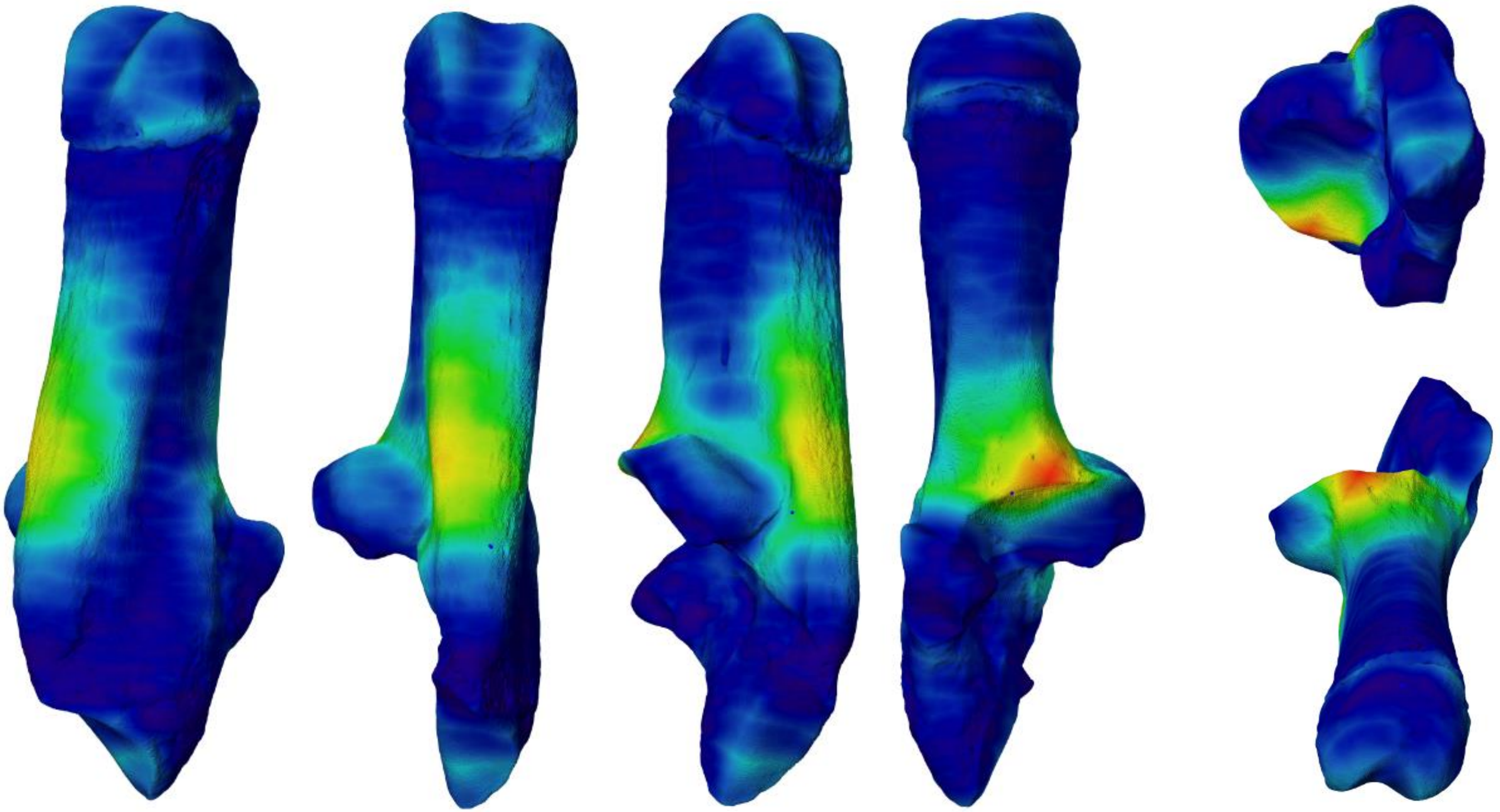

COMP1OS2

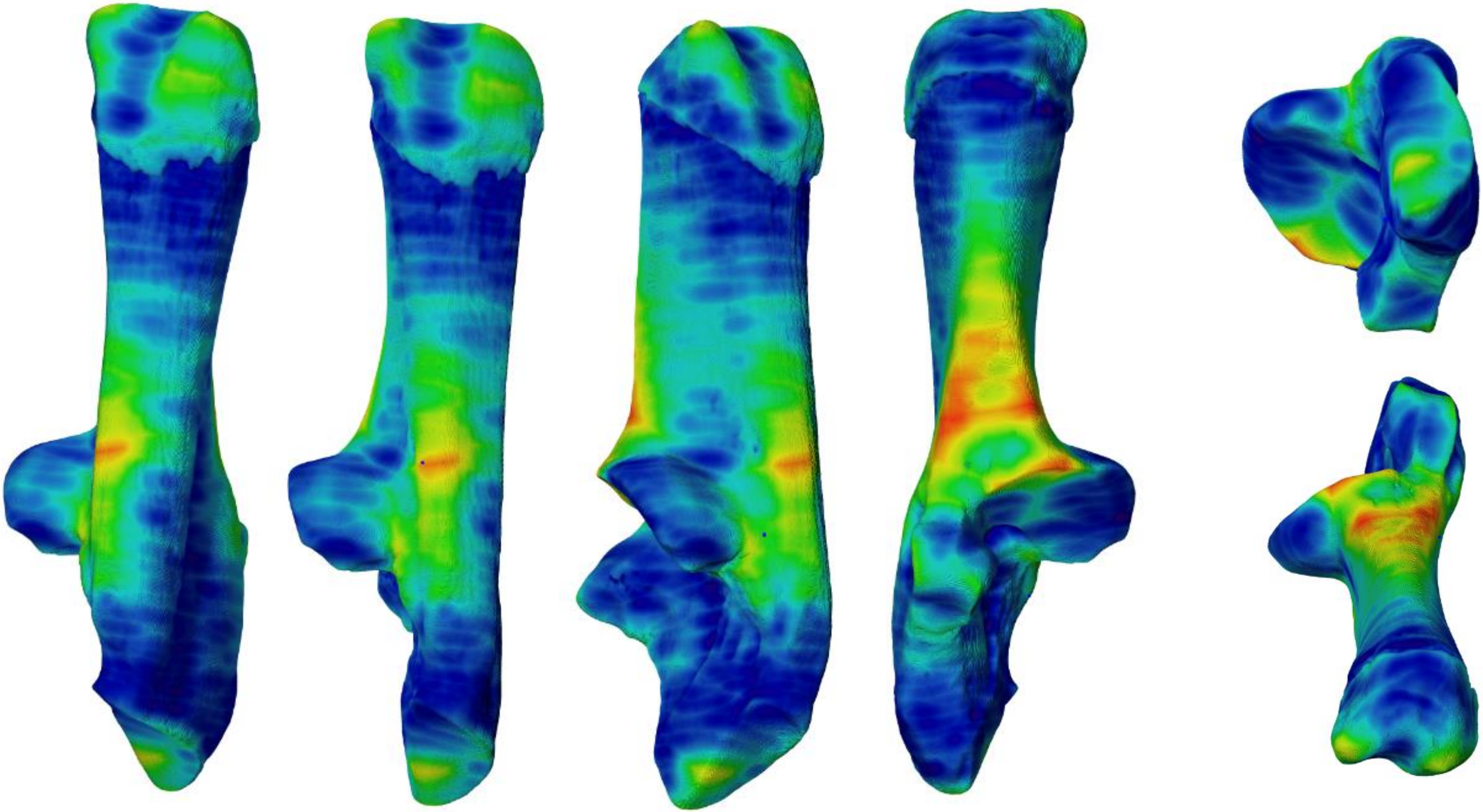

COMP1OS3

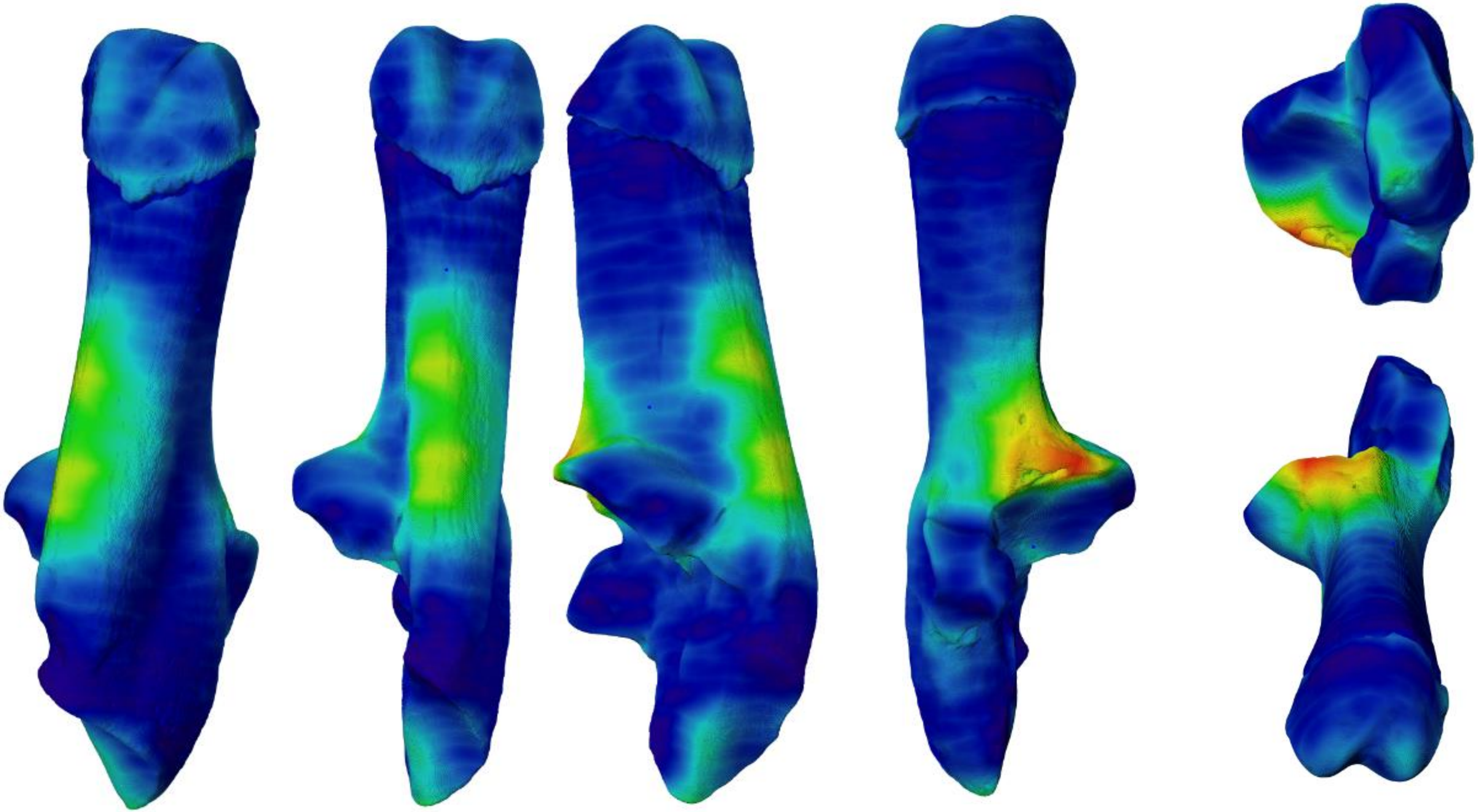

COMP1OS4

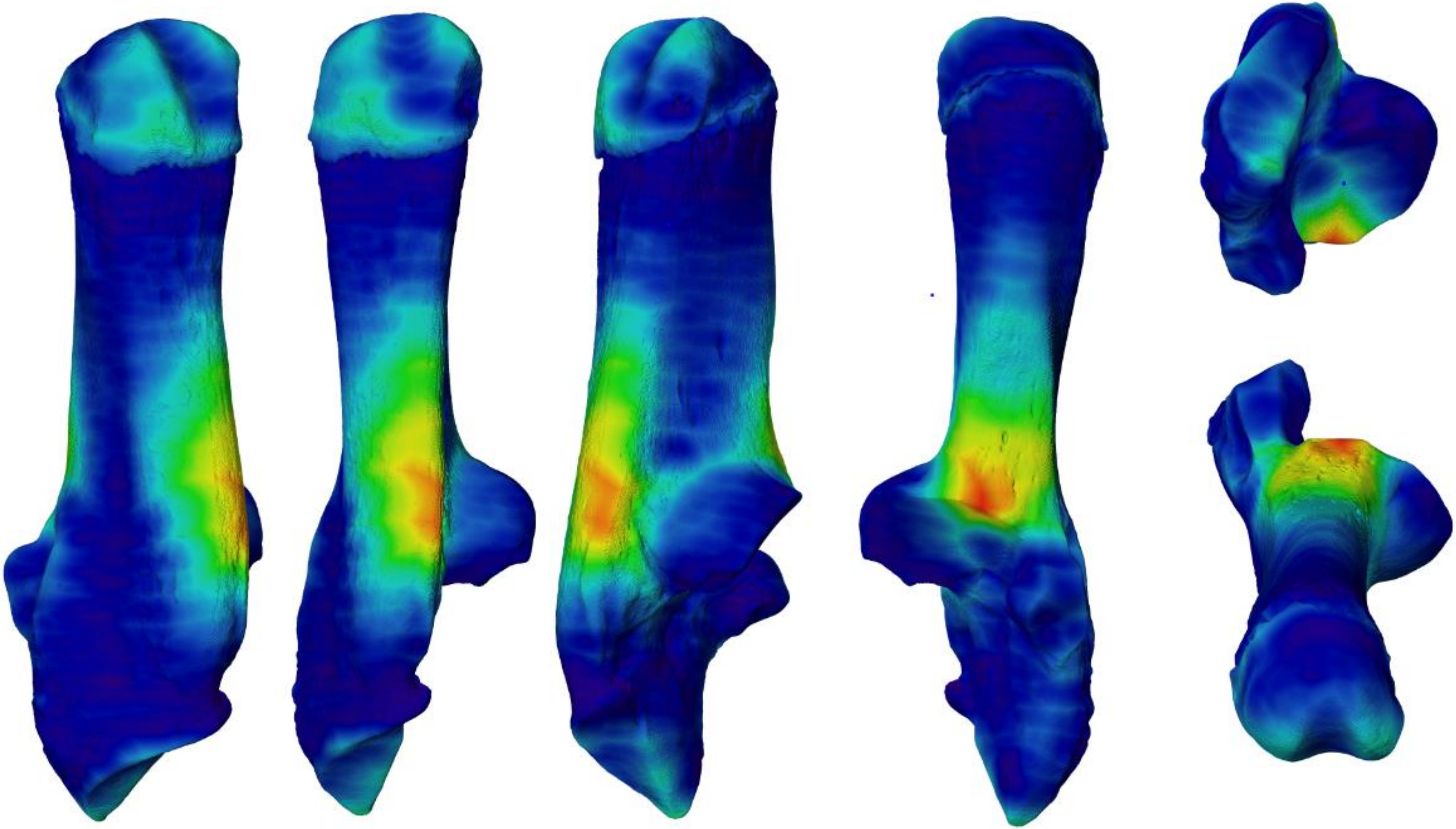

COMP3OS1

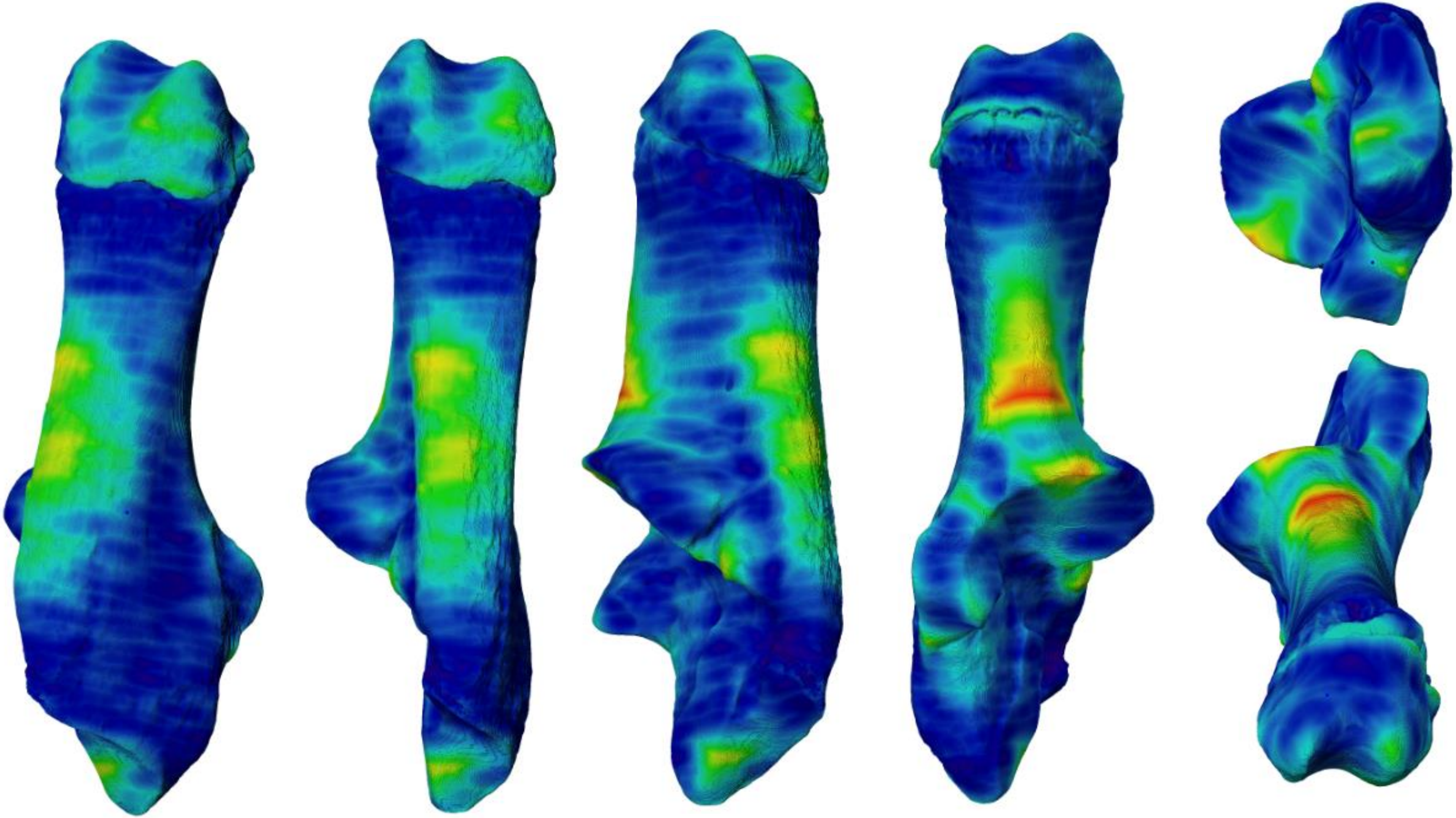

COMP3OS2

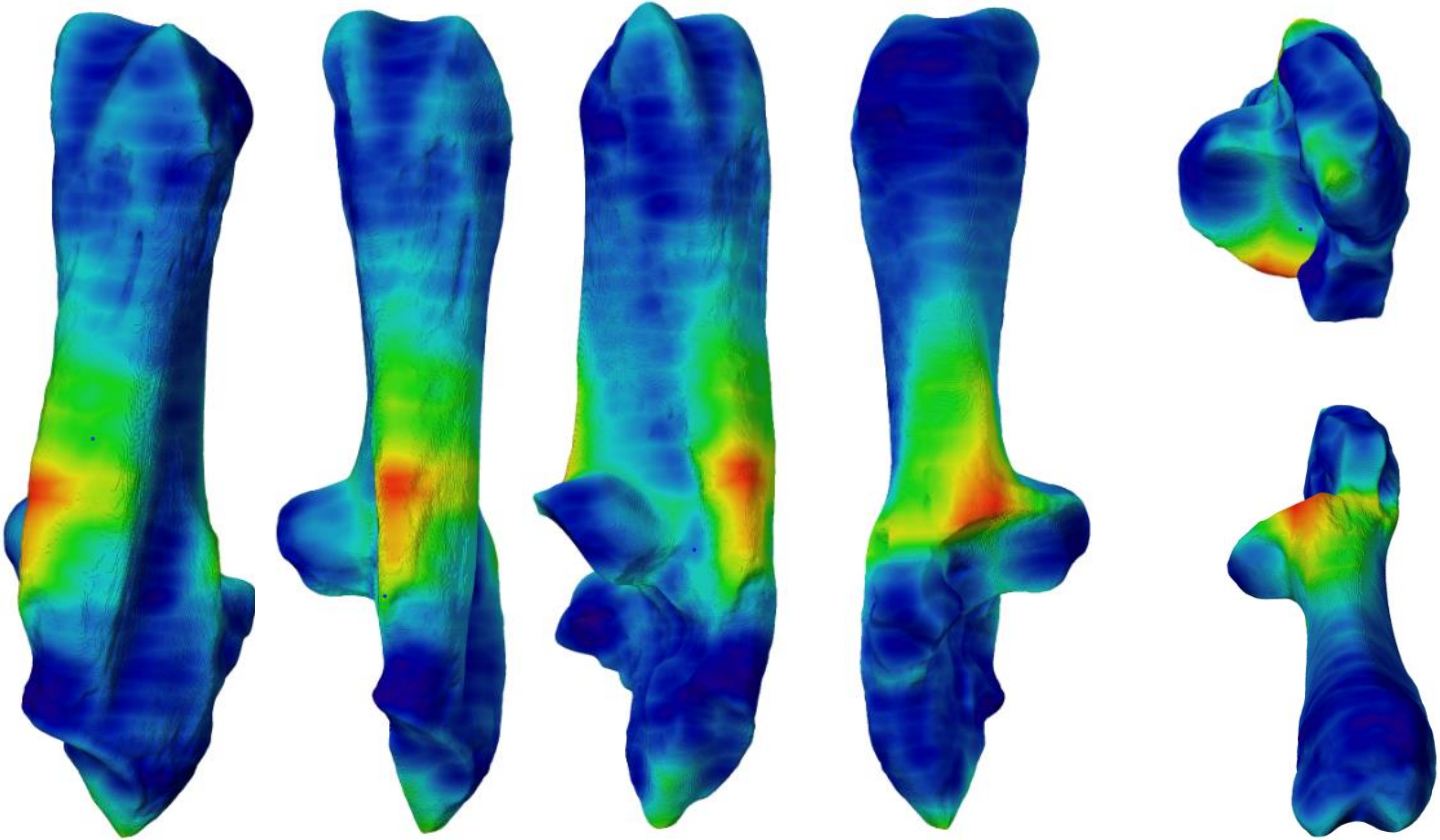

COMP3OS3

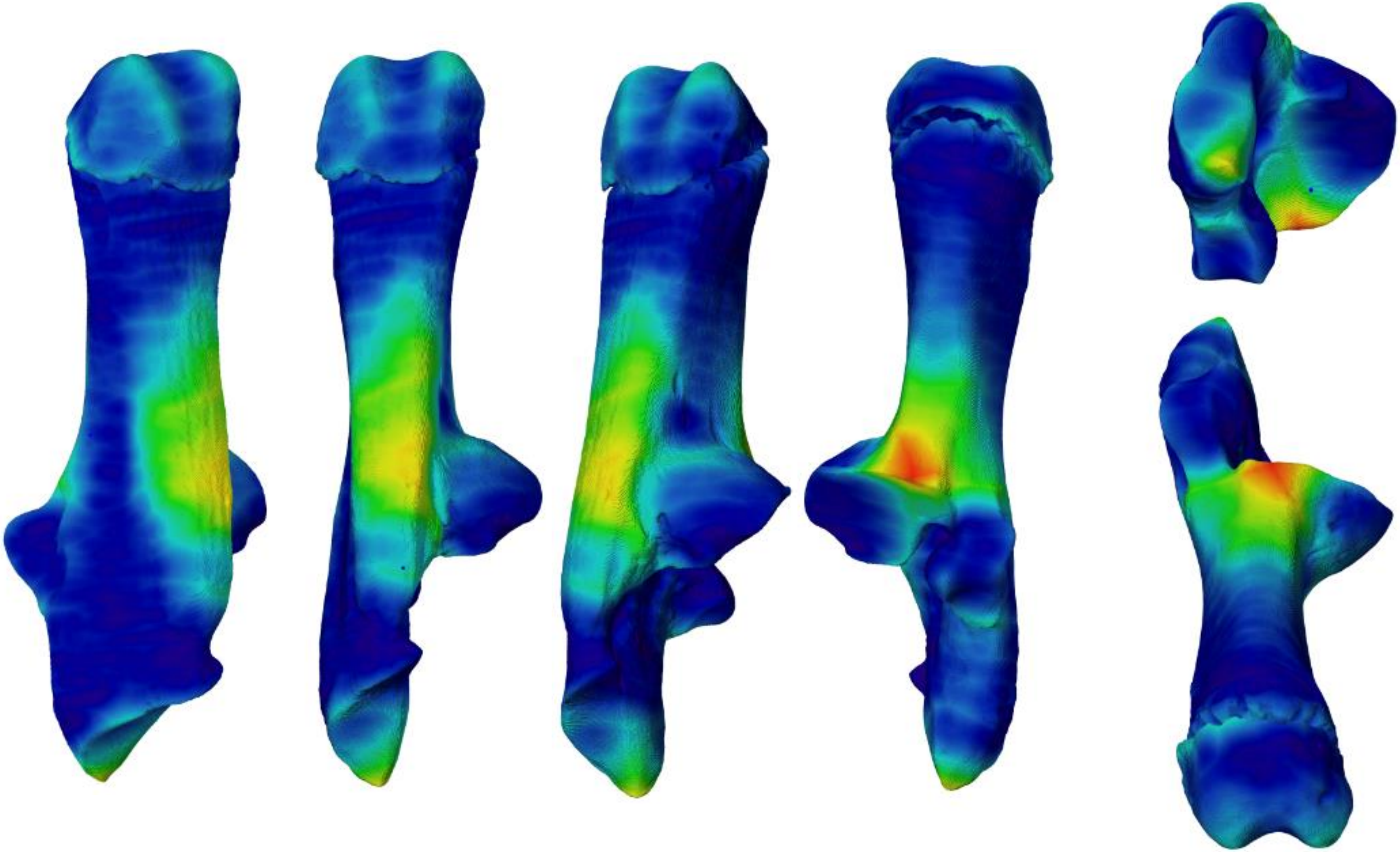

COMP3OS4

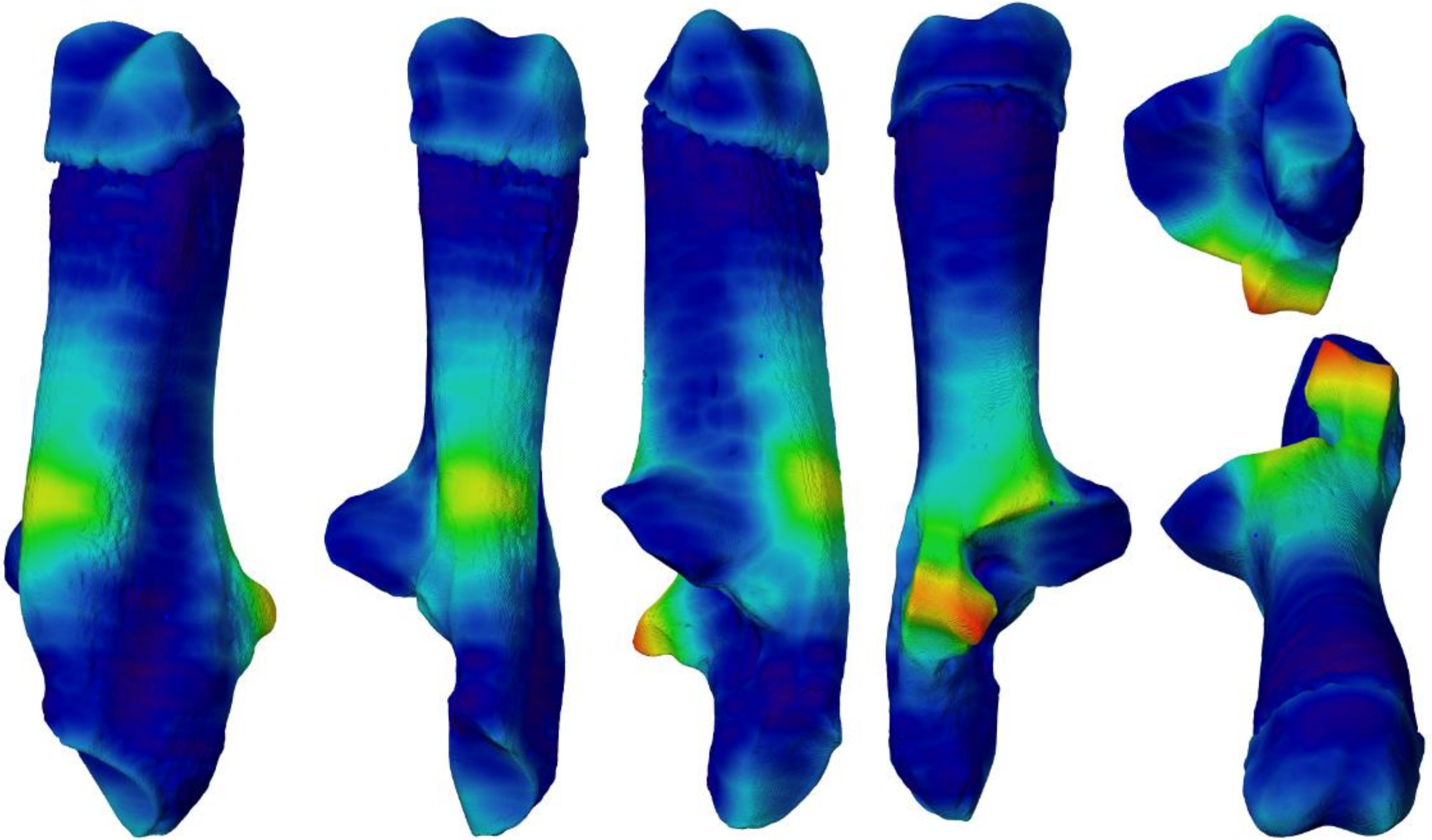

COMP5OS2

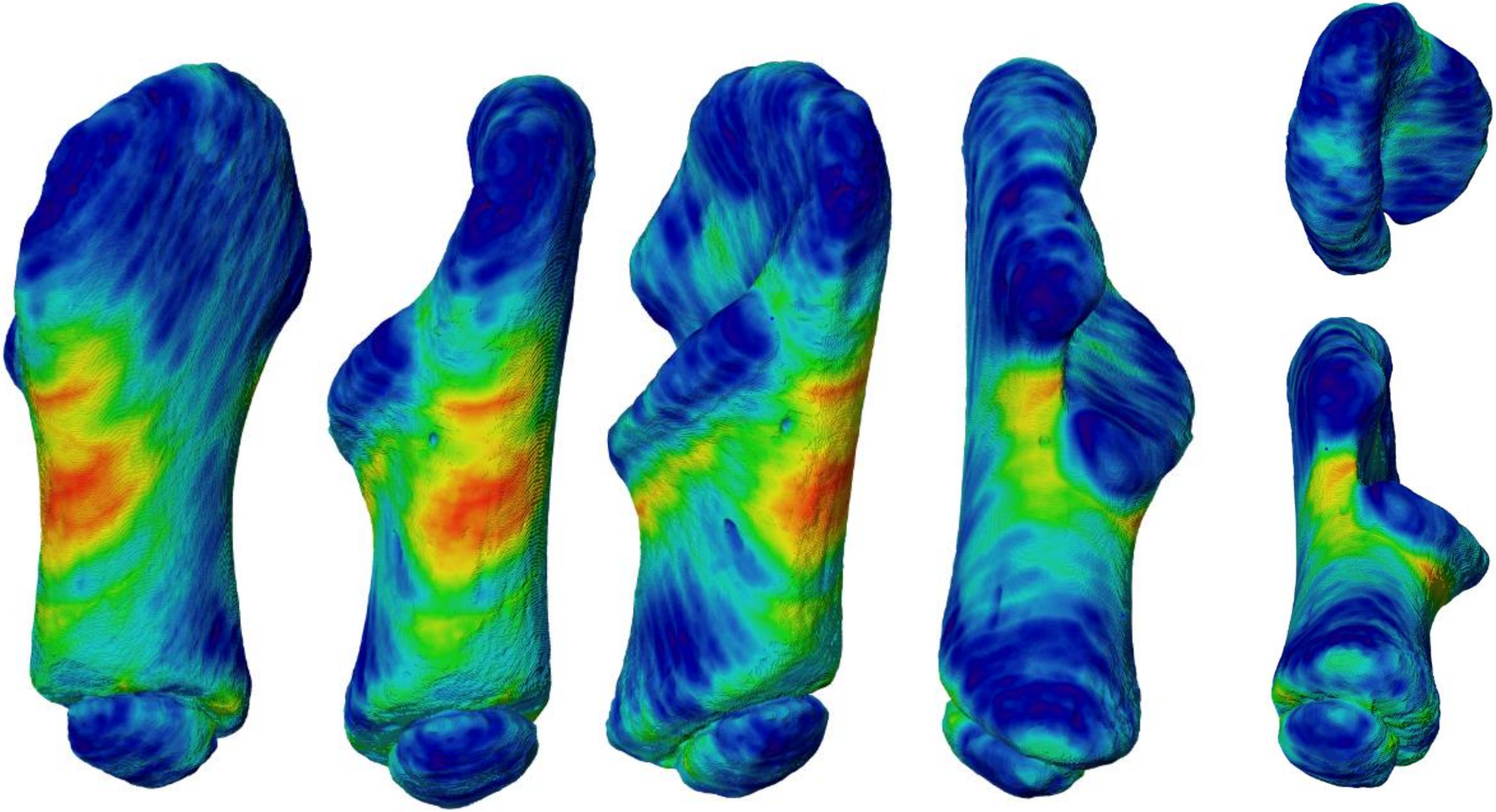

COMP5OS4

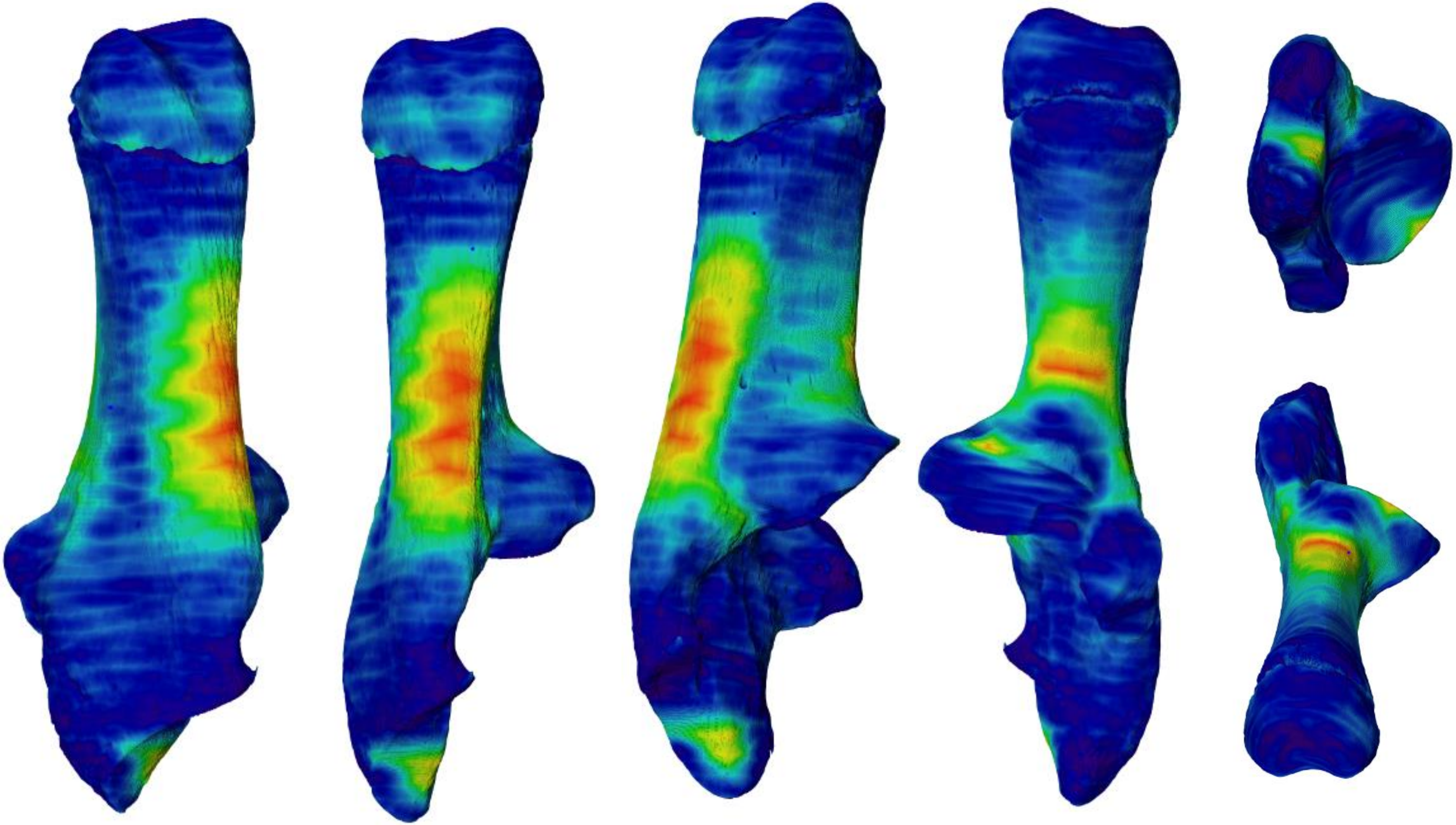

Noyenens2os2

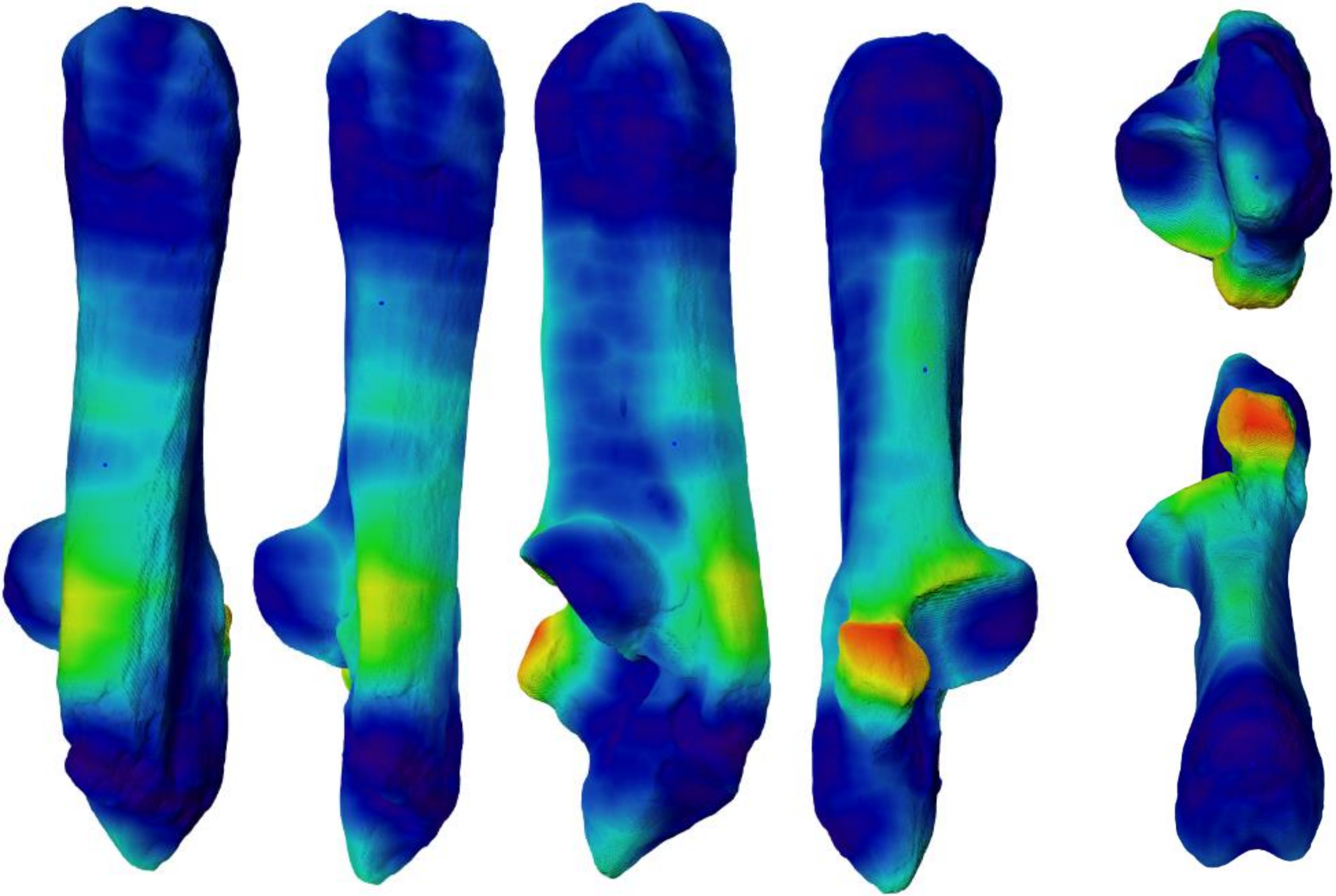

NoyenEns2os3

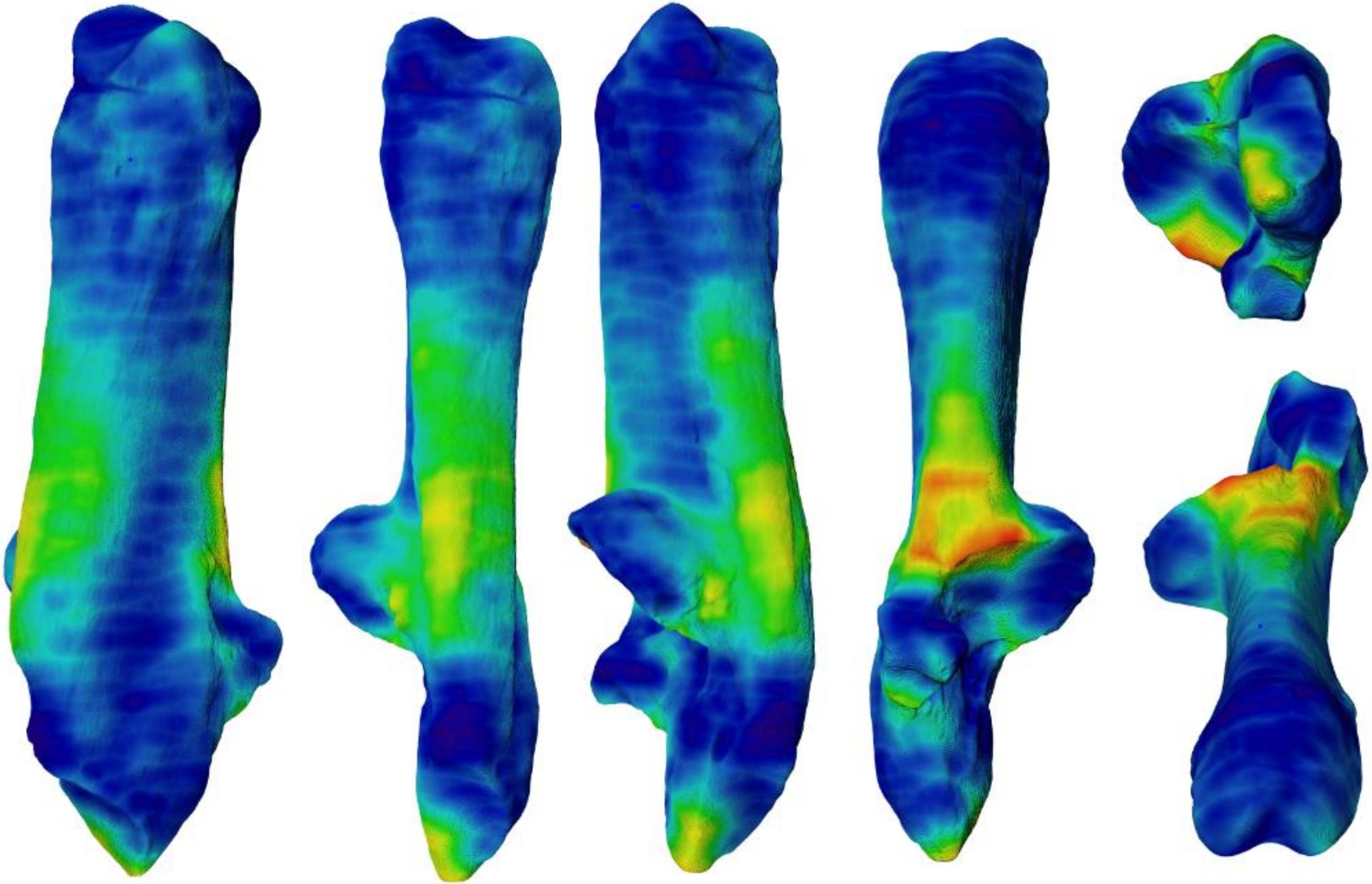

Pradat175

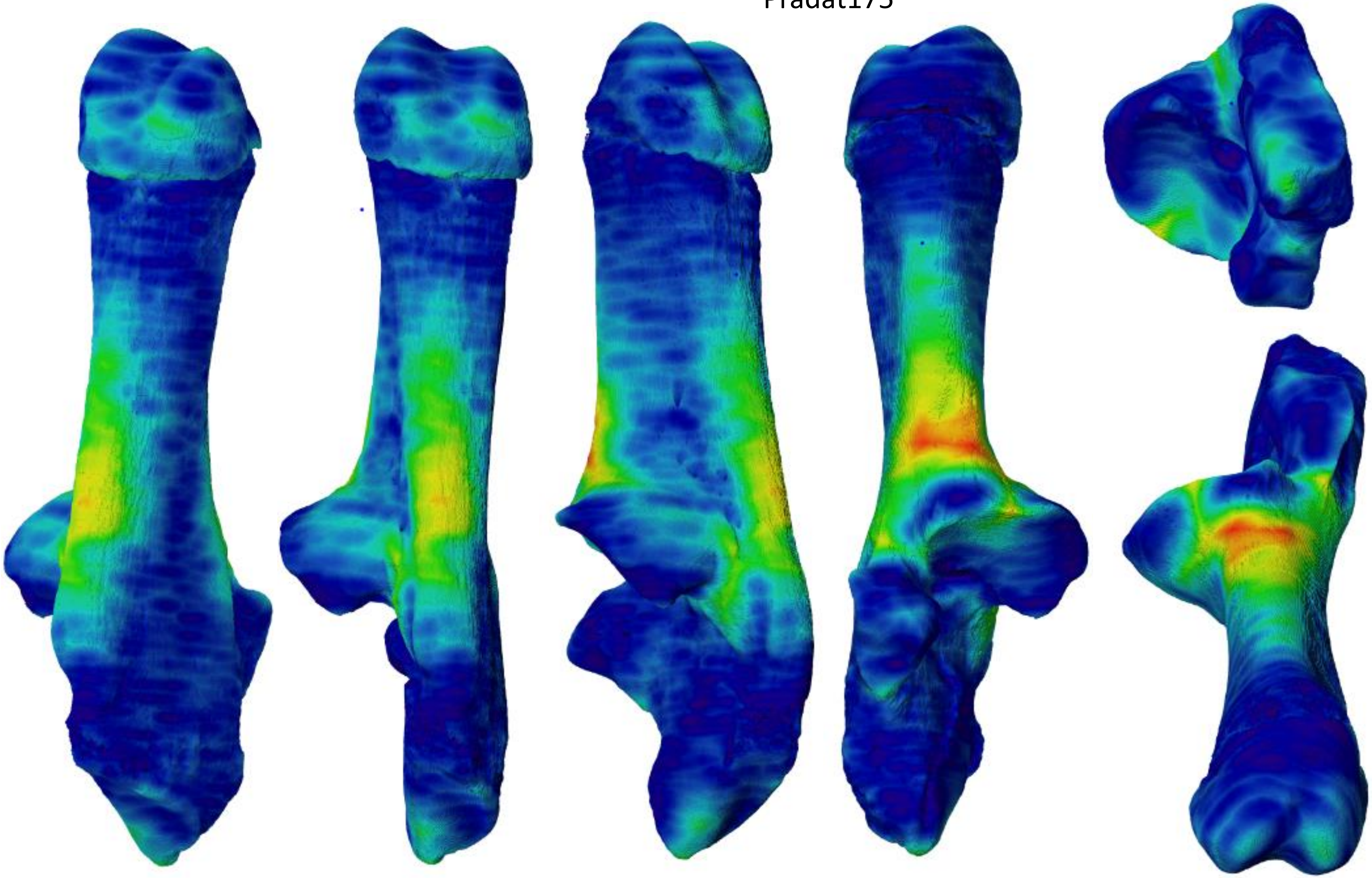

Pradat185

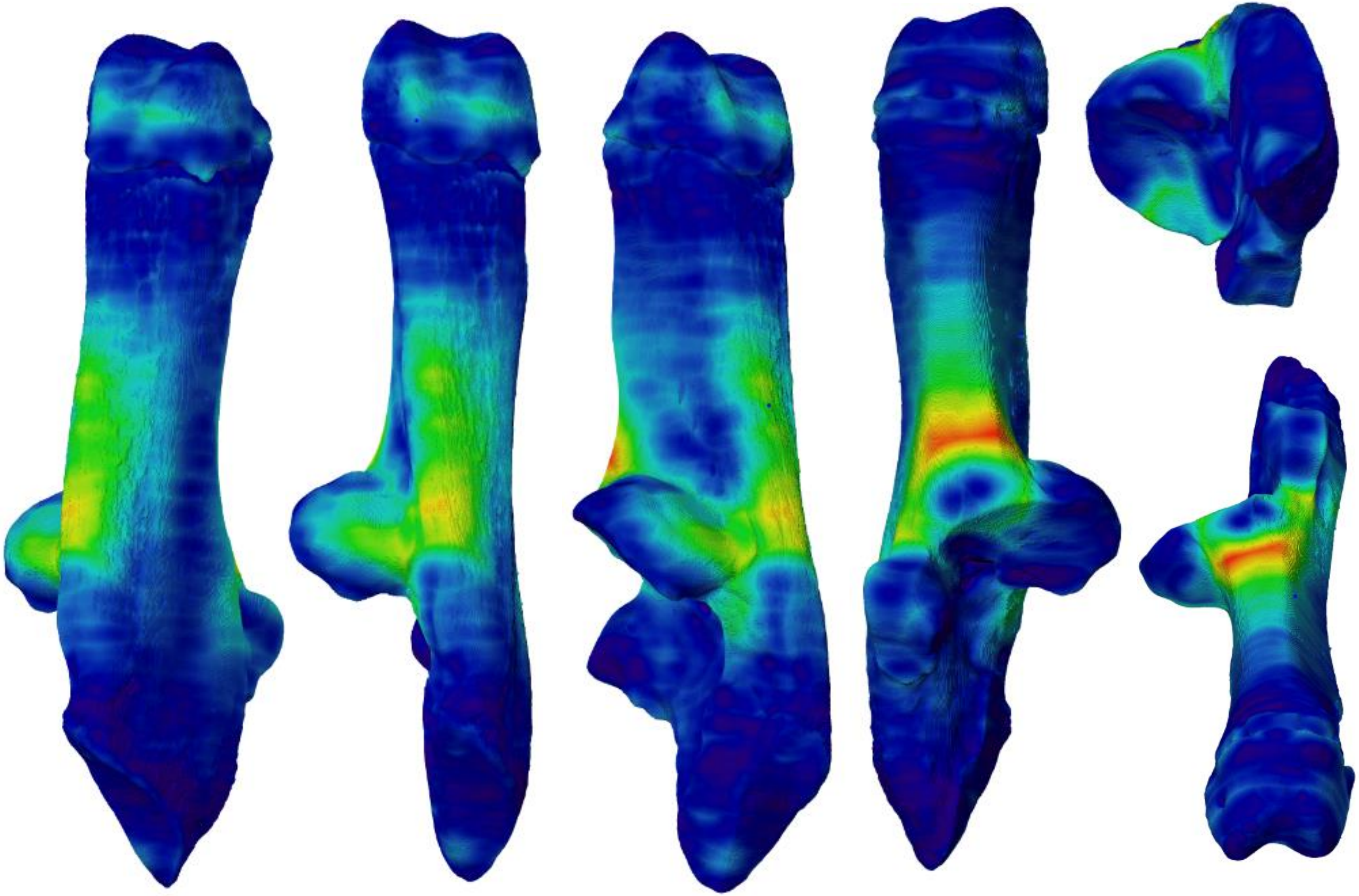

Pradat188

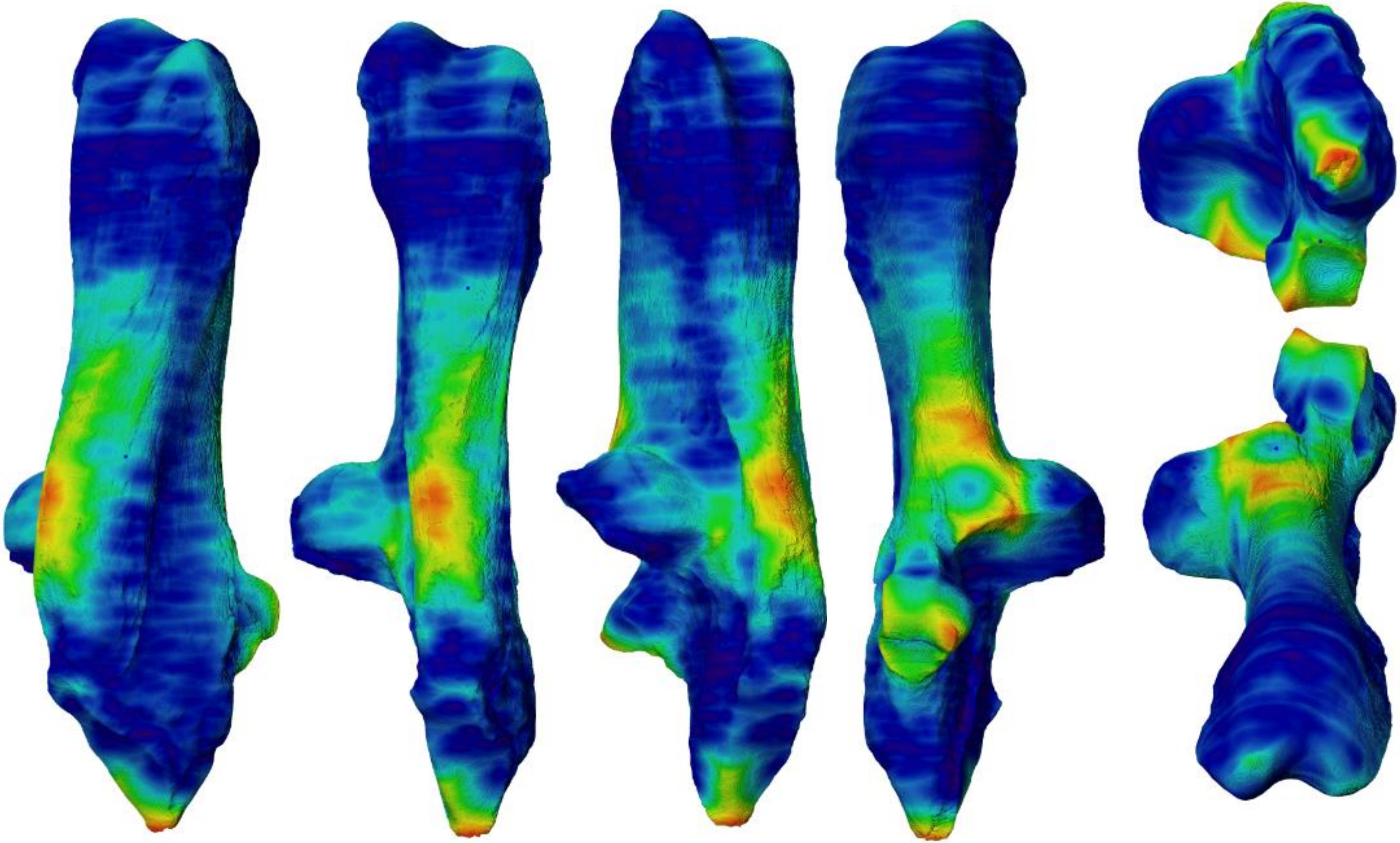
