## Supplementary material for "Can growth in captivity alter the calcaneal microanatomy of a wild ungulate?": R-script

library(ade4)

library(factoextra)

library(ggplot2)

library(tidyverse)

library(ggpubr)

library(rstatix)

\####

sangliers<-read.table("G:/Mon Drive/Travail/Sanglier nouveau/tableaux/stat/tableau final stats V5.txt",header=T,sep='\t')

summary(sangliers)

sangliers

\### creation of the PCA, distribution of the variance and choice of the number of axes

acp.sangliers<-dudi.pca(sangliers[,4:8])

\# Representing individuals and context on the PCA

#axes 1v2

fviz_pca_ind(acp.sangliers,

label = "none", # hide the labels

habillage = sangliers$Contexte, # color by factor

addEllipses = TRUE, # add ellipses

ellipse.type = "convex" # control the "type" of ellipses

)

\# to assign the colors we want, where the order of enunciation of the colors allows to associate the right group

couleurs <- c("springgreen2","turquoise2","slateblue1","snow3","limegreen")

\# here acp2 corresponds to the plot of acp1 to which we make the desired changes

acp1 <- fviz_pca_ind(acp.sangliers,

label = "none", # hide the labels

habillage = sangliers$Contexte, # color by factor

addEllipses = TRUE, # add ellipses

ellipse.type = "convex") # control the "type" of ellipses

acp1

axis <- theme(axis.line = element_line(colour = "black", size = 0.3, linetype = "solid"),

axis.text.x= element_blank(),

axis.text.y= element_text(colour = "black", size = 15),

axis.ticks= element_line(colour = "black", size = 1))

acp2 <- acp1 + scale_color_manual(values=couleurs) +

scale_fill_manual(values=couleurs) + axis

acp2

#################################################################

###boxplot

P<- ggplot(sangliers, aes(x=Contexte, y=RMaxT, fill=Contexte)) + geom_violin()

\# without tails with colors

P

P<- ggplot(sangliers, aes(x=Origine, y=RMaxT)) + geom_violin()###without the tails without colors

P

#################################################################### stats test

sangliers2<-read.table("G:/Mon Drive/Travail/Sanglier nouveau/tableaux/stat/tableau final stats V5.txt",header=T,sep='\t')

##################### MANOVA

\### check the size of the samples

sangliers2 %>%

group_by(Contexte) %>%

summarise(N = n())

\### identification of aberrant valuessangliers2 %>%

sangliers2 %>%

group_by(Contexte) %>%

identify_outliers(WBV)

\### univariate normality

sangliers2 %>%

group_by(Contexte) %>%

shapiro_test(WBV, C, X.Trab, TC, RMeanT, RMaxT) %>%

arrange(variable)

\###

sangliers2 %>%

select(WBV, C, X.Trab, TC, RMeanT, RMaxT) %>%

mshapiro_test()

\### le test (MANOVA)

res.man<-manova(cbind(WBV, C, X.Trab, TC, RMeanT, RMaxT) ~ Contexte, data = sangliers)

summary(res.man)###faire la MANOVA, --> Are the variables different depending on the context?

summary.aov(res.man)### find out which variable is significantly different according to the context?

###################################ANOVA

\### testing normality

\# Build the linear model

model <- lm(X.Trab~ Contexte, data = sangliers)

\# Create a QQ plot of residuals

ggqqplot(residuals(model))

\# Compute the Shapiro-Wilk normality test

shapiro_test(residuals(model))

\# verify the hypothesis of homogeneity of variances

plot(model, 1)

sangliers %>% levene_test(WBV ~ Contexte)

\# now we can do the ANOVA

res.aov <- sangliers%>% anova_test(WBV ~ Contexte)

res.aov

\## post hoc test

pwc <- sangliers %>% tukey_hsd(WBV ~ Contexte)

pwc

############## variable quanti vs quali : Chi2

\# Creation of vectors corresponding to the number of individuals with or without a thickening for CBd within the 5 contexts:

oui = c(10,4,5,1,4)

non = c(5,7,5,4,2)

\# Creation of a comparative matrix:

tableau = matrix(c(oui, non),2,5,byrow=T) # (2 : number of rows and 4 number of columns (contexts))

\# Performing the chi-square test - the results are saved in "khi_test"

khi_test = chisq.test(tableau)

khi_test # displays the test result

\### if the coditions of the anova not respected, test of krustal wallis

res.kruskal <- sangliers %>% kruskal_test(X.Trab ~ Contexte)

res.kruskal

###post hoc de krustal

pwc <- sangliers %>%

dunn_test(X.Trab ~ Contexte, p.adjust.method = "bonferroni")

pwc

#######################regression

plot(sangliers[,3]~sangliers[,18])### basic if you want to observe a correlation graphically.

p<-lm(sangliers[,3]~sangliers[,18])### check statistically if there is a correlation between our variables, summary(p)

r<-cor(sangliers[,3],sangliers[,18])### look at the intensity of the correlation of my different quantitative parameters between them

na.action(sangliers)

summary(p)

summary(r)
